# Evolutionary and demographic consequences of temperature-induced masculinization under climate warming: the effects of mate choice

**DOI:** 10.1101/2020.06.08.139626

**Authors:** Edina Nemesházi, Szilvia Kövér, Veronika Bókony

**Affiliations:** Lendület Evolutionary Ecology Research Group, Plant Protection Institute, Centre for Agricultural Research, Herman Ottó út 15, 1022 Budapest, Hungary; Konrad Lorenz Institute of Ethology Department of Interdisciplinary Life Sciences University of Veterinary Medicine, Savoyenstr. 1a, A-1160 Vienna, Austria; Conservation Genetics Research Group, Department of Ecology, University of Veterinary Medicine Budapest, István utca 2, 1078 Budapest, Hungary

**Keywords:** sexual selection, sex ratio selection, climate change, mate choice, sex reversal

## Abstract

**Background:** One of the dangers of global climate change to wildlife is distorting sex ratios by temperature-induced sex reversals in populations where sex determination is not exclusively genetic, potentially leading to population collapse and/or sex-determination system transformation. Here we introduce a new concept on how these outcomes may be altered by mate choice if sex-chromosome-linked phenotypic traits allow females to choose between normal and sex-reversed (genetically female) males.

**Results:** We developed a theoretical model to investigate if preference for sex-reversed males would spread and affect demographic and evolutionary processes under climate warming. We found that preference for sex-reversed males 1) more likely spread in ZW/ZZ than in XX/XY sex-determination systems, 2) in populations starting with ZW/ZZ system, it significantly hastened the transitions between different sex-determination systems and maintained more balanced adult sex ratio for longer compared to populations where all females preferred normal males; and 3) in ZZ/ZW systems with low but nonzero viability of WW individuals, a widespread preference for sex-reversed males saved the populations from early extinction.

**Conclusions:** Our results suggest that climate change may affect the evolution of mate choice, which in turn may influence the evolution of sex-determination systems, sex ratios, and thereby adaptive potential and population persistence.

## Background

As the Earth’s climate is warming, the persistence of wildlife populations is threatened by climate-driven changes in abiotic and biotic factors [1], among which the sex ratio can play a crucial role. In some species, offspring sex is determined by environmental temperatures experienced during a sensitive period of gonadal development (environmental or temperature-dependent sex determination). In such species, climate warming may increasingly distort the populations’ sex ratios, leading to loss of genetic diversity and adaptive potential via reduced effective population sizes and, ultimately, to demographic collapse [2]. However, species that have genetic sex determination, where the sex chromosomes or other genetic elements trigger male or female sexual development, are not safe from climate-induced sex-ratio shifts either. Sex reversals, where genetically female individuals become phenotypic males or *vice versa*, have been observed in various poikilothermic taxa including fish [3], amphibians [4–7], reptiles [8], and invertebrates [9]. With increasing interest in this topic, sex reversal has been demonstrated in a growing number of species, suggesting that this phenomenon may be widespread [8]. In many of these cases, sex reversal is caused by extremely low or high environmental temperatures during early individual development. Exposure to unusually high temperature can cause masculinization in some species and feminization in others, as observed among urodelans and lizards, whereas masculinization seems to be the typical response to high temperatures in anurans and fish [3, 4, 8, 10]. In natural populations of such species, global climate change may potentially distort the sex ratios and, with significant temperature rise, may lead to population extinction [11]. Furthermore, theoretical models suggest that climate change may cause, via temperature-induced sex reversals, drastic changes in the population’s genetic constitution, including novel genotypes, sex-chromosome extinction, and turnovers of the sex-determination system e.g. from genetic to temperature-dependent sex determination as well as from female-heterogametic (ZW/ZZ system; ZW females, ZZ males) to male-heterogametic (XX/XY system; XX females, XY males) sexchromosome systems [11–15].

These theoretical models of climate-driven sex reversals assumed that sex-reversed individuals are not recognized during mating, their reproductive success depending only on their fertility. However, because sex-reversed individuals can differ from normal individuals in fecundity and offspring sex ratio [16–18], individuals may benefit from taking sex reversal into account during mate choice. Choosy females then could influence the sex ratio of their offspring: while mating with a normal male would result in 50% male (XY or ZZ) and 50% female (XX or ZW) offspring, a sex-reversed (XX or ZW) male would produce 100% (XX) or 75% (ZW and WW) female offspring. Such mate-choice decisions may potentially alter the outcomes of climate change, as females might adjust their offspring sex ratios to the population sex ratio distorted by temperature-induced sex reversals and, ultimately, might save the population from extinction. To our knowledge, this idea has never been addressed by theoretical studies, despite the possibility that sex-reversed individuals may be recognized by conspecifics on the basis of phenotypic traits linked to sex chromosomes. For example, there are several taxa with sex-chromosome-linked (hereafter referred to as sex-linked) body colour genes (SI1 Table S1) [19–21]. In such species, females may recognize sex-reversed (genetically female) males by the absence of a Y-linked male colour trait (in XX/XY system) or by the presence of a W-linked female colour trait (in ZW/ZZ system). Many other sexually selected traits, such as pheromones and song, may also often be sex-linked (reviewed by Kirkpatrick and Hall [22]).

In this study we developed an individual-based theoretical model to investigate the role of female choice in the evolutionary and demographic consequences of temperature-induced sex reversal in a warming climate. We focused on masculinization because this seems to be the more frequent response to high developmental temperatures across ectothermic vertebrates [3, 4, 8, 10]. We assumed that females can distinguish between normal and sex-reversed males by sex-linked traits, and some females tend to mate with sex-reversed males due to a genetically encoded preference. We examined if, and under what conditions, female preference for sex-reversed males spreads in a population while such males are becoming more common due to warming climatic conditions, and we investigated how this preference influences the adult sex ratio, the evolutionary changes in the sex-determination system, and the duration of population persistence. Although populations can no longer persist once climate has become so hot that all females get masculinized, sexual selection may cause earlier extinction if it biases the adult sex ratio toward males. Alternatively, if sexual selection leads to less male-biased sex ratios, it may protect the population from demographic and environmental stochasticity and thereby from premature extinction. We investigated scenarios of long-term climate warming (which occurred several times over Earth’s history [23, 24], because models of contemporary climate change project continuous warming for the twenty-first century despite the recent hiatus [25, 26].

## Methods

### The model

We followed the approach of recent theoretical models on the evolution of sex determination, where sex has been assumed to be a threshold trait: a phenotypically discrete trait (i.e. male or female) determined by individual threshold sensitivity for the endogenous level of a non-discrete factor referred to as ‘male signal’ [12–14]. In our model, individuals are diploid and carry a pair of sex chromosomes and two independently inherited autosomal loci (SI1 Table S2). Sex chromosomes are denoted by A and a, corresponding to Z and W in a ZW/ZZ system and to Y and X in an XX/XY system, respectively. Each sex chromosome harbours a sex-determinant locus that encodes male signal: A causes production of the male signal at 1.5 times higher level (*sig_A_*) than a does (*sig_a_*) under the same environmental conditions (for graphical explanation, see SI1 Fig. S1). Male signal expression increases with environmental temperature, such that the individual level of male signal expression (*sig_indiv_*) in an *Aa* individual is calculated with equation 1:

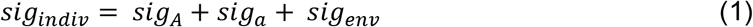

where *sig_env_* is the exogenous level of male signal due to environmental temperatures (see below). All genotypes have the same temperature sensitivity in male signal production (i.e. tparalell reaction norms). We followed Grossen and colleagues [13] in assuming that male signal increases monotonically with temperature, based on the empirical observations of masculinising effects of high temperatures and feminising effects of low temperatures in several amphibians and fish [4, 27], although we note that counter-examples exist and nonlinear temperature reaction norms are also possible [14, 15].

The autosomal locus *thr* encodes the individual threshold for *sig_indiv_* that needs to be exceeded in order to develop male reproductive organs (otherwise, the individual becomes female). We calculated the individual’s threshold value as the sum of the values of the two alleles that the individual carried at the *thr* locus. Although sex determination in temperature-sensitive systems is often assumed to have a polygenic basis [28, 29], there is very little empirical evidence either *pro* or *contra* [30–32]; and the findings of the seminal models of temperature-sensitive sex determination were not sensitive to the assumption of one versus more loci [28]. Therefore, for simplicity we assume that only two *thr* alleles at a single locus are present in the population (however, we explored additional simulations with 10 *thr* alleles; SI2 Fig. S5). Allele values on locus *thr* are set according to the sex-determination system operating in the initial population (see SI1 Fig. S1 and SI1 Table S2): threshold value of homozygotes for the *thr_low_* allele is above the average male signal level of the normal female genotype (*Aa* in system ZW/ZZ and *aa* in system XX/XY) but just below its maximum signal level realized at the temperature range before climate warming, following Schwanz and colleagues [14]. Value of the *thr_high_* allele is set so threshold in *thr_high_* homozygotes equals the average of male signal levels that are determined by the normal female and male genotypes (*Aa* and *AA*, or *aa* and *Aa*, respectively, in ZW/ZZ and XX/XY system) and realized under the temperature variation before climate warming. All simulations start with both *thr* alleles present in the population at 0.5 frequency.

We ran simulations where the initial sex-determination system was either XX/XY or ZW/ZZ. We assumed that sex-reversed males (XX, ZW or WW) are as viable and fecund as normal males, following previous models and empirical data [27, 33, 34]. Note that our previous model predicted that 25% decrease in reproductive success of masculinized individuals had little effect on adult sex ratios and sex chromosome frequencies, whereas their sterility lead to the ZW system behaving exactly like the XY system [11]. Further, we assumed that the WW genotype (*aa* in ZW/ZZ system) is phenotypically equivalent to normal females (i.e. has the same viability, fecundity, and ability to masculinize). Empirical data support that WW individuals can be viable [10, 17, 27, 35, 36], able to reproduce [10, 36, 37] and can also be able to develop into functional males [33, 35]. Viability and fertility of sex-reversed males and WW individuals is likely in ectothermic vertebrates, because in these taxa the sex chromosomes are usually homomorphic [38]. However, in some species the WW genotype is lethal [36], so we explored the effects of reduced WW viability in additional simulations (SI2 Fig. S2-9). We did not allow new mutations to occur on the *thr* locus or the sex-determinant locus, because in the realistically small simulated populations (starting with 200 adults) appearance of a new mutant allele is unlikely over the relatively short evolutionary time simulated here due to rapid climate warming. Note, however, that evolution is still possible due to standing variation in temperature sensitivity [15], which was allowed in our simulations by changes in relative frequencies of the *thr* alleles.

In our model, climate is warming linearly over time, each year increasing the average temperature that the population is exposed to during the breeding season. In each year, each individual may experience a different environmental temperature during the sensitive period of its ontogeny due to spatiotemporal variation in microclimatic conditions. This variation in temperature between and within years is incorporated into our model by a set of parameters defining the exogenous levels of male signal (SI1 Table S2), such that in year *t* the *sig_env_* level for each individual is calculated as described in equation 2.

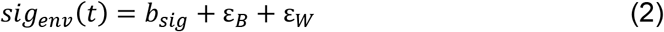

where *b_sig_* is the slope of the yearly increase in mean *sig_env_* levels in the population, ε_*B*_ is the normally distributed error of the yearly mean *sig_env_* levels (between-year climatic variance), and ε_*W*_ is the normally distributed error of individual developmental temperatures (within-year climatic variance; see SI1 Fig. S1). We did not vary the values of *b_sig_*, ε_*B*_, and ε_*W*_ in our simulations, but fixed each at a likely value based on empirical data. We set *b_sig_* = 0.003 (0.3%) assuming no masculinization before 1970 (the start of contemporary, human-induced climate warming) and an increase to 9% masculinization by 2000, based on the first report of sex reversal in a natural amphibian population [5]. We set ε_*B*_ = 0.01 based on the standard deviation of temperature anomalies observed in the Northern Hemisphere between 1970 and 2000 [39]. We set ε_*W*_ = 0.05, a value five times higher than ε_*B*_, which is realistic based on empirical data for reptiles [15]. These settings ensure that, before the start of climate warming, a stable genetic sex-determination system (either XX/XY or ZW/ZZ) persists, with only occasional events of sex reversal in individuals experiencing unusually high temperature (resulting in ca. 0.2% masculinization rate before the start of climate warming; see SI1 Fig. S1 for visual representation). We allowed this stable state to persist for 50 years (i.e. burn-in period, with *b_sig_* = 0), after which we simulated climate warming by increasing the mean value of *sig_env_* each year by *b_sig_*. We chose this relatively rapid, linear increase in masculinization rate to model the climate warming observed in the recent past and expected in the near future [39]. Note that our model does not include temperature *per se*, only its effects on male signal production.

Besides the sex-determinant male signal locus, each sex chromosome harbours another locus that encodes a sexually dimorphic phenotypic trait, which for simplicity we will refer to as colour and model it as a binary trait. We assume that females can recognize normal males by a “male colour” expressed in the presence of chromosome Y (*A*) when we start with an XX/XY sex determination system, and by the absence of a “female colour” encoded on chromosome W (*a*) when the initial system is ZW/ZZ. The autosomal locus *C* determines preference in females for mating partners based on sex-linked colour, and is not expressed in males. Values of *C* alleles determine the probability that a female would choose a normal male if both normal and sex-reversed males were equally available. We allowed a maximum of two *C* alleles in the population: allele *C_N_* encoding a strong but non-exclusive preference (0.9) for normal males, and allele *C_R_* encoding the same extent of preference for sex-reversed males (i.e. 0.1 probability of choosing a normal male; see SI1 Table S2). In our simulations, inheritance of *C* is fully dominant/recessive, where heterozygotes show the same choosiness as homozygotes for the dominant allele do. Starting with either XX/XY or ZW/ZZ sex-determination system, we investigated three different sexual-selection scenarios. In scenario 0% *C_R_*, no preference allele for sex-reversed males exists (all females are *C_N_C_N_* homozygotes). In scenario 10% *C_R_*, the *C_R_* allele is dominant and rare (relative frequency 0.1 in the starting population), thus initially only 19% of females (genotypes *C_R_C_R_* and *C_R_C_N_*) express preference for sex-reversed males. This scenario could be realistic if, for example, males with female-like coloration occurred somewhat regularly in the near past (e.g. due to randomly occurring sex-reversals [6, 40]; see SI1 Table S2), maintaining a certain level of variance in female mating preferences. In scenario 90% *C_R_*, the *C_R_* allele is recessive and widespread (relative frequency 0.9), thus initially 81% of females (*C_R_C_R_*) express preference for sex-reversed males. This latter scenario is possible if allele *C_R_* has spread in the initial population by neutral processes or due to a sensory bias for the “female colour” that was generally not expressed in males while sex reversal was very rare(SI1 Table S2) [41]. This can be seen as a similar case to the famous experiment of Basolo [42] where 93% of females of the swordless fish *Priapella olmeacae* preferred males with artificial swords. For simplicity, we only investigated the effects of relatively low and high *C_R_* frequency, assuming that these two scenarios represent the range of potential effects (i.e. the effects of an initial *C_R_* frequency between 10% and 90% may fall between the effects presented here). We chose to set the initially rare *C* allele to dominant because rare recessive alleles are easily lost by drift. However, we additionally examined several other scenarios, including intermediate inheritance and a *C* allele encoding lack of preference (indiscriminate mating), to assess the sensitivity of our results to these settings (SI2 Fig. S2-9).

We aimed to build a relatively realistic model where a number of parameters affect the life history and demography of iteroparous animals including the age of maturation, annual survival rates differing across life stages, fertility, environmental carrying capacity, and limited number of mating events for each individual per breeding season. We set these parameters to be representative for amphibians using empirical data from the literature, mostly following our previous model [11], although similar parameter settings may be representative for other temperature-sensitive taxa such as fish or reptiles. Every year, the population produces *N* offspring calculated using equation 3:

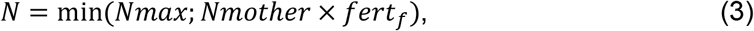

where *Nmax* is the carrying capacity, *Nmother* is the number of adult females that found a mating partner, and *fert_f_* is the average number of offspring each female can recruit in the absence of density-dependence (see SI1 Table S2). For simplicity, we assume that the annual survival rate of juveniles and adults, age of maturity, and maximum life span were independent of both genotypic and phenotypic sex, except for scenarios with reduced WW viability (SI1 Table S2). Each year, adults participate in a single breeding event, during which mate choice is constrained by the availability of mating partners: within a breeding season, each female can mate with one male, while each male can mate with maximum three females (“libido”; see SI1 Table S2). Females in a randomized order, one after another, choose a single mate from the pool of still available phenotypic males according to the females’ preference and relative frequency of the available male genotypes, resulting in altogether *Nmother* parent pairs. Thus, in an XX/XY system for example, a female with a dominant *C_R_* allele would mate with an XY male with a probability of *C_R_* × P_XY_, where *C_R_* is the strength of preference for normal males and P_XY_ is the proportion of mating opportunities with XY males out of all available matings (i.e. number of available males multiplied by their remaining “libido”). Parent pair of each of the *N* offspring is chosen randomly, and each offspring randomly inherits one sex chromosome and one *thr* and *C_R_* allele from each parent, and receives a *sig_env_* value, based on which its phenotypic sex is determined. For settings of all parameter values in detail and their justification see SI1 Table S2. Our model was developed in R programming language, and the simulations were run in R 3.4.3 [43]. The source code is available in SI3.

### Statistical analyses

We ran 100 simulations for each scenario (specific R code for these simulations is available in SI4). Changes over time in the relative frequency of genotypes, allele frequencies and sex ratios in each simulation are shown in SI5 Fig. S10-16. We compared the dynamics of sexdetermination system evolution, sex ratios, and population persistence among scenarios by calculating the following values from each run. First, to identify transitions between different sex-determination systems, we assumed that in an XX/XY system (i.e. *aa* females, *Aa* males) at least 95% of the adult phenotypic males have the *Aa* genotype, while less than 5% are sex-reversed (*aa*), whereas a ZW/ZZ system (i.e. *Aa* females, *AA* males) persists as long as less than 5% of the adult males carry chromosome *a*. When more than 5% of phenotypic males are sex-reversed individuals, we assumed that a mixed sex-determination system is operating. Due to the stochastic nature of our model, there was a notable annual fluctuation in genotype frequencies among adult males. Therefore, we defined the start and endpoint of each sexdetermination period as the first year after which the above conditions prevailed for at least 50 consecutive years. In practice this meant that the proportion of a given genotype would never exceed or fall below the specified value again, as masculinization rate of each genotype kept increasing in the continuously warming environment. For each simulation, we calculated the length of each sex-determination period in years, and also the year of extinction which occurred when no females were left in any age groups in the population.

In order to investigate changes in adult sex ratio (ASR, the proportion of phenotypic males among adults) over evolutionary time, we identified the sex-determination periods in which ASR deviated from 0.5 by visually inspecting our results, and for this period in each run, we calculated the average of yearly ASR values and the first year when average ASR across 5 consecutive years increased above 0.6.

For each variable above (genotype frequencies, ASR, and the length of each sexdetermination system), we compared the three scenarios (0% *C_R_*, 10% *C_R_* and 90% *C_R_*) pairwise, starting the simulations either from ZW/ZZ or XX/XY system. To this end, we entered the values calculated from each run as dependent variable in a linear model, using scenario as fixed factor, and we calculated three linear contrasts (post-hoc tests for the pairwise differences among the three scenarios) using the function ‘lsmeans’ in R package lsmeans [44]. All p-values of all these linear contrasts were adjusted simultaneously by Bonferroni correction (‘bonferroni’ method in the R function ‘p.adjust’). We chose this strict correction method to be conservative about statistical significance in our analyses. For scenarios in the sensitivity analysis where the distribution of data did not meet the requirements of linear models, we used pairwise median tests with Bonferroni correction.

For scenarios 10% *C_R_* and 90% *C_R_* we evaluated if the relative frequency of allele *C_R_* changed during the period when co-occurrence of normal and sex-reversed males allowed females to choose between them (i.e. when both normal and sex-reversed males were available with >5% frequency among phenotypic males) and the strength of sexual selection was not exceeded by the effect of drift (due to reduced effective population size, see below) on allele frequencies. For each run, we calculated the average relative frequency of allele *C_R_* among adults across 5 years at the start and end of this period, and compared the start and end values with a paired t-test (R function ‘t.test’) across all simulations within each scenario. The four p-values from these t-tests were adjusted simultaneously by Bonferroni correction.

To better understand the forces behind the changes of *C_R_* frequency in the population, we recorded for each year in each run the relative frequency of allele *C_R_* among adult males and females separately and also among the offspring’s preference alleles inherited from fathers and mothers separately. Also, we calculated the following values as detailed in SI1. For each year in each run we calculated the effective population size (N_e_), the selection coefficient (*s*) of *C_R_* resulting from sex-ratio selection, and linkage disequilibria (LD) between allele *C_R_* and 1) chromosome *A* and 2) the *thr_low_* allele. For each run, we recorded the year when the ultimate population decline started, i.e. when N_e_ permanently decreased below the average N_e_ of the 10 years before the start of climate warming when the population was in a stable state (SI1). Because our populations had overlapping generations, we estimated generation time (T; mean age of reproduction) as detailed in SI1. In all scenarios, generation time was about 3 years in our simulations.

## Results

### Consequences of climate warming

When we assumed that all females preferred normal males (scenario 0% *C_R_*), our model showed that a continuous rise in environmental temperatures caused changes in population structure in terms of genotypes and phenotypic sexes, leading to evolutionary switches between sex-determination systems and, ultimately, to population extinction (Fig. 1a,d). When starting with an XX/XY system, the increasing frequency of masculinized individuals quickly skewed the ASR towards phenotypic males and resulted in a mixed sex-determination system (hereafter referred to as ‘final period’) in which phenotypic males with both *Aa* (XY) and *aa* (XX) genotypes were present (Fig. 1a). Because individuals possessing one or two copies of the *thr_high_* allele were less susceptible to temperature-induced masculinization, and thus were able to remain females during the earliest stages of climate warming, this allele spread and usually became fixed in the population (Fig. 1a, SI5 Fig. S14a). However, even homozygotes for the *thr_high_* allele started to masculinize as climate got hotter, leading to population extinction after ca. 42 generations (Fig. 1a).

**Figure 1.**
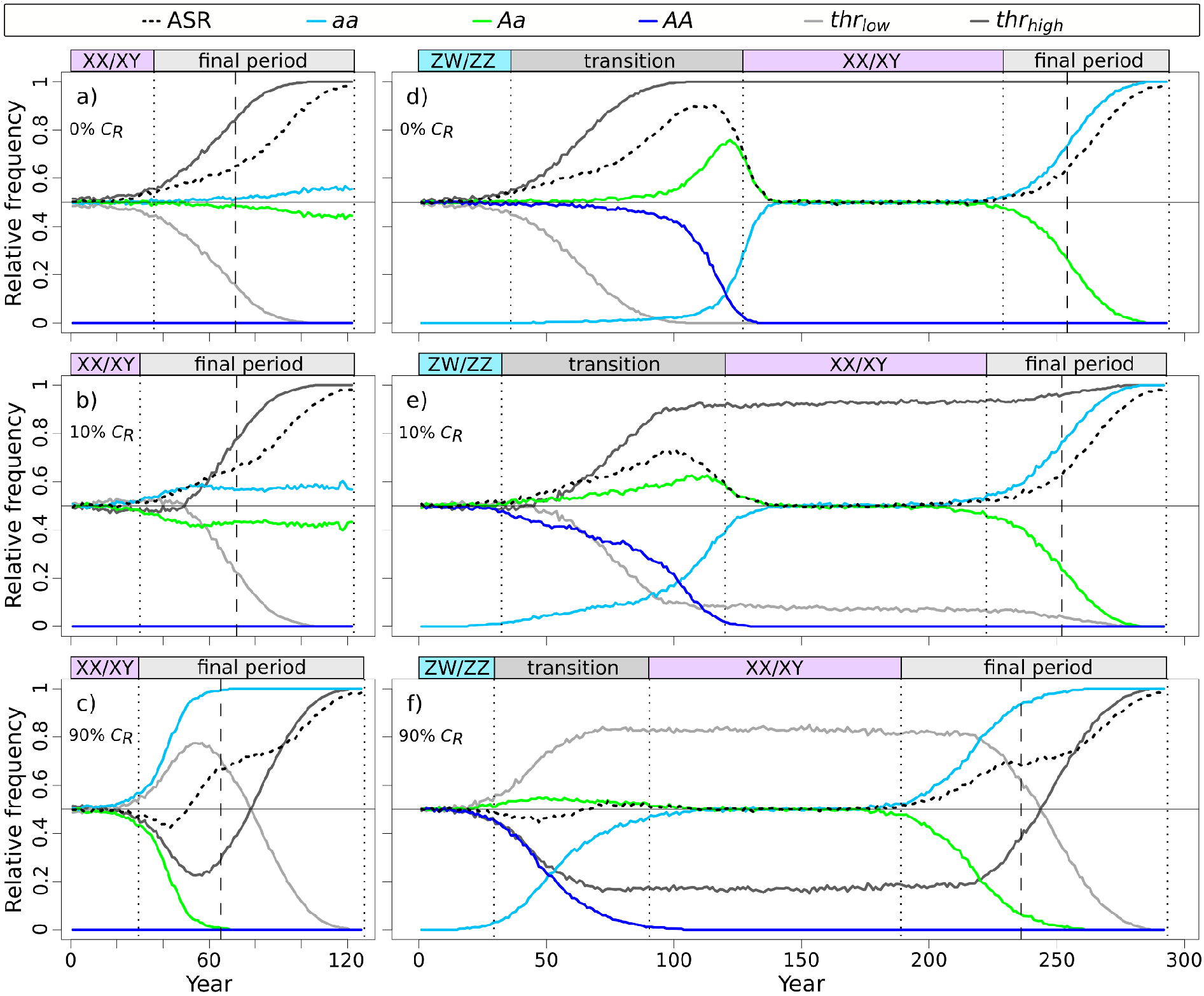
Changes in relative frequency of males (ASR), sex-chromosome genotypes and threshold alleles among adults. On the left: simulations starting with XX/XY system with scenarios 0% *C_R_* (a), 10% *C_R_* (b) and 90% *C_R_* (c). On the right: simulations starting with ZW/ZZ system with scenarios 0% *C_R_* (d), 10% *C_R_* (e) and 90% *C_R_* (f). Each curve indicates median values calculated from 100 simulations (see SI5 Fig. S10-14 for graphical results of all simulations). Vertical dotted lines indicate the end of each sex-determination period and dashed lines indicate the start of ultimate population decline. Generation time was 3 years. Year zero refers to the start of climate warming.

When starting with a ZW/ZZ system, the earliest stages of climate warming caused similar increases in ASR and frequency of the *thr_high_* allele as in the XX/XY system (Fig. 1d). However, the ZW/ZZ system adapted to climate change by transitioning into an XX/XY system as follows. Increasing frequency of masculinization led to a mixed sex-determination system in which phenotypic males with both *AA* (ZZ) and *Aa* (ZW, corresponding to XY) genotypes were present (Fig. 1d). During this transitional mixed sex-determination period, the ASR became strongly male-biased, which decreased the effective population size such that in some runs the population barely escaped extinction (SI5 Fig. S16d). Under male-biased ASR, sex ratio selection favoured the reproduction of non-preferred *Aa* males (masculinized individuals) because those produced less male-biased offspring for two reasons. First, mating between two *Aa* individuals produced fewer genotypic males (25% *AA* offspring, in contrast to 50% from normal matings). Second, such mating events produced 25% *aa* (WW) offspring, a novel genotype in the ZW/ZZ system which was resistant to temperature-induced masculinization as long as climate warming was mild, due to its genetically low levels of male signal. These “resistant females” rapidly accumulated because increasingly high numbers of sex-reversed males could reproduce: as the proportion of masculinized individuals increased among phenotypic males, more and more females were forced to accept them. Because all females preferred normal *AA* males, and the female phenotype became more and more restricted to the *aa* genotype, dis-assortative mating occurred between the two homozygote genotypes and produced an excess of *Aa* genotypes, while genotype *AA* disappeared from the population because *aa* females could not produce *AA* offspring. This way, the population transitioned to an XX/XY system, where all phenotypic females had the *aa* genotype (WW became XX) and all phenotypic males had the *Aa* genotype (ZW became XY), producing 0.5 progeny sex ratios and returning the ASR to 0.5 as well (Fig. 1d). This system persisted until *aa* individuals started to get masculinized by the ever-increasing temperatures, once again skewing the ASR towards phenotypic males. During this ‘final period’, the frequency of mating between masculinized and normal *aa* individuals increased, thereby *A* (Y) became rare or even extinct, and when no phenotypic females were left the population died out, ca. 98 generations after the start of climate warming (Fig. 1d).

### Effects of preference for sex-reversed males

When we allowed females to vary in mating preference, the presence of allele *C_R_* significantly changed the temporal dynamics of the above processes and the magnitude of ASR skew (Fig. 1–2, SI1 Table S3). The evolutionary switches between sex-determination systems happened faster: the initial period of both starting systems and the transition period between ZW/ZZ and XX/XY were shorter (but not the XX/XY system that evolved from the ZW/ZZ system), and the final period lasted longer (Fig. 2). These changes were greater when the initial frequency of the *C_R_* allele was higher (Fig. 2). Ultimately, these changes did not alter the population’s extinction time when starting from a ZW/ZZ system (although a few populations in the 90% C_R_ scenario died out prematurely; see SI5 Fig. S16f), but the XX/XY system starting with a widespread *C_R_* allele survived slightly longer (Fig. 2). Notably, however, all these differences in duration were biologically small, averaging only a few generations (SI1 Table S3).

**Figure 2.**
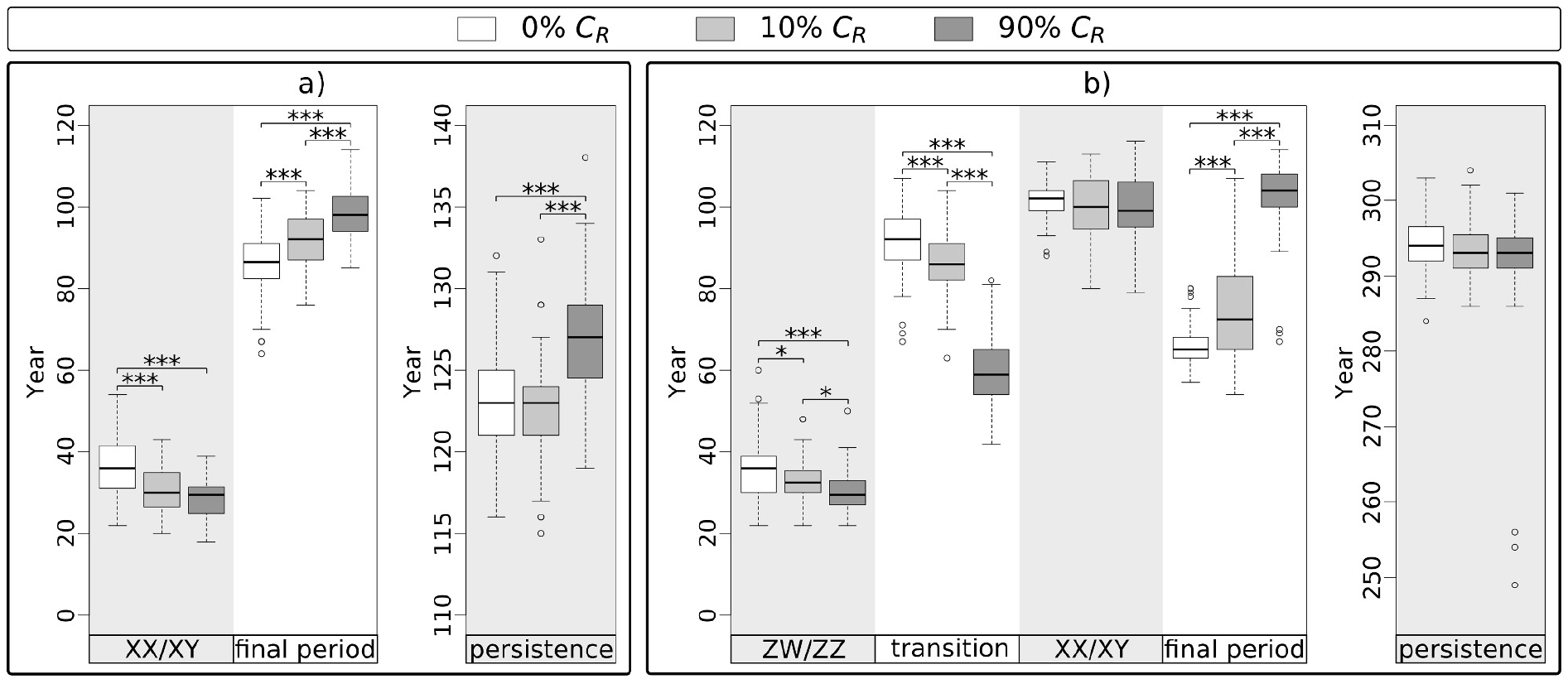
Length of each sex-determination period and overall population persistence in each scenario. Each box plot shows the distribution (thick middle line: median, box: interquartile range; whiskers extend to the most extreme data points within 1.5 × interquartile range from the box) of the results of 100 simulations, starting with XX/XY system (a) or ZW/ZZ system (b). Significant pairwise differences are indicated above the box plots as: * 0.01<p<0.05, *** p<0.001 (for details see SI1 Table S3). Generation time was 3 years. Year zero refers to the start of climate warming.

Average ASR was not affected by the presence of allele *C_R_* during the initial ZW/ZZ and XX/XY periods, nor during the XX/XY period evolved from the ZW/ZZ system: trivially, ASR remained near 0.5 in these periods as these were defined by the scarcity of sex-reversed males and thus little variation in female choice (Fig. 1, SI1 Table S3). However, during the mixed sexdetermination period following the initial sex-chromosome system, the presence and frequency of allele *C_R_* significantly influenced when and how much ASR shifted towards males (Fig. 1, Fig. 3, SI1 Table S3). When the initial frequency of the C_R_ allele was 10%, it had little effect on ASR in the XX/XY system (Fig. 1b), but populations starting with ZW/ZZ system had significantly less male-biased ASR during the transitional period between ZW/ZZ and XX/XY (on average, 63% males instead of 72%) and reached 0.6 ASR about 3 generations later than populations where all females preferred normal males (Fig. 1e, Fig. 3). When the initial frequency of allele *C_R_* was 90%, it had even greater effects on ASR (Fig. 1f, Fig. 3). First, in both systems, ASR decreased slightly below 0.5 temporarily after the initial sex-determination system ended, returning then to 0.5 (Fig. 1c, f). After that, ASR remained close to 0.5 throughout the transition period following the ZW/ZZ system, keeping the population in a balanced sex ratio for ca. 50 generations longer compared to the other two scenarios (Fig. 1d-f, Fig. 3a). Starting with the XX/XY system, ASR increased to 0.6 slightly later when allele *C_R_* was widespread in the initial population compared to scenario 0% *C_R_* (Fig. 1a-c, Fig. 3a), although the slope of ASR increase was alternatingly steeper and shallower before shifting ultimately to 1 (Fig. 1a-c).

**Figure 3.**
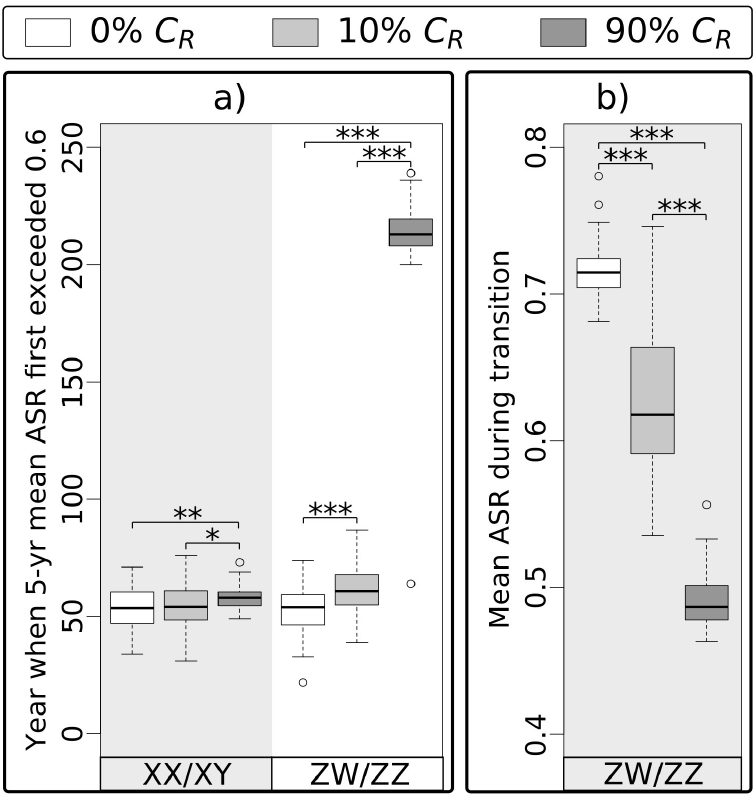
Differences in adult sex ratios (ASR, proportion of males) among scenarios. The year when mean ASR across 5 consecutive years first exceeded 0.6, starting with either XX/XY or ZW/ZZ system (a). Mean ASR during the transition period between the initial ZW/ZZ and a newly formed XX/XY system (b). Significant pairwise differences are indicated above the box plots as: * p<0.05, ** p<0.01, *** p<0.001 (for details see SI1 Table S3). Generation time was 3 years. Year zero refers to the start of climate warming.

### Background of preference effects

Starting with XX/XY system and a rare (10%) dominant *C_R_* allele, the accumulation of masculinized individuals was slightly sped up by the presence of females preferring them. Because individuals possessing the *thr_low_* allele were the first to masculinize, they were the ones chosen by females possessing *C_R_*, leading to positive LD between *thr_low_* and *C_R_* (Fig. 4a) and keeping *thr_low_* from decreasing for several decades (Fig. 1b), which resulted in earlier transition into the mixed system of the final period. However, the rare *C_R_* allele had little if any effect on ASR, because its spread was selected against for the following reasons. During the first decades, masculinized individuals (*aa* males) and the females choosing them (females with *C_R_*) produced female-biased offspring (all *aa* genotypes, facing relatively low masculinization rates at this early stage), which was not beneficial because the ASR was still close to 0.5 (Fig. 5a). As ASR became more and more male-biased, sex-ratio selection increasingly favoured *C_R_* (Fig. 5a), but sexual selection acted against it for the following reasons. Because females carrying *C_R_* preferred to mate with sex-reversed males (*aa*, i.e. those without chromosome A), negative LD arose between *C_R_* and *A* (Fig. 4a). As the majority of females did not possess allele *C_R_* and thus preferred normal males, sexual selection favoured males with chromosome *A*, and thereby acted against *C_R_* due to the negative LD between *C_R_* and A. Thus, males carrying *C_R_* were less likely to become fathers than those without *C_R_* (i.e. proportion of *C_R_* was lower among the alleles passed on by fathers than among the alleles in adult males; see light and dark blue lines in Fig. 5a) and consequently the proportion of *C_R_* inherited by offspring from their fathers was lower than expected based on its proportion among adult males (Fig. 5a). Therefore, the frequency of allele *C_R_* decreased significantly (on average by 0.032, 95% confidence interval [CI]: 0.025-0.039, t_99_ = 8.56, p<0.001) from the start of the final mixed-system period until the beginning of ultimate population decline, and the *C_R_* allele generally vanished before population extinction (Fig. 1b, SI5 Fig. S15a).

**Figure 4.**
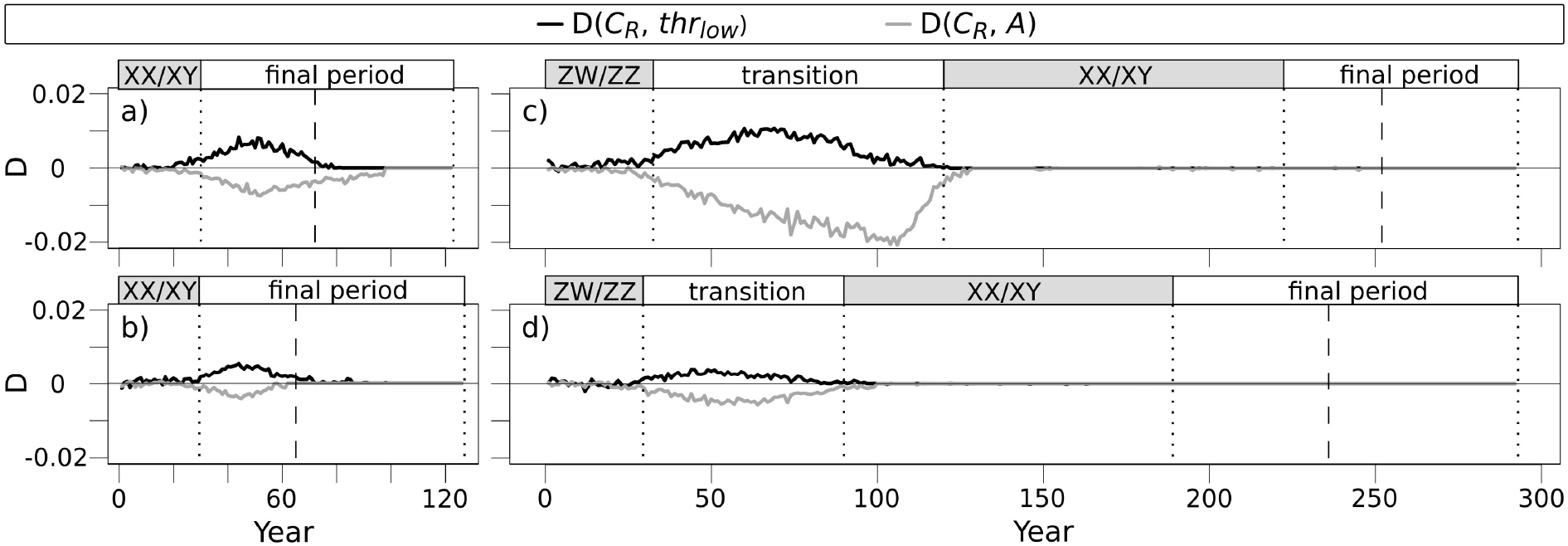
Linkage disequilibrium coefficient (D) between preference allele *C_R_* and either allele *thr_low_* or sex chromosome *A*. On the left: simulations starting with XX/XY system with scenarios 10% *C_R_* (a) and 90% *C_R_* (b). On the right: simulations starting with ZW/ZZ system with scenarios 10% *C_R_* (c) and 90% *C_R_* (d). Each curve indicates median values calculated from 100 simulations. Vertical dotted lines indicate the end of each sex-determination period and the dashed lines indicate the start of ultimate population decline. Note that negative linkage disequilibrium (LD) with A is equivalent with positive LD with a. Year zero refers to the start of climate warming.

When the XX/XY system started with a common (90%) recessive *C_R_* allele, more drastic changes occurred. Since the majority of females preferred sex-reversed males, as soon as the latter started to spread they flooded the population with female-biased offspring that skewed the ASR towards females (Fig. 1c) and passed on *C_R_* and *thr_low_*, increasing the frequency of both alleles significantly (Fig. 1c, Fig. 5b) and creating LD between *C_R_* and *A* and between *C_R_* and *thr_low_* (Fig. 4b). Due to the widespread preference for *aa* males and the positive LD between a and *C_R_*, the relative frequency of *C_R_* was higher in males becoming fathers than in the adult male population (Fig. 5b); thus, sexual selection spread *C_R_* (Fig. 5b; average increment in *C_R_* frequency was 0.029, 95% CI: 0.020-0.039, t_99_=6.132, p<0.001) despite its disadvantage in sex-ratio selection while climate warming was relatively mild. The spread of *C_R_* was accompanied by a rapid shift towards male-biased ASR because the widepsread preference for sex-reversed males increased the frequency of *thr_low_* to ca. 80% (Fig. 1c). Notably, sexual selection for the lack of chromosome A (Y) eradicated it from the population about 20 generations after the start of climate warming, leaving only *aa* genotypes (Fig. 1c) and ending all selection on *C_R_* (Fig. 5b). At this point, the role of the sex-chromosome-linked sex-determining locus was taken over by the autosomal threshold locus, as individuals possessing (more copies of) thrhigh could resist masculinization for some time. During this time, sex-ratio selection favoured thrhigh and started to spread it until fixation, which temporarily slowed the increase of masculinization rate and ASR (SI5 Fig. S13c, Fig. 1c), maintaining somewhat higher effective population size compared to scenarios with no or rare *C_R_* (SI5 Fig. S16-17), and thereby slightly increasing the population’s survival time (Fig. 2a).

**Figure 5.**
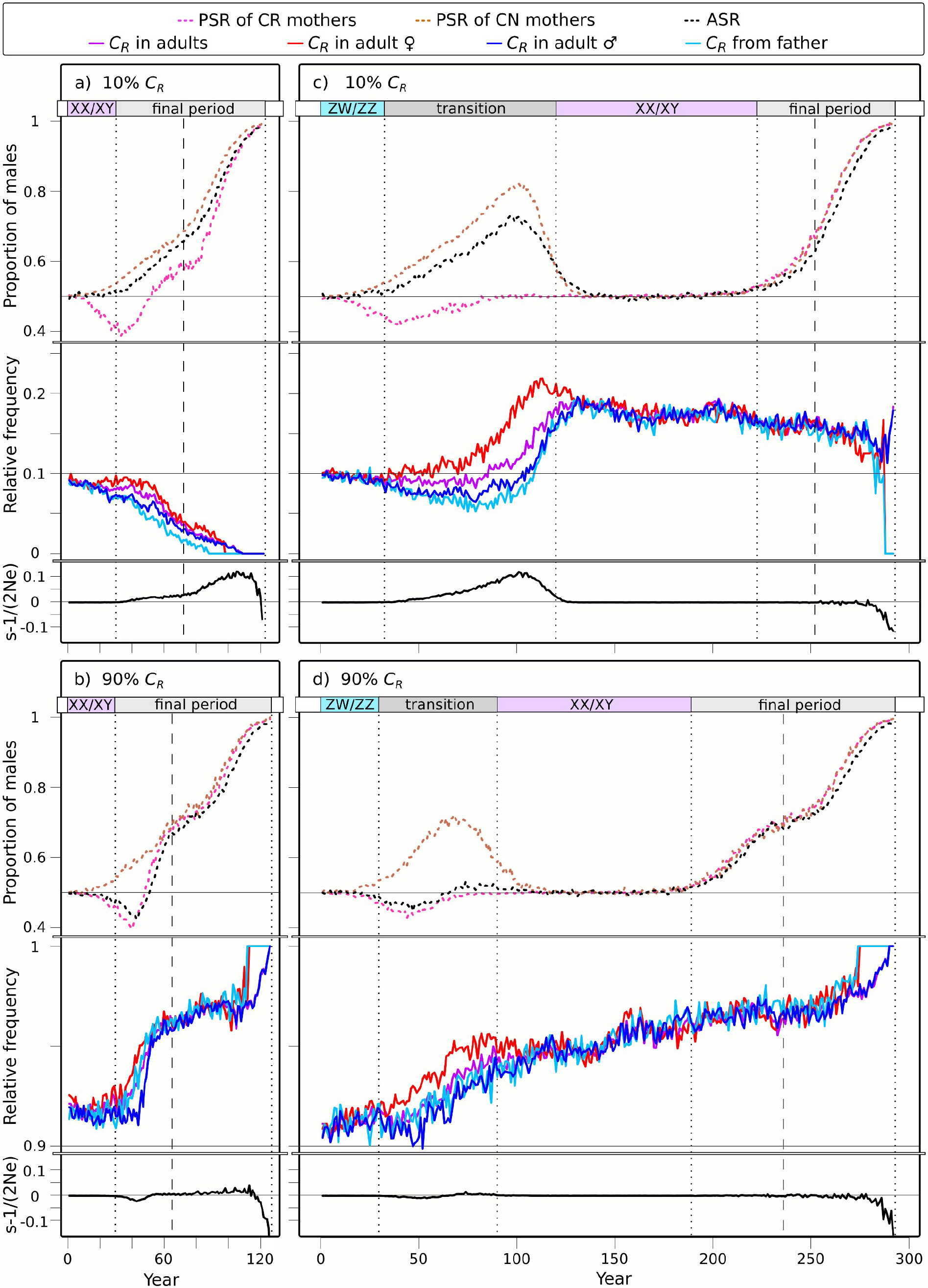
Consequences of mate choice: progeny sex ratio, inheritance of allele *C_R_* and sex-ratio selection acting on allele *C_R_*. On the left: simulations starting with XX/XY system with scenarios 10% *C_R_* (a) and 90% *C_R_* (b). On the right: simulations starting with ZW/ZZ system with scenarios 10% *C_R_* (c) and 90% *C_R_* (d). In each panel, the top three curves show the changes in adult sex ratio (ASR) and progeny sex ratio (PSR) of mothers preferring sex-reversed and normal males (CR and CN mothers, respectively); the four curves in the middle show the relative frequency of allele *C_R_* across and among sexes (*C_R_* in adults, *C_R_* in adult ♀, *C_R_* in adult ♂) and relative frequency of *C_R_* among preference alleles inherited by offspring from their fathers (*C_R_* from father; note how it differs from *C_R_* in adult ♂). Mothers passed on allele *C_R_* with a relative frequency corresponding to its presence among adult females (not shown). The bottom curve of each panel shows the strength of sex-ratio selection (s) relative to the effective population size (Ne), expressed as *s*-1/(2Ne) (this value is shown instead of s itself, because the effect of genetic drift can override s in small populations). Each curve indicates median values calculated from 100 simulations (see SI5 Fig. S15 for the relative frequencies of *C_R_* in all simulations). Vertical dotted lines indicate the end of each sex-determination period and dashed lines indicate the start of ultimate population decline. Note that the top, middle, and bottom part of each panel has three different Y axes and with different scales. Year zero refers to the start of climate warming.

Starting with ZW/ZZ system and dominant allele *C_R_* at 10% frequency, similar LD of *C_R_* with chromosome *a* (W) and allele *thr_low_* (Fig. 4c) as described above slowed the decrease of *thr_low_* and accelerated the accumulation of *aa* genotypes and thereby the transition into XX/XY system (Fig. 1e). Because not all females preferred normal (*AA*) males, this scenario produced a milder excess of *Aa* genotypes during the transition period (Fig. 1e) compared to the scenario without *C_R_* (Fig. 1d), and the slightly better mating success of masculinized individuals resulted in faster accumulation of “resistant females” (*aa*), slowing the ASR increase and keeping it less male-biased during the transition period (Fig. 1e, Fig. 3b). Nevertheless, ASR was skewed enough so sex-ratio selection favoured parents that produced less male-biased offspring, i.e. females choosing masculinized individuals, thus the frequency of *C_R_* almost doubled by the end of the transition period (Fig. 5c; during ZW/ZZ to XX/XY transition, relative frequency of allele *C_R_* increased on average by 0.076, 95% CI: 0.059-0.093; paired t-test: t_99_ = 9.0, p<0.001). Once the *AA* genotype went extinct, both sex-ratio selection for allele *C_R_* (and chromosome *a*; Fig. 4c) and sexual selection against it ended, and the further processes followed a similar course as in the scenario without *C_R_* (Fig. 1d-e).

When the ZW/ZZ system started with the recessive *C_R_* allele at 90% frequency, the effects seen in the previous scenario became stronger, *via* similar mechanisms. Since the majority of females preferred masculinized individuals, the accumulation of *aa* genotypes and the transition to XX/XY system were even faster (Fig. 1f, Fig. 2b). During the transition period, there was only a small excess of *Aa* genotypes (Fig. 1f) because *AA* males were soon replaced by *Aa* males as the latter spread and were preferred by most females (who increasingly had the *aa* genotype), but the frequency of *thr_low_* increased greatly due to the widespread preference for individuals with low masculinization thresholds (Fig. 1f). Since masculinized individuals enjoyed high mating success, ASR became slightly female-biased at the start of the transition period, but soon returned to 0.5 and remained close to it afterwards (Fig. 1f, Fig. 3b) because the production of “resistant females” compensated for the increasing masculinization rate. The frequency of allele *C_R_* slightly increased during the transition period (by 0.021 on average, 95% CI: 0.012-0.030, t_99_ = 4.707, p<0.001; Fig. 5d) due to sexual selection (i.e. the more preferred sex-reversed males carried allele *C_R_* and passed it on to the next generation with higher than expected probability: Fig. 5d) and LD with chromosome a (W) and allele *thr_low_* (Fig. 4d), until genotype *AA* went extinct. Overall preference for sex-reversed males kept the frequency of *thr_low_* around 80% until nearly 75 generations after the start of climate warming, causing a relatively early and sudden ASR increment after the XX/XY system ceased to persist (Fig. 1f). Thereafter, sex-ratio selection favoured allele *thr_high_* over *thr_low_*, but could not prevent the population from a relatively early decline compared to the other scenarios (Fig. 1f, SI5 Fig. S16f). Thus, when chromosome A went extinct and the role of the sex-determining locus was taken over by the autosomal threshold locus, *thr_high_* was already close to fixation and so this new sex-determination system persisted for only a few generations before the population died out (Fig. 1f).

### Sensitivity analysis

The above-described effects of *C_R_* were very similar when we assumed a multiallelic *thr* locus (instead of biallelic) or intermediate inheritance of *C* alleles (instead of fully dominant/recessive), or when the value of *C_N_* was set to 0.5, encoding indiscriminate mating, or when viability of WW offspring was reduced by 25% or 50% (SI2 Fig. S2-7). Changes in relative frequency of *C_R_* were also very similar (SI2 Fig. S2-7), except that it showed little if any change over time when initial frequency of *C_R_* was 10% and *C_N_* encoded indiscriminate mating (SI2 Fig. S3-4).

However, when WW viability was reduced by 75%, presence or absence of *C_R_* made considerable difference (SI2 Fig. S8; Fig. 6). When *C_R_* was absent, the population could transition to an XX/XY system and persist for ca. 270 years in only 26% of simulations; the rest died out after ca. 120 years (Fig. 6). When *C_R_* was present, it saved the population from early extinction by enabling the switch to XX/XY system in 56% of simulations when *C_R_* was rare and in 100% when *C_R_* was widespread (Fig. 6). Due to high mortality of WW offspring, relative frequency of *C_R_* started to decrease in both scenarios, but it increased rapidly just before the end of transition when ASR became highly male-biased (SI2 Fig. S8).

**Figure 6.**
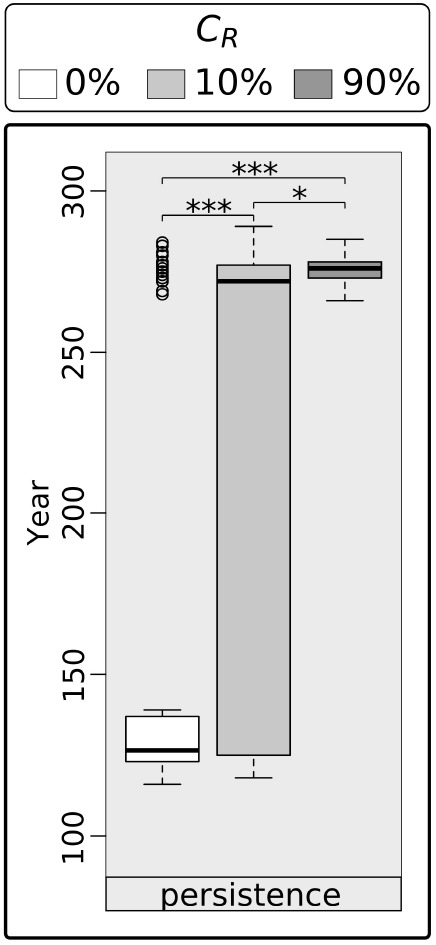
Population persistence in ZZ/ZW systems with 25% WW viability. Each box plot shows the distribution (thick middle line: median, box: interquartile range; whiskers extend to the most extreme data points within 1.5 × interquartile range from the box) of the results of 100 simulations. Significant pairwise differences are indicated above the box plots as: * 0.01<p<0.05, *** p<0.001. Generation time was 3 years. Year zero refers to the start of climate warming.

In contrast, when we assumed that the WW genotype was not viable, the ZW/ZZ system always behaved like the XX/XY system (no transition in sex determination; population extinction after ca. 120 years of warming) and the presence of *C_R_* had very little effect on the shifting of ASR towards males (SI2 Fig. S9). Relative frequency of both the rare and the widespread *C_R_* decreased steadily in these scenarios as females mating with sex-reversed males lost 25% of their offspring because of being WW (SI2 Fig. S9).

## Discussion

We investigated how female preference for sex-reversed males would affect evolution in an increasingly masculinizing environment. Our simulations showed that the presence and frequency of such a preference allele (*C_R_*) influenced both the temporal dynamics of evolution of sex-determination systems and the changes in adult sex ratio across a wide range of circumstances, and under certain conditions it also affected the timing of population extinction. Furthermore, we found that a rare, dominant *C_R_* allele may spread in populations with ZW/ZZ sex-determination sytem more likely than in populations with XX/XY system. We discuss each of these main findings in detail below.

In our simulations, increasing masculinization rate under continuous climate warming resulted in a process where different sex-determination systems replaced one another. This agrees with the findings of previous models [11–14] and experimental data [16] suggesting that climate change may cause turnovers between different sex-determination systems and thus may have contributed to the variability of sex-determination systems across ectothermic vertebrates [27, 45, 46]. Our present results demonstrate that the speed of these turnovers may be enhanced by sexual selection if females can recognize and prefer to mate with sex-reversed males. Specifically, in our simulations, the initial sex-determination system (either XX/XY or ZW/ZZ) evolved sooner into a mixed sex-determination system, and when the initial system was ZW/ZZ, the transitional mixed system also evolved sooner into an XX/XY system when *C_R_* allele was present in the population. These effects were stronger when the frequency of *C_R_* was higher. Furthermore, a widespread *C_R_* allele facilitated a turnover in the XX/XY system that was not seen when *C_R_* was rare or absent: the original male-determining sex chromosome went extinct and its role was taken over by an autosome (harbouring the original threshold locus). These results suggest the idea that, over the evolutionary past with alternatingly warmer and colder climates, variation in female preferences for sex-linked traits might have catalized the diversification of sex-determination systems in taxa that are prone for sex reversals. Further models could test this idea by simulating climate warming followed by a period of stable climate [15].

As the production of viable WW offspring, which can resist masculinization for long, was key to the transiton from ZZ/ZW to XX/XY systems, considerably reduced WW viability had major impacts on the outcomes. Complete WW lethality erased any difference between the ZZ/ZW and XX/XY systems and any effect of *C_R_*, whereas 25% WW viability resulted in strikingly divergent fates with and without *C_R_*, whereby populations possessing *C_R_* had much better chances of persisting for more than twice as long as populations without *C_R_*. This shows that sexual selection for sex-reversed males may prevent premature extinction in certain circumstances. However, when the WW genotype was at least fairly viable, *C_R_* had very little effect on population persistence in our simulations. The latest possible extinction time was determined by the value of allele *thr_high_* and *b_sig_* (which were fixed in our simulations): when climate warming caused masculinization in all *thr_high_* homozygotes, no more female offspring could be produced, after which the population could persist only as long as the remaining females survived. Populations with ZW/ZZ initial system persisted longer when they could transition into an XX/XY system, but within both initial systems, females disappeared from the populations at roughly the same time in all scenarios. The only exception in scenarios with viable WW was that extinction happened a few years later when *C_R_* had high frequency in populations with XX/XY initial system, where there were slightly more females (i.e. non-masculinized XX individuals) in the final years before population extinction, because with the disappearance of chromosome Y only individuals with low endogenous male signal levels remained. Although population persistence may be prolonged by occurrence of new, mutant alleles causing lower male signal levels or higher threshold levels, the time needed for such evolutionary changes was typically long in previous models [12–14]. We did not allow for such mutations in our model because we assumed that the rapid climate change would not allow enough time for new mutants to appear and spread before extinction in our small populations. Bearing this caveat in mind, our results suggest that female preference for sex-reversed males alone cannot grant much longer persistence for a population under continuous, rapid climate warming if the WW genotype does not suffer from markedly increased mortality rate. However, in our simulations, population persistence was not threatened by anything else than climate-driven masculinization. In reality, further effects of climate change or other perturbations may also harm the populations [47], and resilience against these perturbations might be affected by *C_R_*, as discussed next.

We found that female preferences can have a significant effect on changes of adult sex ratio during climate warming. While the male to female ratio was around 1:1 during ZW/ZZ and XX/XY sex-determination periods, it markedly shifted towards males in the mixed periods in scenarios where all females favoured normal males. By contrast, presence of allele *C_R_* prolonged the time before the ASR became strongly male-biased (except when starting from an XX/XY system with a rare *C_R_*), and during the transition from ZZ/ZW to XX/XY system it kept the ASR less male-biased (when *C_R_* was rare or when WW had reduced viability) or close to a healthy 0.5 (when *C_R_* was widespread). This way, the presence of allele *C_R_* helped maintaining a higher effective population size when populations without *C_R_* suffered population bottlenecks. Populations with less biased ASR and higher effective population size are more likely to survive environmental perturbations such as anthropogenic habitat loss and disease outbreaks that currently parallel and interact with the effects of global climate change [47]. In this respect, our results suggest that ZW/ZZ populations with widespread preference for sex-reversed males might have the highest adaptive potential and the best chances to cope with contemporary climate change. Understanding the factors underlying the variance in adaptability is important because a recent meta-analysis suggests that animals’ adaptive responses to climate change may be insufficient [1].

Our present results support the previous findings that XX/XY and ZW/ZZ systems may differ in their responses to climate change [11, 14]. Which of the two systems is more resilient depends on several conditions, such as the fertility of sex-reversed males and the viability of WW individuals [11]. Here we found that the ZW/ZZ system may maintain healthier sex ratios for longer than the XX/XY system if the sex-reversed males can reproduce like normal males and produce viable, fertile WW offspring that are resistant to masculinization, and this allowed the former to persist more than two times longer than the latter. Although our knowledge on WW or masculinized individuals in nature is scarce, there is a limited number of studies suggesting that the above conditions stand in various taxa [10, 17, 27, 35–37, 48]. Morover, most ectothermic vertebrates possess homorphic sex chromosomes that show no signs of degeneration, suggesting that sex-reversed and WW individuals should be viable and fertile in the majority of these taxa [4, 38, 45]. Under these conditions, our results confirm that female-heterogametic systems may be less vulnerable in a masculinizing environment compared to male-heterogametic systems [11]. Our present model contributes to this picture with the finding that the frequency of female preference for sex-reversed males is a further condition that may lead to different effects of climate change in the two sex-chromosome systems. For example, after ca. 60 years of warming, while both systems had similarly male-biased ASR when *C_R_* was rare, the ZZ/ZW system had much more balanced ASR than the XX/XY system when *C_R_* was widespread (Fig. 1). Furthermore, our results show that the two systems respond to climate warming identically when the WW genotype is lethal, and also when WW viability is poor and *C_R_* is absent, but the ZZ/ZW system is more likely to outlive the XX/XY system even with poorly viable WW if *C_R_* is present.

A further difference between the two sex-chromosome systems in our model was seen in the changes in *C_R_* frequency when *C_R_* was competing with an allele encoding preference for normal males. We found that a rare *C_R_* allele spread in the population when the initial sexdetermination system was ZW/ZZ, but tended to disappear instead when sex determination was initially XX/XY. The major difference between the two systems was that sex-ratio selection that favoured allele *C_R_* was stronger in the ZW/ZZ system, because females carrying *C_R_* produced WW offspring that were resistant to masculinization, thus their progeny sex ratios were more advantageous when ASR was male-biased. This stronger sex-ratio selection in the ZW/ZZ system counteracted the prevailing sexual selection that acted against *C_R_* in both systems due to the widespread preference for normal males and the LD between *C_R_* and the sex chromosomes. Thus, our results suggest that a rare autosomal preference for sex-reversed males is more likely to spread in ZW/ZZ than XX/XY systems. This finding parallels previous theoretical studies that showed that the ZZ/ZW system is particularly prone to evolve sex-linked preferences for sexually antagonistic traits [49] and a new male ornament is more protected against random loss in ZW/ZZ compared to XX/XY systems [50].

By contrast, when *C_R_* was widespread in the population, sexual selection in both systems favoured sex-reversed males and consequently allele *C_R_* that was more frequent in such males, while sex-ratio selection played virtually no role in the spread of this allele, because adult sex ratio hardly deviated from 0.5 as long as both normal and sex-reversed males were present. Both sexual selection and sex-ratio selection have long been known to be major driving forces of species evolution [41, 51, 52]; and these forces together can even lead to speciation [53]. Occurrence of novel combinations of genetic sex and sex-linked coloration under sexual selection can lead to rapid sympatric speciation [54]; thus, our findings raise the possibility that climate-driven sex reversals might contribute to speciation. Because sex-ratio selection, which spread *C_R_* in our simulated populations, was due to the masculinizing effect of rising environmental temperature, our results demonstrate that climate change may influence the evolution of female mate choice. This finding parallels the conclusions of a modelling study showing that environmental pollution may disrupt sexual selection and thereby decrease population fitness [55]. Taken together, these theoretical results highlight that various forms of ongoing anthropogenic environmental change worldwide may be driving changes in mating preferences, which then can have knock-on consequences on adaptive potential and population viability.

Our main conclusion is that sexual selection for sex-linked traits may influence the effects of climate change on the demography and evolution of populations with temperature-sensitive sex development. This provokes several further questions for future studies. On the one hand, the genetic architecture of sexually selected traits and the genetic and environmental determinants of sex are poorly known for many taxa. To assess how much wildlife is at risk by climate change, we need more empirical information filling these knowledge gaps. On the other hand, we also need theoretical studies on how further factors affect our predictions. For example, in species where the temperature reaction norm is non-linear, climate warming may lead to ZZ feminization and ultimately the loss of W chromosome [15]. In bearded dragons (*Pogona vitticeps*), for instance, ZZ individuals develop into females at high temperatures, and surprisingly, these sex-reversed females enjoy a fecundity advantage over normal females [16]. In such case, males might prefer sex-reversed females, which may complicate the effects of climate warming similarly to what we found here for female choice. As growing evidence suggests that sex reversal is more likely widespread rather than a rare oddity in ectotherms [8], exploring the complexity of its consequences is an important emerging research avenue.

## Acknowledgements

The study was funded by the National Research, Development and Innovation Office of Hungary (NKFIH 115402) and EN was financially supported by the Ministry of Human Capacities (National Program for Talent of Hungary, NTP-NFTÖ 17-B-0317). We are grateful to Lisa E. Schwanz and three anonymous referees for constructive comments on earlier versions of the manuscript.

## Authors’ contributions

The study was conceived and designed by EN, VB and SK. The R code was written by EN with help from SK. Statistical analyses were performed by EN with help from VB. The first draft was written by EN and VB, and all authors contributed to finalizing the manuscript.

## References

1. Radchuk V, Reed T, Teplitsky C, van de Pol M, Charmantier A, Hassall C, et al. Adaptive responses of animals to climate change are most likely insufficient. Nat Commun. 2019;10:3109. doi:10.1038/s41467-019-10924-4.

2. Mitchell NJ, Janzen FJ. Temperature-dependent sex determination and contemporary climate change. Sex Dev. 2010;4:129–40.

3. Ospina-Álvarez N, Piferrer F. Temperature-dependent sex determination in fish revisited: Prevalence, a single sex ratio response pattern, and possible effects of climate change. PLoS One. 2008;3:2–4.

4. Eggert C. Sex determination: the amphibian models. Reprod Nutr Dev. 2004;44:539–49. doi:10.1051/rnd:2004062.

5. Alho JS, Matsuba C, Merilä J. Sex reversal and primary sex ratios in the common frog (Rana temporaria). Mol Ecol. 2010;19:1763–73. doi:10.1111/j.1365-294X.2010.04607.x.

6. Lambert MR, Tran T, Kilian A, Ezaz T, Skelly DK. Molecular evidence for sex reversal in wild populations of green frogs (*Rana clamitans*). PeerJ. 2019;7:e6449. doi:10.7717/peerj.6449.

7. Nemesházi E, Gál Z, Ujhegyi N, Verebélyi V, Mikó Z, Üveges B, et al. Novel genetic sex markers reveal high frequency of sex reversal in wild populations of the agile frog (*Rana dalmatina*) associated with anthropogenic land use. Preprint at https://doi.org/10.22541/au.158775693.35677255/v2 (2020).

8. Holleley CE, Sarre SD, O’Meally D, Georges A. Sex reversal in reptiles: reproductive oddity or powerful driver of evolutionary change? Sex Dev. 2016;10:279–87.

9. Reinboth R, editor. Intersexuality in the Animal Kingdom. Berlin, Heidelberg: Springer Berlin Heidelberg; 1975. doi:10.1007/978-3-642-66069-6.

10. Chardard D, Penrad-Mobayed M, Chesnel A, Pieau C, Dournon C. Thermal sex reversals in amphibians. In: Valenzuela N, Lance V, editors. Temperature-dependent sex determination in vertebrates. Washington: Smithsonian Books; 2004. p. 59–67.

11. Bókony V, Kövér S, Nemesházi E, Liker A, Székely T. Climate-driven shifts in adult sex ratios via sex reversals: the type of sex determination matters. Philos Trans R Soc B Biol Sci. 2017;372:20160325.

12. Quinn AE, Sarre SD, Ezaz T, Marshall Graves J a, Georges A. Evolutionary transitions between mechanisms of sex determination in vertebrates. Biol Lett. 2011;7:443–8.

13. Grossen C, Neuenschwander S, Perrin N. Temperature-dependent turnovers in sex-determination mechanisms: a quantitative model. Evolution (N Y). 2011;65:64–78. doi:10.1111/j.1558-5646.2010.01098.x.

14. Schwanz LE, Ezaz T, Gruber B, Georges A. Novel evolutionary pathways of sex-determining mechanisms. J Evol Biol. 2013;26:2544–57.

15. Schwanz LE, Georges A, Holleley CE, Sarre SD. Climate change, sex reversal and lability of sex-determining systems. J Evol Biol. 2020;33:270–81.

16. Holleley CE, O’Meally D, Sarre SD, Marshall-Graves JA, Ezaz T, Matsubara K, et al. Sex reversal triggers the rapid transition from genetic to temperature-dependent sex. Nature. 2015;523:79–82. doi:10.1038/nature14574.

17. Parnell NF, Streelman JT. Genetic interactions controlling sex and color establish the potential for sexual conflict in Lake Malawi cichlid fishes. Heredity (Edinb). 2013;110:239–46. doi:10.1038/hdy.2012.73.

18. Senior AM, Nat Lim J, Nakagawa S. The fitness consequences of environmental sex reversal in fish: a quantitative review. Biol Rev. 2012;87:900–11.

19. Lindholm A, Breden F. Sex chromosomes and sexual selection in poeciliid fishes. Am Nat. 2002;160:S214–24.

20. McKinnon JS, Pierotti ME. Colour polymorphism and correlated characters: genetic mechanisms and evolution. Mol Ecol. 2010;19:5101–25.

21. Kottler VA, Schartl M. The colorful sex chromosomes of teleost fish. Genes (Basel). 2018;9:233.

22. Kirkpatrick M, Hall DW. Sexual selection and sex linkage. Evolution (N Y). 2004;58:683–91.

23. Savin SM. The history of the Earth’s surface temperature during the past 100 million years. Annu Rev Earth Planet Sci. 1977;5:319–55.

24. Hansen JE, Sato M. Paleoclimate implications for human-made climate change. In: Climate Change. Vienna: Springer Vienna; 2012. p. 21–47. doi:10.1007/978-3-7091-0973-1_2.

25. England MH, Kajtar JB, Maher N. Robust warming projections despite the recent hiatus. Nat Clim Chang. 2015;5:394–6. doi:10.1038/nclimate2575.

26. Cheng L, Abraham J, Hausfather Z, Trenberth KE. How fast are the oceans warming? Science (80-). 2019;363:128–9.

27. Devlin RH, Nagahama Y. Sex determination and sex differentiation in fish: an overview of genetic, physiological, and environmental influences. Aquaculture. 2002;208:191–364.

28. Bull JJ, Vogt RC, Bulmer MG. Heritability of sex ratio in turtles with environmental sex determination. Evolution (N Y). 1982;36:333–41.

29. Janzen FJ. Heritable variation for sex ratio under environmental sex determination in the common snapping turtle (Chelydra serpentina). Genetics. 1992;131:155–61.

30. Schroeder AL, Metzger KJ, Miller A, Rhen T. A novel candidate gene for temperature-dependent sex determination in the common snapping turtle. Genetics. 2016;203:557–71.

31. Wessels S, Sharifi RA, Luehmann LM, Rueangsri S, Krause I, Pach S, et al. Allelic variant in the anti-Müllerian hormone gene leads to autosomal and temperature-dependent sex reversal in a selected nile tilapia line. PLoS One. 2014;9:e114341.

32. Chandler CH, Phillips PC, Janzen FJ. The evolution of sex-determining mechanisms: lessons from temperature-sensitive mutations in sex determination genes in Caenorhabditis elegans. J Evol Biol. 2009;22:192–200.

33. Chardard D, Dournon C. Sex reversal by aromatase inhibitor treatment in the newt Pleurodeles waltl. J Exp Zool. 1999;283:43–50.

34. Senior AM, Johnson SL, Nakagawa S. Sperm traits of masculinized fish relative to wild-type males: a systematic review and meta-analyses. Fish Fish. 2016;17:143–64.

35. Kallman KD. A new look at sex determination in Poeciliid fishes. In: Turner BJ, editor. Evolutionary Genetics of Fishes. New York: Plenum Press, New York; 1984. p. 95–171.

36. Wallace H, Badawy GMI, Wallace BMN. Amphibian sex determination and sex reversal. Cell Mol Life Sci. 1999;55:901–9. doi:10.1007/s000180050343.

37. Roco ÁS, Olmstead AW, Degitz SJ, Amano T, Zimmerman LB, Bullejos M. Coexistence of Y, W, and Z sex chromosomes in *Xenopus tropicalis*. Proc Natl Acad Sci U S A. 2015;112:E4752–4761. doi:10.1073/pnas.1505291112.

38. Ezaz T, Stiglec R, Veyrunes F, Marshall Graves JA. Relationships between vertebrate ZW and XY sex chromosome systems. Curr Biol. 2006;16:R736–43. doi:10.1016/j.cub.2006.08.021.

39. Morice CP, Kennedy JJ, Rayner NA, Jones PD. Quantifying uncertainties in global and regional temperature change using an ensemble of observational estimates: the HadCRUT4 data set. J Geophys Res Atmos. 2012;117:1–22.

40. Perrin N. Random sex determination: When developmental noise tips the sex balance. BioEssays. 2016;38:1218–26.

41. Ryan MJ, Keddy-Hector A. Directional patterns of female mate choice and the role of sensory biases. Am Nat. 1992;139:S4–35. https://www.jstor.org/stable/2462426.

42. Basolo AL. Phylogenetic evidence for the role of a pre-existing bias in sexual selection. Proc R Soc B Biol Sci. 1995;259:307–11.

43. R Core Team. R: A language and environment for statistical computing. R ver. 3.1.2. R Foundation for Statistical Computing, Vienna, Austria. http://www.r-project.org; 2014.

44. Lenth R V. Least-squares means: the R Package lsmeans. J Stat Softw. 2016;69:1–33. doi:10.18637/jss.v069.i01.

45. Hillis DM, Green DM. Evolutionary changes of heterogametic sex in the phylogenetic history of amphibians. J Evol Biol. 1990;3:49–64.

46. Miura I. Sex Determination and sex chromosomes in Amphibia. Sex Dev. 2017;8526:298–306. doi:10.1159/000485270.

47. Hayes TB, Falso P, Gallipeau S, Stice M. The cause of global amphibian declines: a developmental endocrinologist’s perspective. J Exp Biol. 2010;213:921–33.

48. Veltsos P, Rodrigues N, Studer T, Ma W-J, Sermier R, Leuenberger J, et al. Y-chromosome haplotypes of varying differentiation to the X are not associated with male fitness in common frogs. bioRxiv. 2019. doi:10.1101/659565.

49. Muralidhar P. Mating preferences of selfish sex chromosomes. Nature. 2019;570:376–9. doi:10.1038/s41586-019-1271-7.

50. Reeve HK, Pfennig DW. Genetic biases for showy males: Are some genetic systems especially conducive to sexual selection? Proc Natl Acad Sci U S A. 2003;100:1089–94.

51. Maynard Smith J. Theories of sexual selection. Trends Ecol Evol. 1991;6:146–51.

52. Charnov EL. Sex ratio selection in an age-structured population. Evolution (N Y). 1975;29:366–8.

53. Seehausen O, Van Alphen JJM, Lande R. Color polymorphism and sex ratio distortion in a cichlid fish as an incipient stage in sympatric speciation by sexual selection. Ecol Lett. 1999;2:367–78.

54. Lande R, Seehausen O, van Alphen JJM. Mechanisms of rapid sympatric speciation by sex reversal and sexual selection in cichlid fish. Genetica. 2001;112–113:435–43.

55. Senior AM, Nakagawa S, Grimm V. The evolutionary consequences of disrupted male mating signals: an agent-based modelling exploration of endocrine disrupting chemicals in the guppy. PLoS One. 2014;9:e103100.

