## Supplementary material for "Evolutionary and demographic consequences of temperature-induced masculinization under climate warming: the effects of mate choice": SI1

**Table of contents:**

**Figures** ........................................................... **2**

Fig S1. Sex as a threshold trait and manifestation of female choosiness.

**Tables** ........................................................... **3**

Table S1. Examples for species exhibiting sex-linked body colour

Table S2. Model parameters and settings used in the simulations

Table S3. Linear contrasts

**Additinal information for Methods** ..................... **12**

Formulas for calculating effective population size, sex ratio selection, linkage and generation time.

**References** ........................................................... **15**

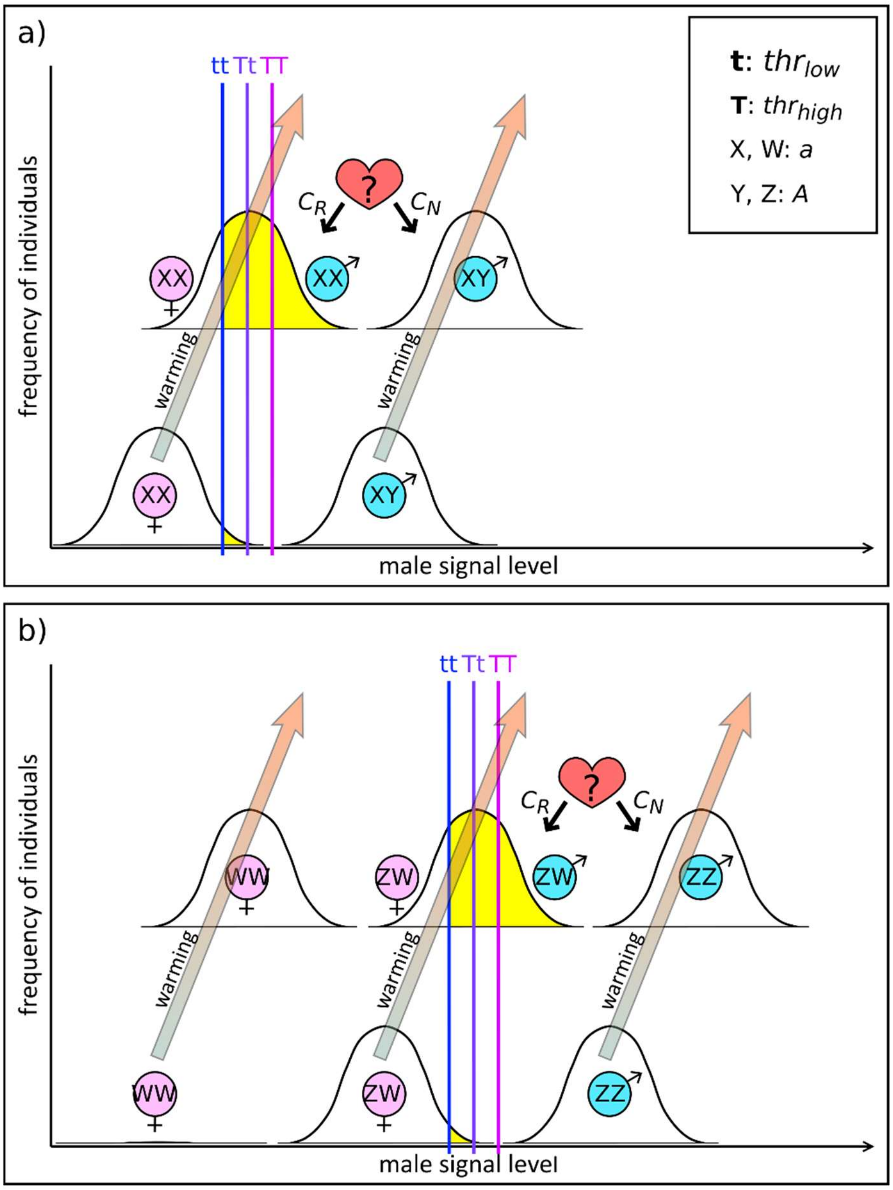


**Fig S1. Sex as a threshold trait and manifestation of female choosiness.** Sex chromosomes *A* and *a* produce different amounts of the 'male signal' factor. An individual becomes male if its male signal level exceeds its individual threshold defined by the individual's *thr* genotype (**tt**, **Tt** or **TT**). Production of male signal is increased by environmental temperature experienced during a sensitive period of early individual development; the Gaussian curves represent the distribution of individual signal levels due to variation in environmental temperatures. Values of the two *thr* alleles were set so individuals would generally develop sexual phenotype corresponding to their sex-chromosome genotype before climate warming (shown by the bottom curves in each panel), and only occasional events of masculinization would occur in some **tt** individuals (shown by yellow area under the curves). Accordingly, in an XX/XY system (a), *aa* individuals are predominantly females, and *Aa* individuals are males, while in a ZW/ZZ system (b), *Aa* individuals are females and *AA* individuals are males. Climate warming (illustrated by blue-orange arrows) increases the production of male signal by both sex chromosomes, resulting in more frequent masculinization (shown by the top curves in each panel). When both normal and sex-reversed males are present, females expressing allele $C_R$ will prefer sex-reversed males, while those expressing $C_N$ will more frequently choose normal males as mating partners.

**Table S1. Examples for species exhibiting sex-linked body colour.**

| Class | Species | Chromosome (system)[1] | Description | Publication |
|---|---|---|---|---|
| Insecta | *Danaus chrysippus* | neo-W (ZW) | In a hybrid zone of two subspecies *(D. c. chrysippus and dorippus)* in East Africa, fusion occurred in *D. c. chrysippus* between chromosome W and an autosome which previously coded a distinguishing colour of the two subspecies. | (Smith et al. 2016) |
| | *Papilio glaucus* | W (ZW) | *P. g. glaucus* females inherit a W-linked allele that causes melanic colour. Surprisingly, in *P. g. glaucus x canadensis* hybrids this melanism is suppressed. | (Hagen and Scriber 1989) |
| Actinopterygii[2] | *Cynotilapia afra* and *Pseudotrophe us elongatus* | W (ZW and XY) | Experimental hybridization of these species suggested that in a complex system of ZW/ZZ and XX/XY, the presence of chromosome W causes a female-like body colour. | (Parnell and Streelman 2013) |
| | *Neochromis omnicaeruleus* | X (XY) | Blotched pattern is determined by colour genes in linkage with X-linked female-determining genes (dominant over male-determining function of Y). There is a large variation in mating preferences for body colour across males collected from a single population. | (Pierotti et al. 2009) |
| | *Oryzias latipes* | Y (XY) | In d-rR strain medaka, white pigmentation is X-linked recessive trait, while orange pigmentation is Y-linked and dominant. Sex-reversed XY d-rR strain females show male-type (orange/red) pigmentation. | (Aida 1921; Edmunds et al. 2000) |
| | *Poecilia reticulata* | Y (XY) | There are several Y-linked body colour genes in this species, and female preference might have facilitated the reduction of X/Y recombination at these loci. | (Wright et al. 2017) |
| | *Xiphophorus pygmaeus* | Y (XY) | Alleles at a Y-linked gene cause either blue or gold body colour in males. Female preference for colour differs among populations. | (Kingston et al. 2003) |
| | *Gambusia holbrooki* | Y (XY) | Breeding tests suggest that the rare melanic body colour variant (mottled-black) is Y-linked and its expression is influenced by temperature. | (Horth 2006) |

| | | | | |
|---|---|---|---|---|
| Amphibia | *Rana rugosa* | X (XY) | The allele encoding a wild-type colour is dominant (can be present on both Y and X). A recessive allele coding whitish-yellow body colour was found on X. | (Miura et al. 2011) |
| Reptilia | *Python regius* | X and Y (XY) | Inheritance pattern of "coral glow" colour suggests sex-linkage of the responsible (incomplete dominant) allele with some level of recombination between X and Y. | (Mallery Jr. and Carrillo 2016) |
| Aves | *Erythrura gouldiae* | Z (ZW) | A Z-linked locus with two alleles determines head colour where red is dominant over black (two coexisting morphs). Mating preference for morph also seems to be Z-linked. | (Pryke 2009) |

[1] Chromosome: the sex chromosome harbouring a gene encoding a distinguishing body colour or pattern. System: sex determination system (ZW is short for ZW/ZZ and XY is short for XX/XY).

[2] Among fish, occurrence of sex-linked colour is especially well documented across Poeciliidae (e.g. *Poecilia* and *Xiphophorus*) species (Lindholm and Breden 2002; McKinnon and Pierotti 2010).

**Table S2. Model parameters and settings used in the simulations.**

| Parameter | Notation | Value | Further information |
|---|---|---|---|
| Sex ratio of offspring and juveniles in the initial population: proportion of ZW or XY individuals. | *Aa0* | 0.5 | Corresponding to (Bókony et al. 2017): XY0, ZW0. |
| Number of adult females in the initial population. The number of adult males is set to the same value. | *NF0* | 100 | Corresponding to (Bókony et al. 2017): NF0. |
| Levels of genetic male signal produced by a single "a" or "A" allele on the sex-determining locus. | *sig_a* | 1 | Lacking empirical data and given that there is no consensus across similar models (Quinn et al. 2011; Schwanz et al. 2013), values of *sig_a* and *sig_A* were chosen arbitrarily. |
| | *sig_A* | 1.5 | |
| The initial sex determination system is defined by parameter thr0. If thr0×2 ≥ *sig_a*+*sig_A*, then the initial system is ZW/ZZ; otherwise, it is XX/XY. Furthermore, if *set_thr* is FALSE, thr0 defines the mean value of the normal distribution of which thr0 alleles will be chosen. | *thr0* | 1.375 | With the above settings of *sig_a* and *sig_A* this value defines a ZW/ZZ system. |
| | | 1.125 | With the above settings of *sig_a* and *sig_A* this value defines an XX/XY system. |
| Number of different threshold alleles present in the initial population. If *set_thr* is TRUE, the value of this paramater is ignored. | *n_thr0* | 2 | We followed (Schwanz et al. 2013) when we assumed that a single locus would define individual threshold for male signal. Number of alleles was set following empirical data by (Chandler et al. 2009; Wessels et al. 2014; Schroeder et al. 2016). Notably in reality this threshold may be influenced by multiple loci (Roff 1998; Holleley et al. 2016). |
| Vector defining the relative frequency of each threshold allele in the initial population, if set_thr_prob is TRUE. Relative frequency vaules must be given in the same order as allele values. | *thr_prob0* | c(0.5, 0.5) | Relative frequency vaules must be given in the same order as allele values. |
| Values of threshold alleles are defined by the vector def_thr0, if set_thr is TRUE. | *def_thr0* | c(1.31, 1.375) | Individual threshold for the level of male signal that is required for male development is the sum of the two threshold alleles carried. We set the lower threshold allele value so individual threshold of homozygotes would be just above the male signal value of genetically female individuals (following (Schwanz et al. 2013). The higher allele value was arbitrarily set to the mean of male signal values of genetic males and genetic females. With the above settings of *'sig_a'* and *'sig_A'*, |
| | | c(1.06, 1.125) | |

| | | | |
|---|---|---|---|
| | | | *def_thr0*=c(1.31, 1.375) refers to a ZW/ZZ system, while *def_thr0*=c(1.06, 1.125) refers to an XX/XY system. |
| Boolean defining if values of the threshold alleles are chosen manually (TRUE) or picked randomly from a normal distribution around *thr0* (FALSE). If TRUE, parameter *'def_thr0'* defines the threshold alleles. | *set_thr* | TRUE | In preliminary studies, we also explored simulations with randomly chosen thr allele values, but for simplicity, we finally decided to set them manually. |
| Boolean defining if relative frequencies of thr allele values are set manually in the initial population (i.e. as defined by thr_prob0). If FALSE, a frequency value will be chosen for each threshold allele from a normal distribution around 0.5, with values being forced to be between 0.25 and 0.75. (Note: this works only with exactly two thr alleles). | *set_thr_prob* | TRUE | In preliminary studies, we also explored simulations with randomly chosen initial thr allele frequencies, but for simplicity, we finally decided to set them manually. |
| Mean level of environmental male signal in the initial population. | *sig_env0* | 0 | Corresponding to (Bókony et al. 2017): m_masc0. |
| Extent of annual increase in environmental male signal, referring to climate warming. | *b_sig* | 0.003 | Similar, but not the same as (Bókony et al. 2017): b_masc. After 30 years of warming, probability of sex reversal is about 9% among genetically female individuals with the *'def_thr0'* values used here. We used this setting based on the findings of (Alho et al. 2010). |
| Standard deviation of the annual mean of environmental male signal, referring to annual stochasticity in climate warming. | *sd_sig* | 0.01 | Defines the error term $\varepsilon_B$ in the regression equation for calculating $sig_{env}$ (see Methods). Corresponding to (Bókony et al. 2017): sd_masc. |
| Standard deviation of the individual levels of environmental male signal, referring to individual differences in temperature experienced during the sensitive period of sexual development. | *sd_indivsig* | 0.05 | Defines the error term $\varepsilon_W$ in the regression equation for calculating $sig_{env}$ (see Methods). Individuals developing in different microhabitats in the same breeding site and/or at different times during the same breeding season may experience somewhat different temperatures. |
| Boolean defining whether aa individuals can become males or not. If TRUE, then all aa individuals will always become | *aa_no_masc* | FALSE | FALSE: corresponding to p_rel_WW_masc=1 in (Bókony et al. 2017), TRUE: corresponding to p_rel_WW_masc=0 in (Bókony et al. 2017). |

| | | | |
|---|---|---|---|
| females, regardless of environmental conditions. | | | In our simulations we assumed that aa (WW) individuals can sex-reverse. |
| Vector containing the preference alleles present in the initial population (values between 0 and 1 should be chosen, meaning the probability that the female would choose the normal male from a pool of one normal male and one sex-reversed male). Individual female preference is calculated as the average value of the two alleles carried by the female if *dom_pref* =FALSE, or as the value of the dominant allele if *dom_pref* =TRUE. | *def_pref0* | 0.9 | As a baseline scenario, we ran simulations where all the females preferred normal males. The allele value of 0.9 encodes 90% chance of choosing a normal male over a sex-reversed male. We did not set preference to 100% because females should mate even when only males of less desired quality are present (i.e. refusing reproduction is not beneficial). In the sensitivity analyses, we also ran scenarios assuming that females mate indiscriminately (allele value = 0.5). |
| | | c(0.1, 0.9) | In scenarios 10% $C_R$ and 90% $C_R$, we allowed two alleles on the C locus: one encoding 90% preference for normal males (or no preference, in sensitivity analyses) and the other 10% preference for normal males (i.e. 90% preference for sex-reversed males). |
| Vector containing the relative frequency of each preference allele in the initial population (given in the same order as allele values in *'def_pref0'*). | *pref_prob0* | 1 | This value was set when *'def_pref0'* was set to 0.9, resulting in uniform preference for normal males across all females. |
| | | c(0.1, 0.9) | In this scenario, the allele encoding preference for masculinized individuals is rare (10%). Recent studies suggest that sex reversal may occur naturally in some species (Alho et al. 2010; Perrin 2016; Lambert et al. 2019). Among such conditions, a certain level of variance could be maintained in female mating preferences prior to considerable climate warming. |
| | | c(0.9, 0.1) | In this scenario, the allele encoding preference for masculinized individuals is common (90%). A pre-existing sensory bias can be present in the population, e.g. due to some valuable food of similar colour to that appearing on the body of sex-reversed individuals (Ryan 1998; Rodd et al. 2002). This preference allele plays little if any role in mating as long as sex reversal is very rare, thus it is (nearly) neutral in the initial population. |
| Dominance of a specific preference allele (defined under parameter 'dominant'). If FALSE: type of inheritance of female preference is intermediate, if TRUE: one allele is fully dominant over the other(s). | *dom_pref* | TRUE | Due to the lack of empirical data, we assumed that the inheritance of preference alleles is either intermediate or dominant-recessive; in this paper we only show the results of the latter in detail. |

| | | | |
|---|---|---|---|
| Value of the dominant preference allele. This is used only if dom_pref is TRUE. | *dominant* | 0.1<br>0.9 | We assumed that the allele encoding 90% preference for sex-reversed males was dominant when rare but recessive when common. We did not study the other two combinations because a rare recessive allele is unlikely to have enough time to spread under rapid climate change while a common dominant allele has a very high chance of fixation. |
| Boolean defining if proportion of female offspring should be calculated for mothers with respect for their relative preference for sex-reversed or normal males. *(Note: this function works only if def_pref0 contains exactly two values!)* | calc_success | TRUE | This option enables the user to track the progeny sex ratio of females that prefer normal or sex-reversed males. This helps the interpretation of results but increases run time considerably. |
| Boolean defining if coefficient (D) of linkage disequilibrium should be calculated between the allele encoding relative preference for sex-reversed males and 1) the threshold allele of lower value, 2) chromosome A. Furthermore, proportion of this preference allele will be calculated among alleles inherited from 1) fathers and 2) mothers. *(Note: this function works only if exactly two preference alleles and two threshold alleles are set in the initial population!)* | calc_linkage | TRUE | Linkage disequilibrium (LD) may occur between loci linked to different chromosomes (that is, D values deviating from zero (Slatkin 2008). This option enables the user to track D values over time, which helps the interpretation of results but increases run time. |
| Maximum number of offspring that can survive density-dependent selection during the early developmental (larval) stage. | *N_max* | 2000 | Corresponding to (Bókony et al. 2017): N_max. |
| Survival rate following the larval stage until the end of the first winter; may depend on genotypic sex (AA, Aa or aa). | *phi_AA* | 0.3 | Corresponding to (Bókony et al. 2017): phi_ZZ. |
| | *phi_Aa* | 0.3 | Corresponding to (Bókony et al. 2017): phi_WZ and phi_XY. |
| | *phi_aa* | 0.3 | Corresponding to (Bókony et al. 2017): phi_WW and phi_XX. In most of our simulations, survival was independent of genotype. However, in sensitivity analyses, we ran scenarios with reduced WW survival (*phi_aa* = 0.225, 0.15, 0.075, or zero). |

| | | | |
|---|---|---|---|
| Annual survival rate of juveniles (i.e. from the end of the first winter until sexual maturity). Independent from genotype and phenotype. | *phi_juv* | 0.4 | Corresponding to (Bókony et al. 2017): phi_juv. |
| Annual survival of mature individuals; depends on phenotypic sex. | *phi_m* | 0.5 | Annual survival rate of phenotypic males. Corresponding to (Bókony et al. 2017): phi_m. |
| | *phi_f* | 0.5 | Annual survival rate of phenotypic females. Corresponding to (Bókony et al. 2017): phi_f. |
| Mean number of offspring annually produced by a female which can successfully develop into viable juveniles if there is no density-dependent selection. | *f_fert* | 200 | Corresponding to (Bókony et al. 2017): fert. |
| Fertility of aa or Aa males relative to AA males. If any of these numbers is <1, then the number of offspring produced each year is min(*N_max*; *Nmother* × *f_fert* × mean(*m_fert*), where *N_max* is the carrying capacity, *Nmother* is the number of females that found a mating partner, *f_fert* is the average number of offspring each female can recruit in the absence of density-dependence, and *m_fert* is the relative fertility of each male that engaged in mating compared to the fertility of normal males. | *m_fert_aa* | 1 | Corresponding to (Bókony et al. 2017): alpha_WW, alpha_XX. |
| | *m_fert_Aa* | 1 | Corresponding to (Bókony et al. 2017): alpha_WZ.<br>In our simulations we assumed that sex-reversed males are as fertile as normal males. Additionally, we ran simulations where sex-reversed males' fertility was 75% of normal males' (based on the meta-analysis of (Senior et al. 2012); these results are not shown in this paper. |
| Maximum number of clutches that a male can fertilize during each breeding season. | *libido* | 3 | A male can successfully mate with multiple females within the same breeding season. However, sperm quality and quantity is limited, and males can get exhausted after a few fertilizations (Hettyey et al. 2009). |
| Age of sexual maturity, measured in years. If set to 1, the selection specified in phi_juv will not affect these individuals. | *mat_m* | 2 | Age of sexual maturity in phenotypic males. Corresponding to (Bókony et al. 2017): mat_m. |
| | *mat_f* | 2 | Age of sexual maturity in phenotypic females. Corresponding to (Bókony et al. 2017): mat_f. |
| Maximum age (measured in years) that an individual can reach. | *lifespan* | 12 | Corresponding to (Bókony et al. 2017): lifespan. |
| Number of simulation runs with the same settings. | *n_runs* | 100 | Populations were affected by several stochastic factors and repeated simulations were therefore needed, but simulations ran for considerable time. |

| | | | |
|---|---|---|---|
| Maximum number of years to simulate after the burn-in period. Running will stop earlier if the population goes extinct. | *t_Max* | 400 | The simulation stops when one of the phenotypic sexes disappears from the population, but maximum 400 years after burn-in (our simulated populations never persisted that long). |
| Number of years which allows the initial population to reach a stable state without climate change (burn-in period). | *t_burn_in* | 50 | The burn-in period allows the population to reach a stable structure (age groups, sex chromosome genotypes, etc.) before climate warming starts. |
| Boolean determining if plots of the burn-in period are to be shown from the first run. Useful for specifying the optimal value of t_burn_in. | *show_burn_in* | TRUE | When testing new settings, it can be useful to inspect graphs of sex ratios, allele frequencies etc. in the initial population. |

**Table S3. Linear contrasts comparing the three scenarios in either XX/XY or ZW/ZZ initial sex-determination system.** All p-values in the table were adjusted by simultaneous Bonferroni correction.

| **Starting system** | **Variable** | **Scenarios[1]** | **p** | **Mean1** | **Mean2** | **Difference** | **SE** | **Difference in generations** |
|---|---|---|---|---|---|---|---|---|
| XX/XY | Year of extinction | 0% - 10% | 1 | 122.77 | 122.81 | 0.04 | 0.44 | 0.01 |
| | | **0% - 90%** | <0.0001 | 122.77 | 126.72 | 3.95 | 0.44 | 1.32 |
| | | **10% - 90%** | <0.0001 | 122.81 | 126.72 | 3.91 | 0.44 | 1.3 |
| | Length of XX/XY (years) | **0% - 10%** | <0.0001 | 36.88 | 30.87 | -6.01 | 0.84 | -2 |
| | | **0% - 90%** | <0.0001 | 36.88 | 28.49 | -8.39 | 0.84 | -2.8 |
| | | 10% - 90% | 0.164 | 30.87 | 28.49 | -2.38 | 0.84 | -0.79 |
| | Length of final period (years) | **0% - 10%** | <0.0001 | 85.89 | 91.94 | 6.05 | 0.98 | 2.02 |
| | | **0% - 90%** | <0.0001 | 85.89 | 98.23 | 12.34 | 0.98 | 4.11 |
| | | **10% - 90%** | <0.0001 | 91.94 | 98.23 | 6.29 | 0.98 | 2.1 |
| | Year when 5-yr average ASR first exceeded 0.6 | 0% - 10% | 1 | 53.90 | 54.70 | 0.80 | 1.08 | 0.27 |
| | | **0% - 90%** | 0.003 | 53.90 | 58.20 | 4.30 | 1.08 | 1.43 |
| | | **10% - 90%** | 0.042 | 54.70 | 58.20 | 3.50 | 1.08 | 1.17 |
| ZW/ZZ | Year of extinction | 0% - 10% | 1 | 294.26 | 293.35 | -0.91 | 0.72 | -0.3 |
| | | 0% - 90% | 0.090 | 294.26 | 292.07 | -2.19 | 0.72 | -0.73 |
| | | 10% - 90% | 1 | 293.35 | 292.07 | -1.28 | 0.72 | -0.43 |
| | Length of ZW/ZZ (years) | **0% - 10%** | 0.003 | 35.73 | 32.57 | -3.16 | 0.79 | -1.05 |
| | | **0% - 90%** | <0.0001 | 35.73 | 29.96 | -5.77 | 0.79 | -1.92 |
| | | **10% - 90%** | 0.038 | 32.57 | 29.96 | -2.61 | 0.79 | -0.87 |
| | Length of transition (years) | **0% - 10%** | <0.001 | 91.42 | 86.42 | -5.00 | 1.08 | -1.67 |
| | | **0% - 90%** | <0.0001 | 91.42 | 59.68 | -31.74 | 1.08 | -10.58 |
| | | **10% - 90%** | <0.0001 | 86.42 | 59.68 | -26.74 | 1.08 | -8.91 |
| | Length of XX/XY (years) | 0% - 10% | 1 | 101.22 | 99.63 | -1.59 | 0.98 | -0.53 |
| | | 0% - 90% | 1 | 101.22 | 99.27 | -1.95 | 0.98 | -0.65 |
| | | 10% - 90% | 1 | 99.63 | 99.27 | -0.36 | 0.98 | -0.12 |
| | Length of final period (years) | **0% - 10%** | <0.0001 | 65.89 | 74.73 | 8.84 | 1.25 | 2.95 |
| | | **0% - 90%** | <0.0001 | 65.89 | 103.16 | 37.27 | 1.25 | 12.42 |
| | | **10% - 90%** | <0.0001 | 74.73 | 103.16 | 28.43 | 1.25 | 9.48 |
| | Mean ASR during transition | **0% - 10%** | <0.0001 | 0.72 | 0.63 | -0.09 | 0.00 | - |
| | | **0% - 90%** | <0.0001 | 0.72 | 0.49 | -0.23 | 0.00 | - |
| | | **10% - 90%** | <0.0001 | 0.63 | 0.49 | -0.14 | 0.00 | - |
| | Year when 5-yr average ASR first exceeded 0.6 | **0% - 10%** | <0.001 | 53.22 | 61.20 | 7.98 | 1.76 | 2.66 |
| | | **0% - 90%** | <0.0001 | 53.22 | 212.38 | 159.16 | 1.76 | 53.05 |
| | | **10% - 90%** | <0.0001 | 61.20 | 212.38 | 151.18 | 1.76 | 50.39 |

[1] Scenarios: 0% $C_R$, 10% $C_R$ and 90% $C_R$ (referring to the proportion of allele $C_R$ in the initial population that encoded strong female preference for sex-reversed males).

### Formulas for calculating generation time, effective population size, sex-ratio selection andlinkage

Generation time (T; mean age of reproduction) was calculated following (Case 1999):

$$T = \frac{1}{R_0} \sum_{x=1}^{max} l_x b_x x \,,$$

where $R_0$ is the average number of female offspring produced by a female over her lifetime, $l_x$ is the annual survival rate and $b_x$ is the average number of daughters produced when she is x years old. In all our simulations, this yielded a generation time of 3 years.

Effective population size ($N_e$) is an important measure for conservation biology and it is strongly affected by ASR. Therefore, we estimated $N_e$ for each year in each run using the formula by (Hartl and Clark 2007: page 124) as

$$N_e = \frac{4N_m N_f}{N_m + N_f} ,$$

where $N_m$ is the number of adult males and $N_f$ is the number of adult females. In each run, we recorded the year when the ultimate population decline started, i.e. when $N_e$ permanently decreased below the early population equilibrium $N_e$, calculated as the average $N_e$ during 10 years before the start of climate warming when the population was in a stable state (i.e. last 10 years of the burn-in period).

In some simulations that we ran as sensitivity tests for our parameter settings, we explored the effects of reduced WW viability. Because in our model density-dependent selection occurred before genotype-dependent mortality, extra mortality of WW individuals caused unrealistically rapid decline in the adult population size. To compensate for this, we calculated $N_e$ based on a corrected adult population size that would be expected if WW mortality occurred during the zygote stage (i.e. before the density-dependent larval mortality). In these specific scenarios $N_e$ was calculated the same way as above, but $N_m$ and $N_f$ respectively were calculated as the number adult males and females saved from the simulations multiplied by the correction factor Cf:

$$Cf = \frac{phi \times \left(N - O_{WW}\right) \times \frac{O_{ZZ}}{O_{ZZ} + O_{ZW}} + phi \times \left(N - O_{WW}\right) \times \frac{O_{ZW}}{O_{ZZ} + O_{ZW}} + phi \times O_{WW}}{phi \times O_{ZZ} + phi \times O_{ZW} + phi \times \frac{phi\_aa}{phi} \times O_{WW}} ,$$

where $O_{ZZ}$, $O_{ZW}$, and $O_{WW}$ are the number of offspring by each genotype calculated by the model for 3 years (i.e. the average generation time) before the current year, N is their total number, phi is the first-year survival of ZZ and ZW offspring, and phi_aa is the first-year survival of WW offspring.

To better understand forces behind the changes of $C_R$ frequency, we calculated *s*, the selection coefficient resulting from sex-ratio selection. In each year in each run, we recorded progeny sex ratio of females expressing preference towards sex-reversed males and normal males, denoted by $x_R$ and $x_N$ respectively. In scenario 10% $C_R$ every female carrying allele $C_R$ preferred sex-reversed males (dominant $C_R$), while in scenario 90% $C_R$ only $C_RC_R$ homozygotes expressed such preference (recessive $C_R$). Selection coefficient *s* against $C_NC_N$ was calculated following the usual definition (e.g.Hartl and Clark 2007: page 226), i.e. as the difference of the relative fitnesses of the two homozygotes. $C_RC_R$ is taken as the reference genotype and the relative fitness of $C_NC_N$ compared to it is calculated as the number of descendants of an average $C_NC_N$ individual two generations later divided by that of an average $C_RC_R$ individual. The genotype in the generation in focus (referred to as generation *P*) that produces more advantageous progeny sex ratio relative to the population sex ratio (i.e. ASR) of the next generation (referred to as generation *F1*) will have more descendants two generations later, i.e. in generation *F2*. In our model the preference locus *C* affects progeny sex ratio directly only in females, and we neglected the smaller indirect effect in males originating from linkage disequilibrium with the threshold locus and supposed that $C_RC_R$ and $C_NC_N$ males have equal progeny sex ratios.

First the relative fitness of $C_NC_N$ *females* was calculated compared to $C_RC_R$ *females* originating from their different progeny sex ratios. We used the male per female ratio (i.e. the odds of being a male) in generation *F1* referred to as *z*, and calculated from ASR as ASR/(1-ASR). *z* gives the relative value of a daughter compared to a son as an average female of generation *F1* will leave z times as many offspring as an average male of the same generation. Let α be the average number of offspring of a male in the generation *F1*, so the average number of offspring of a female in *F1* is *z*α. Furthermore, in the calculation of the number of *F2* descendants we need the number of offspring of an average female in generation *P*, referred to as *β*, and supposed to be the same for $C_RC_R$ and $C_NC_N$ females (as the *C* locus affects the sex ratio of the offspring, but has no effect on their number). An average $C_NC_N$ female of generation *P* has $x_N\beta$ sons and $(1-x_N)\beta$ daughters in *F1*, and in turn the number of her *F2* descendants is as follows:

$$x_N\beta \cdot \alpha + (1-x_N)\beta \cdot z\alpha,$$

while the number of *F2* descendants of an average $C_RC_R$ female is:

$$x_R\beta \cdot \alpha + (1-x_R)\beta \cdot z\alpha.$$

The relative fitness of a $C_NC_N$ female in generation *P* (taking a $C_RC_R$ female as the reference) is:

$$\frac{x_N + (1-x_N)z}{x_R + (1-x_R)z}.$$

The selection coefficient against $C_NC_N$ among females will be denoted by *s'* to distinguish it from *s*, the selection coefficient measuring the strength of sex-ratio selection in the whole population, among both males and females. One can express *s'* in terms of the relative fitness of $C_NC_N$ females using the definition of the selection coefficient:

$$\frac{x_N + (1-x_N)z}{x_R + (1-x_R)z} = 1 - s'$$

As half of the alleles on the *C* locus in generation *F1* comes from generation *P* females and their distribution between the sexes is affected by the mothers' genotype on the *C* locus, while the other half comes from generation *P* males with negligible effect of the fathers' genotype on the *C* locus on offspring sex ratio, we calculate *s* as

$$s \approx \frac{s'}{2} = \frac{\dfrac{x_N + (1-x_N)z}{x_R + (1-x_R)z}}{2}.$$

The above selection coefficient s can predict allele frequency changes between generation *F1* and *F2* using the progeny sex ratios of the females in generation *P*. As we had overlapping generations, when calculating *s* for year *t*, we used the progeny sex ratios recorded in year *t-T*, where *T* is the generation time, but we used year t ASR to calculate *z*, the male per female ratio.

We calculated the coefficient (D) of linkage disequilibrium (LD; see (Slatkin 2008) to explore if and when linkage disequilibria occurred between allele $C_R$ and 1) chromosome *A* and 2) the $thr_{low}$ allele.

$$D = p_{C_R B} p_{C_N b} - p_{C_R b} p_{C_N B} ,$$

where $C_R$ and $C_N$ are the alleles occurring on the preference locus, B and b are the alleles occurring on another locus and p is the probability of co-occurrence of two specified alleles in a single gamete. Deviation of D from zero means linkage disequilibrium between the two loci. If D is positive, the two alleles occur together more frequently than expected in case of independent inheritance, while negative D means that co-occurrence is less frequent than expected. Note that the value of D can range between -0.25 and 0.25, but is typically closer to 0 if the allele frequencies are far from 0.5 (similarly to frequency of $C_R$ in our simulations).

### References


Aida T. 1921. On the inheritance of color in a fresh-water fish, *Aplocheilus latipes* Temmick and Schlegel, with special reference to sex-linked inheritance. Genetics [Internet] 6:554–573. Available from: http://www.pubmedcentral.nih.gov/articlerender.fcgi?artid=1200522&tool=pmcentrez&rendertype=abstract

Alho JS, Matsuba C, Merilä J. 2010. Sex reversal and primary sex ratios in the common frog (Rana temporaria). Mol. Ecol. [Internet] 19:1763–1773. Available from: http://doi.wiley.com/10.1111/j.1365-294X.2010.04607.x

Bókony V, Kövér S, Nemesházi E, Liker A, Székely T. 2017. Climate-driven shifts in adult sex ratios via sex reversals: the type of sex determination matters. Philos. Trans. R. Soc. B Biol. Sci. 372:20160325.

Case TJ. 1999. Demographic Relationships. In: Case TJ, editor. An illustrated guide to theoretical ecology. 1st ed. New York: Oxford University Press. p. 79–102.

Chandler CH, Phillips PC, Janzen FJ. 2009. The evolution of sex-determining mechanisms: lessons from temperature-sensitive mutations in sex determination genes in Caenorhabditis elegans. J. Evol. Biol. 22:192–200.

Edmunds JSG, McCarthy RA, Ramsdell JS. 2000. Permanent and functional male-to-female sex reversal in d-rR strain medaka *(Oryzias latipes)* following egg microinjection of o,p'-DDT. Environ. Health Perspect. 108:219–224.

Hagen RH, Scriber JM. 1989. Sex-linked diapause, color, and allozyme loci in Papilio glaucus: Linkage analysis and significance in a hybrid zone. J. Hered. 80:179–185.

Hartl DL, Clark AG. 2007. Principles of population genetics. fourth edi. Sunderland, Massachusetts: Sinauer Associates, Inc. Publishers

Hettyey A, Vági B, Hévizi G, Török J. 2009. Changes in sperm stores, ejaculate size, fertilization success, and sexual motivation over repeated matings in the common toad, *Bufo bufo* (Anura: Bufonidae). Biol. J. Linn. Soc. 96:361–371.

Holleley CE, Sarre SD, O'Meally D, Georges A. 2016. Sex reversal in reptiles: reproductive oddity or powerful driver of evolutionary change? Sex. Dev. 10:279–287.

Horth L. 2006. A sex-linked allele, autosomal modifiers and temperature-dependence appear to regulate melanism in male mosquitofish (Gambusia holbrooki). J. Exp. Biol. 209:4938–4945.

Kingston JI, Rosenthal GG, Ryan MJ. 2003. The role of sexual selection in maintaining a colour polymorphism in the pygmy swordtail, *Xiphophorus pygmaeus*. Anim. Behav. 65:735–743.

Lambert MR, Tran T, Kilian A, Ezaz T, Skelly DK. 2019. Molecular evidence for sex reversal in wild populations of green frogs *(Rana clamitans)*. PeerJ [Internet] 7:e6449. Available from: https://peerj.com/articles/6449

Lindholm A, Breden F. 2002. Sex chromosomes and sexual selection in poeciliid fishes. Am. Nat. 160:S214–S224.

Mallery Jr. CS, Carrillo MM. 2016. A case study of sex-linkage in *Python regius* (Serpentes: Boidae), with new insights into sex determination in Henophidia. Phyllomedusa 15:29–42.

McKinnon JS, Pierotti ME. 2010. Colour polymorphism and correlated characters: genetic mechanisms and evolution. Mol. Ecol. 19:5101–5125.

Miura I, Kitamoto H, Koizumi Y, Ogata M, Sasaki K. 2011. An X-linked body color gene of the frog Rana rugosa and its application to the molecular analysis of gonadal sex differentiation. Sex. Dev. 5:250–258.

Parnell NF, Streelman JT. 2013. Genetic interactions controlling sex and color establish the potential for sexual conflict in Lake Malawi cichlid fishes. Heredity (Edinb). [Internet] 110:239–246. Available from: http://www.pubmedcentral.nih.gov/articlerender.fcgi?artid=3668650&tool=pmcentrez&rendertype=abstract

Perrin N. 2016. Random sex determination: When developmental noise tips the sex balance. BioEssays 38:1218–1226.

Pierotti MER, Martín-Fernández JA, Seehausen O. 2009. Mapping individual variation in male mating preference space: multiple choice in a color polymorphic cichlid fish. Evolution (N. Y). 63:2372–2388.

Pryke SR. 2009. Sex chromosome linkage of mate preference and color signal maintains assortative mating between interbreeding finch morphs. Evolution (N. Y). 64:1301–1310.

Quinn AE, Sarre SD, Ezaz T, Marshall Graves J a, Georges A. 2011. Evolutionary transitions between mechanisms of sex determination in vertebrates. Biol. Lett. 7:443–448.

Rodd FH, Hughes KA, Grether GF, Baril CT. 2002. A possible non-sexual origin of mate preference: are male guppies mimicking fruit? Proc. R. Soc. B Biol. Sci. 269:475–481.

Roff DA. 1998. Evolution of threshold traits: the balance between directional selection, drift and mutation. Heredity (Edinb). 80:25–32.

Ryan MJ. 1998. Sexual selection, receiver biases, and the evolution of sex differences. Science (80-. ). 281:1999–2003.

Schroeder AL, Metzger KJ, Miller A, Rhen T. 2016. A novel candidate gene for temperature-dependent sex determination in the common snapping turtle. Genetics 203:557–571.

Schwanz LE, Ezaz T, Gruber B, Georges A. 2013. Novel evolutionary pathways of sex-determining mechanisms. J. Evol. Biol. 26:2544–2557.

Senior AM, Nat Lim J, Nakagawa S. 2012. The fitness consequences of environmental sex reversal in fish: a quantitative review. Biol. Rev. 87:900–911.

Slatkin M. 2008. Linkage disequilibrium: uderstanding the evolutionary past and mapping the medical future. Nat. Rev. Genet. [Internet] 9:477–485. Available from: https://www.ncbi.nlm.nih.gov/pmc/articles/PMC5124487/pdf/nihms-831771.pdf

Smith DAS, Gordon IJ, Traut W, Herren J, Collins S, Martins DJ, Saitoti K, Ireri P, Ffrench-Constant R. 2016. A neo-W chromosome in a tropical butterfly links colour pattern, male-killing, and speciation. Proc. R. Soc. B [Internet] 283:20160821. Available from: http://dx.doi.org/10.1098/rspb.2016.0821%5Cnhttp://rspb.royalsocietypublishing.org.%5Cnhttp://rspb.royalsocietypublishing.org/lookup/doi/10.1098/rspb.2016.0821

Wessels S, Sharifi RA, Luehmann LM, Rueangsri S, Krause I, Pach S, Hoerstgen-Schwark G, Knorr C. 2014. Allelic variant in the anti-Müllerian hormone gene leads to autosomal and temperature-dependent sex reversal in a selected nile tilapia line. PLoS One 9:e114341.

Wright AE, Darolti I, Bloch NI, Oostra V, Sandkam B, Buechel SD, Kolm N, Breden F, Vicoso B, Mank JE. 2017. Convergent recombination suppression suggests role of sexual selection in guppy sex chromosome formation. Nat. Commun. [Internet] 8:14251. Available from:

http://www.nature.com/doifinder/10.1038/ncomms14251
