## Supplementary material for "Evolutionary and demographic consequences of temperature-induced masculinization under climate warming: the effects of mate choice": SI2

#### Table of contents:

Figure S2. Intermediate inheritance with two strong preference alleles (CR=0.1, CN=0.9)

Figure S3. CR versus no preference (CR=0.1, CN=0.5; dominant/recessive)

Figure S4. CR versus no preference (CR=0.1, CN=0.5), intermediate inheritance

Figure S5. Multiallelic threshold

Figure S6. 75% WW survival

Figure S7. 50% WW survival

Figure S8. 25% WW survival

Figure S9. WW lethality

#### General information:

Figures in this document show the results of simulations that we ran to assess sensitivity of our results described in the main text for certain parameter settings. All figures show median values calculated from 100 simulations. Vertical dotted lines indicate the end of each sex-determination period:

- in simulations starting with XX/XY system: XX/XY, and final period
- if the initial system was ZW/ZZ: ZW/ZZ, transition, XX/XY, and final period

Dashed lines indicate the start of ultimate population decline.
For comparison with the main results, see Figure 1 (ASR, relative frequencies of genotype and threshold alleles) and Figure 5 (relative frequency of $C_R$ allele) in the main text.

**Figure S2. Intermediate inheritance with two strong preference alleles ($C_R$=0.1, $C_N$=0.9)**

In this situation, the expression of female preference is distributed in the starting populations as follows:

- 10% $C_R$ scenario: 1% prefer sex-reversed males, 18% show no preference, 81% prefer normal males
- 90% $C_R$ scenario: 81% prefer sex-reversed males, 18% show no preference, 1% prefer normal males

The results (ASR, relative frequencies of sex-chromosome genotypes and threshold alleles, timing of transitions and extinction) are very similar to the results shown in the main text (dominant/recessive inheritance), except for one difference: starting with a rare $C_R$ in an XX/XY system, $C_R$ does not disappear (although its frequency decreases steadily).

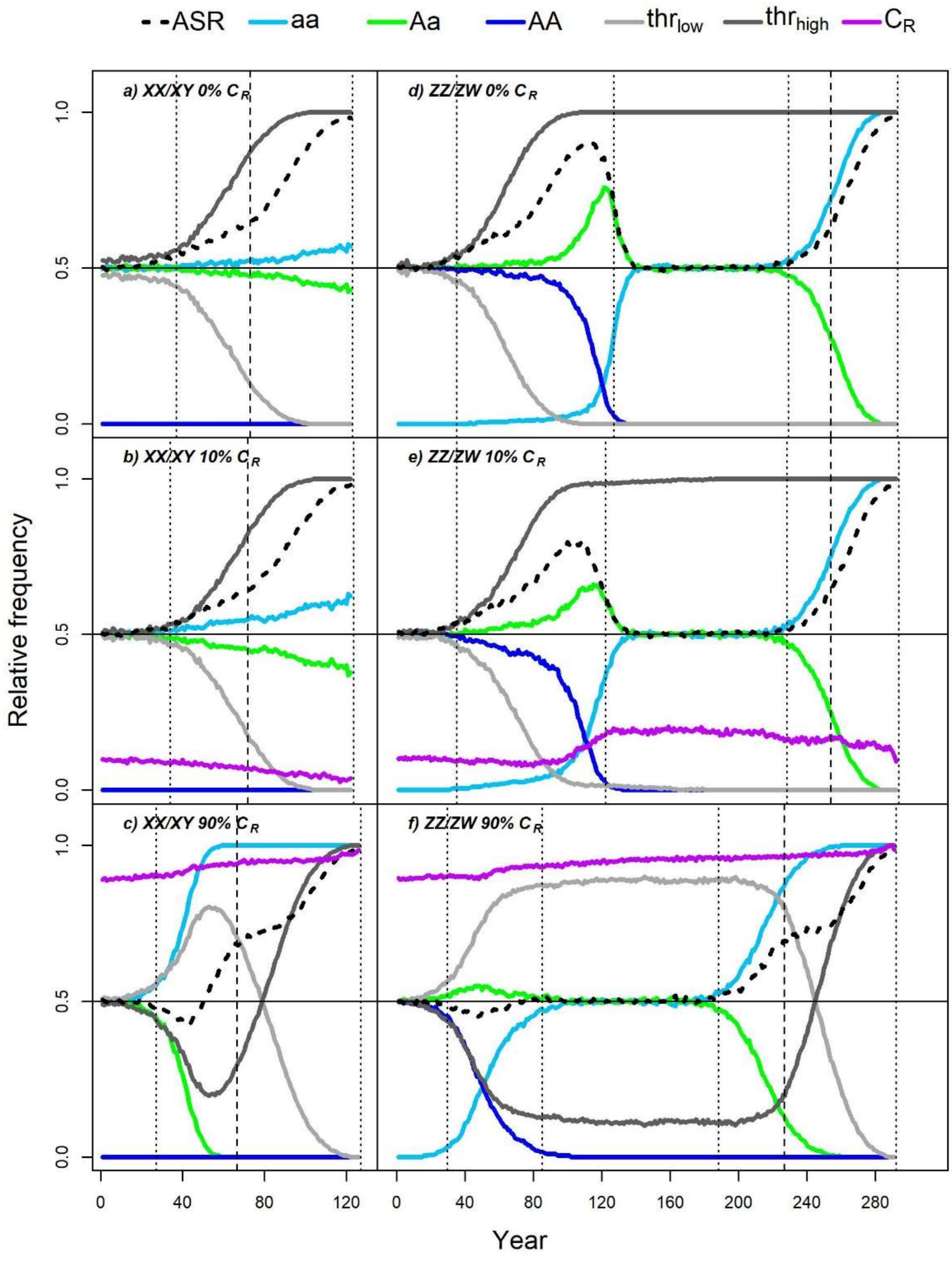

**Figure S3. $C_R$ versus no preference ($C_R$=0.1, $C_N$=0.5; dominant/recessive)**

In this situation, the expression of female preference is distributed in the starting populations as follows:

- 10% $C_R$ scenario: 19% prefer sex-reversed males, 81% show no preference
- 90% $C_R$ scenario: 81% prefer sex-reversed males, 19% show no preference

In these scenarios, the lack of preference for normal males results in faster extinction of the original male genotype, later spread of $thr_{high}$, and less marked ASR bias towards males during the ZZ/ZW-XX/XY transition compared to the scenarios in the main text. Also, the frequency of $C_R$ shows little change over time in the 10% $C_R$ scenarios (it is not selected against when rare because the majority of the females are indiscriminate). However, the effects of $C_R$ on ASR and transition times are very similar to the scenarios in the main text.

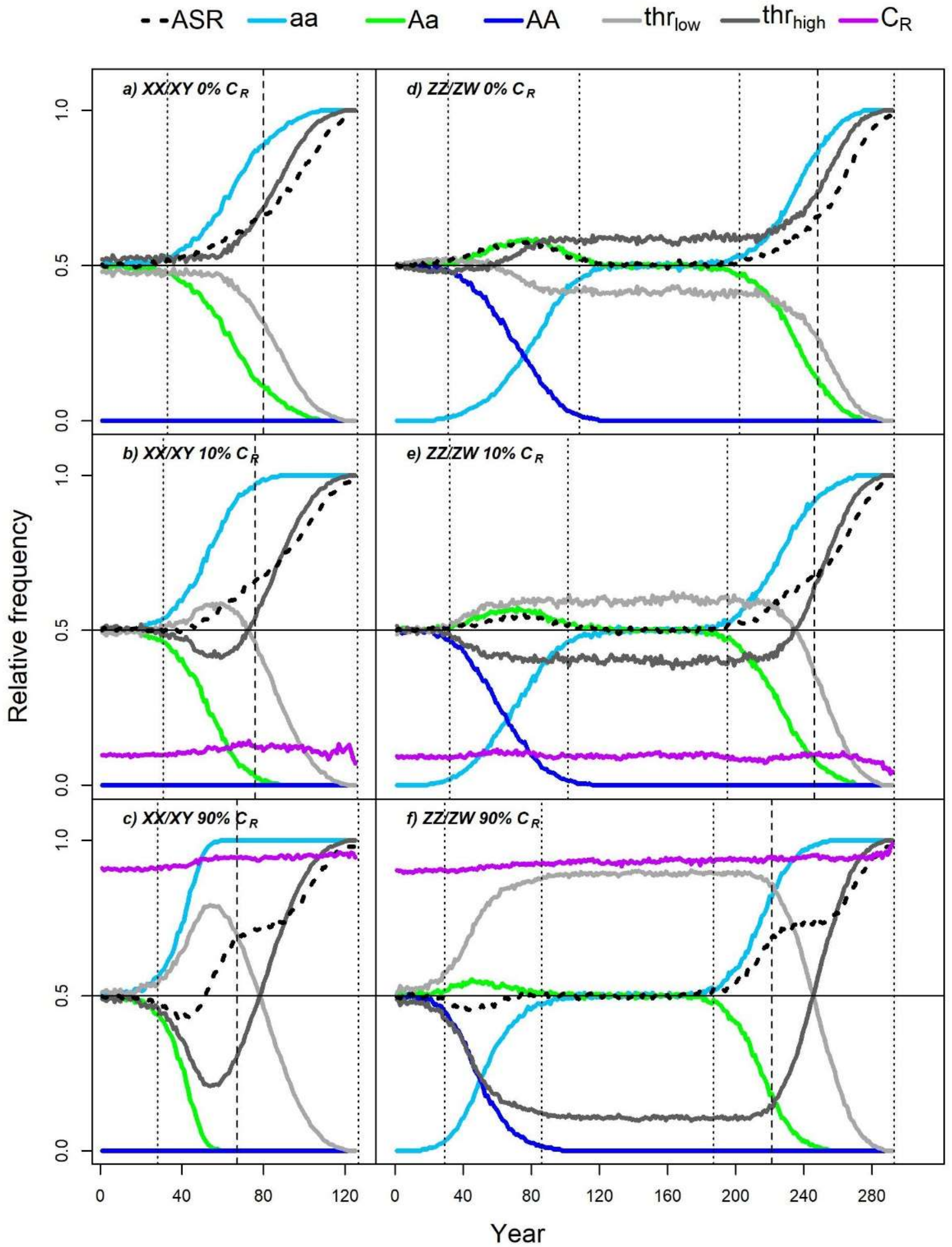

**Figure S4. $C_R$ versus no preference ($C_R$=0.1, $C_N$=0.5), intermediate inheritance**

In this situation, the expression of female preference is distributed in the starting populations as follows:

- 10% $C_R$ scenario: 1% prefer sex-reversed males with 0.9 probability, 18% prefer sex-reversed males with 0.7 probability, 81% show no preference
- 90% $C_R$ scenario: 81% prefer sex-reversed males with 0.9 probability, 18% prefer sex-reversed males with 0.7 probability, 1% show no preference

These results are very similar to the previous case ($C_R$ versus no preference; dominant/recessive). Thus, the frequency of $C_R$ changes very slightly or not at all when it is "competing" with a no-preference allele; this result does not depend on the type of inheritance. However, the effects of $C_R$ on ASR and transition times are very similar to the scenarios in the main text.

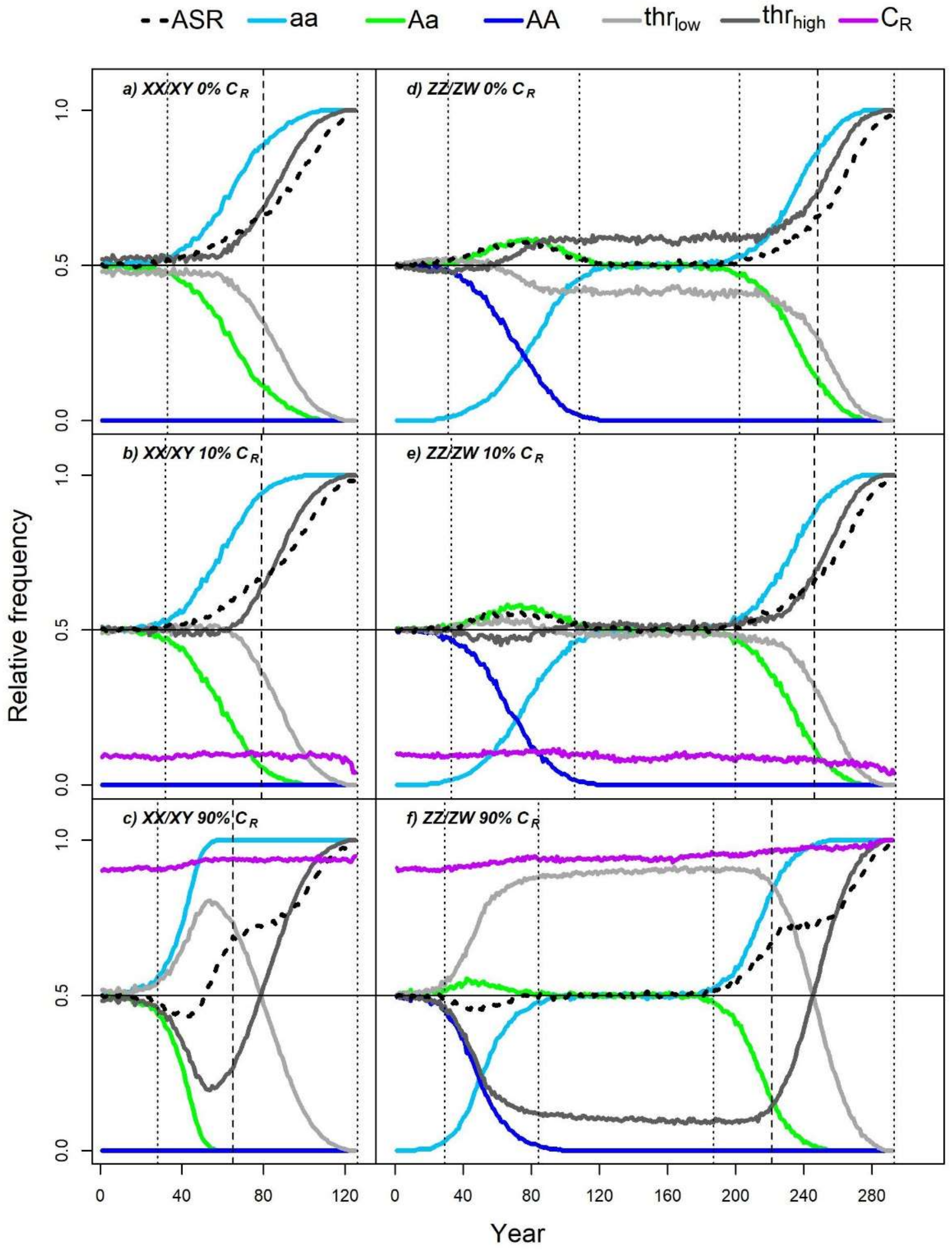

**Figure S5. Multiallelic threshold**

The number of *thr* alleles was set to 10, such that each allele started with a frequency of 0.1, and their values were evenly distributed between the lowest and highest value used in the main-text simulations.

The results are similar to those in the main text. Note: in the graphs below, solid grey curves show the frequency of the lowest and highest *thr* allele (from the original 10 alleles).

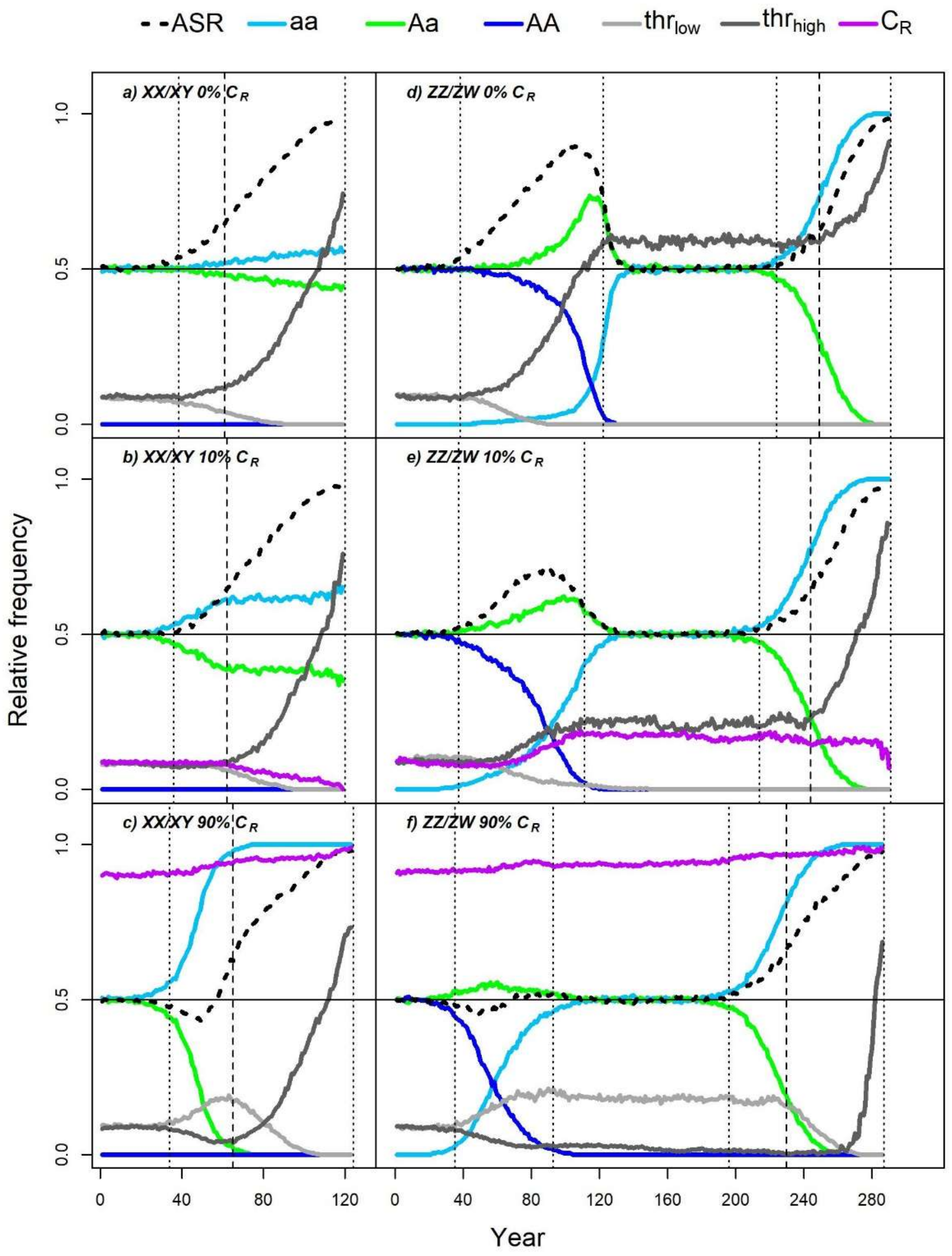

### Scenarios with reduced WW viability

**Figure S6. 75% WW survival**

When the survival of WW offspring is 75% of the survival of other genotypes, the spread of $C_R$ and its effects are similar to the scenarios with 100% WW viability: $C_R$ reduces the ASR bias and speeds up the transition. Because of the higher mortality of WW individuals (which are more likely to develop into phenotypic females), the newly evolved XX/XY system maintains a somewhat male-biased ASR until system destabilization.

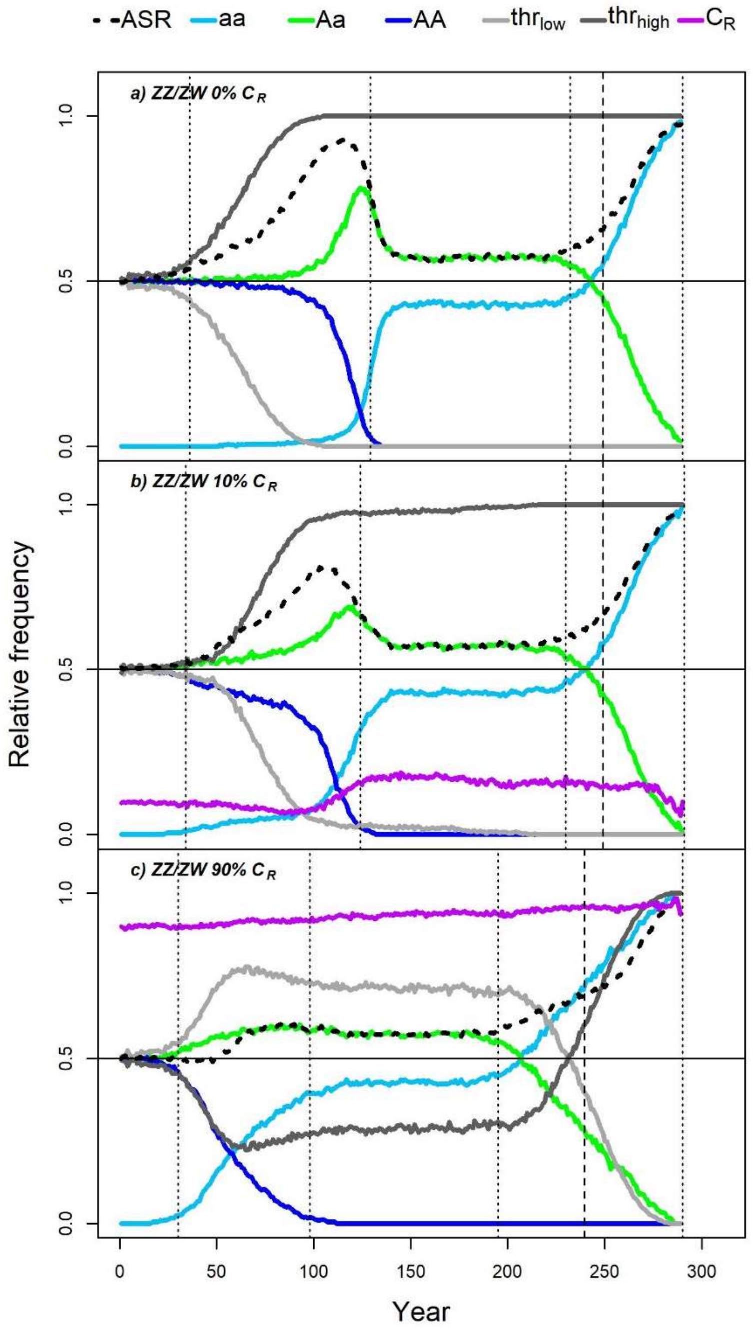

**Figure S7. 50% WW survival**

Again, the effects of $C_R$ are similar to the 100% WW viability scenarios. Relative frequency of the widespread $C_R$ allele increases only slightly, whereas the rare $C_R$ allele disappears, although it increases for a short time when ASR becomes highly male-biased before transition. After transition, ASR remains relatively male-biased.

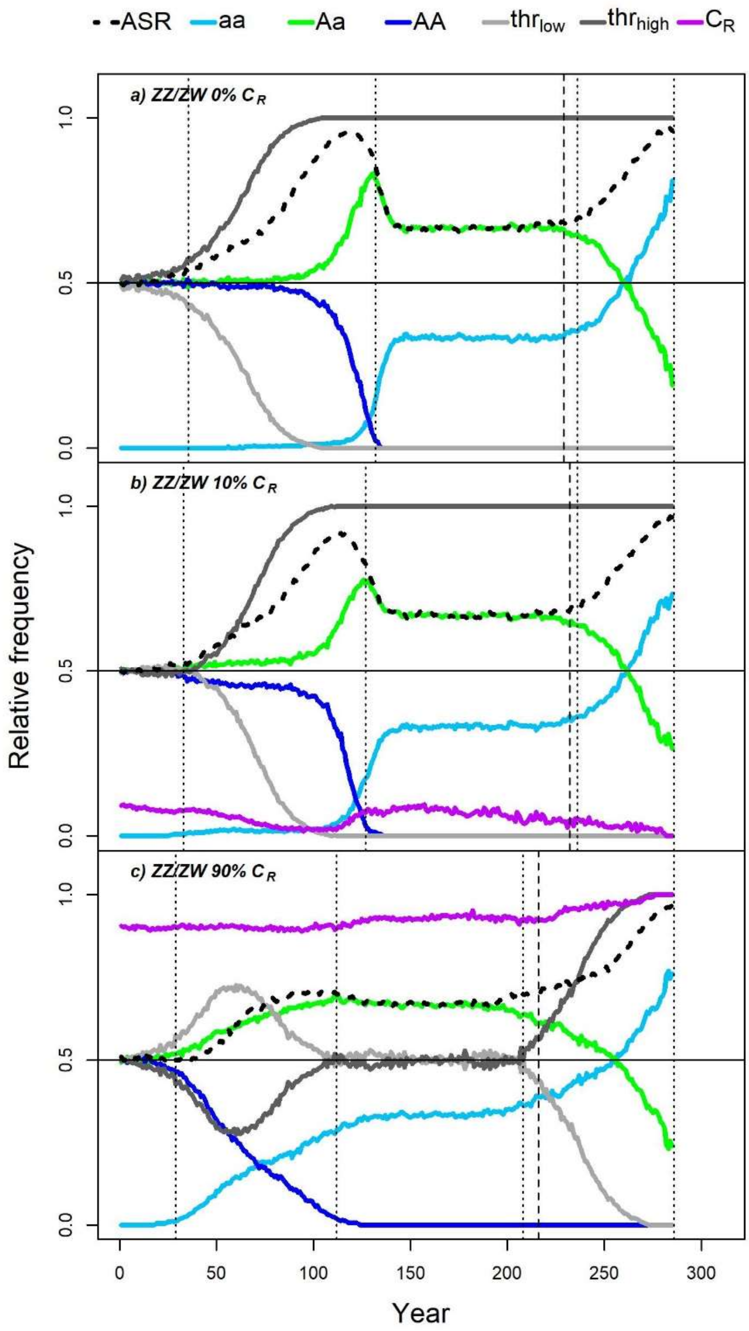

**Figure S8. 25% WW survival**

When the survival of WW offspring is only 25% of the survival of other genotypes, most of the populations cannot transition to an XX/XY system and die out after ca. 120 years when no $C_R$ allele is present. However, the presence of $C_R$ saves the population from this early extinction by enabling the switch to XX/XY system (in slightly more than half of the simulations when $C_R$ is rare; always when $C_R$ is widespread; see Fig. 6 in the main text). This switch happens rapidly when ASR has become highly male-biased; during this short time the relative frequency of $C_R$ increases. After transition, the population maintains a strongly male-biased ASR.

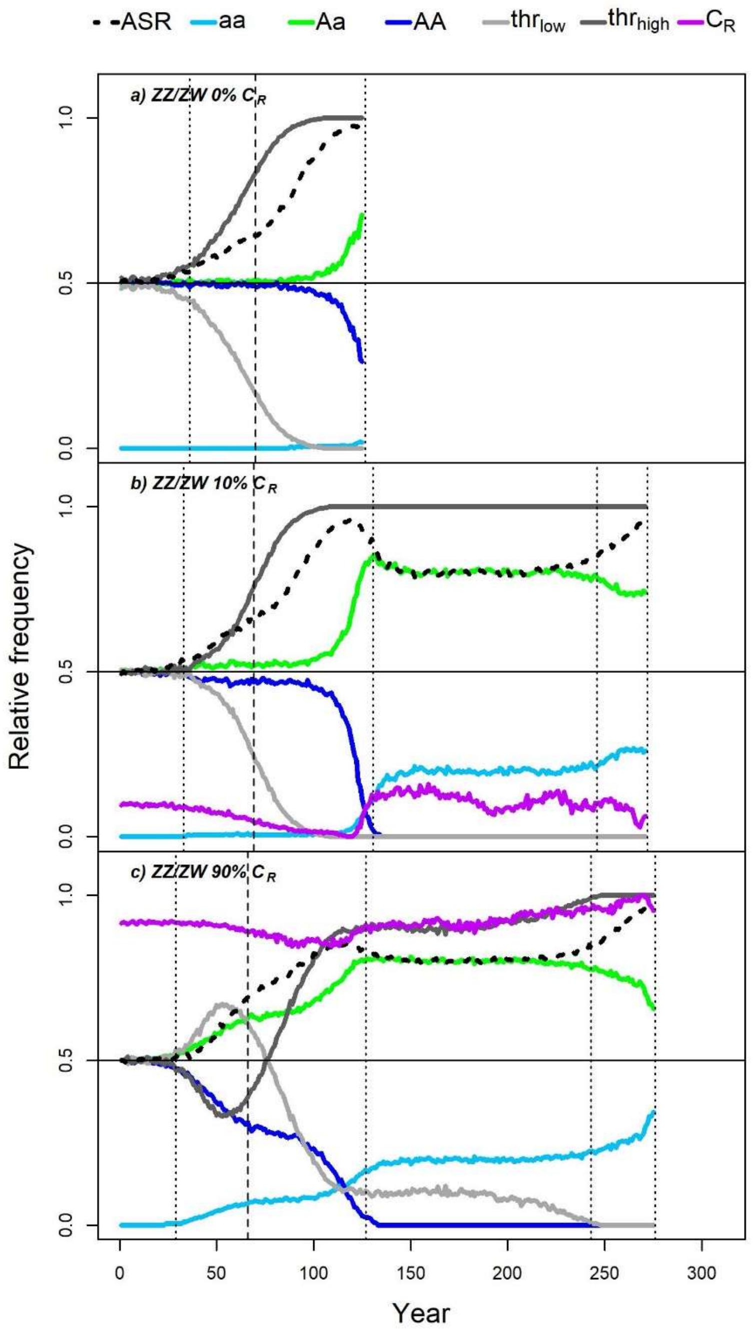

**Figure S9. WW lethality**

WW lethality makes the ZZ/ZW system behave similarly to the XX/XY system: no transition from ZZ/ZW to XX/XY, and the population dies out after ca. 120 years. The effects of $C_R$ are also similar: it has little effect on ASR change or extinction time, but it makes the starting system destabilize slightly earlier than when no $C_R$ is present. The only difference is that $C_R$ never spreads.

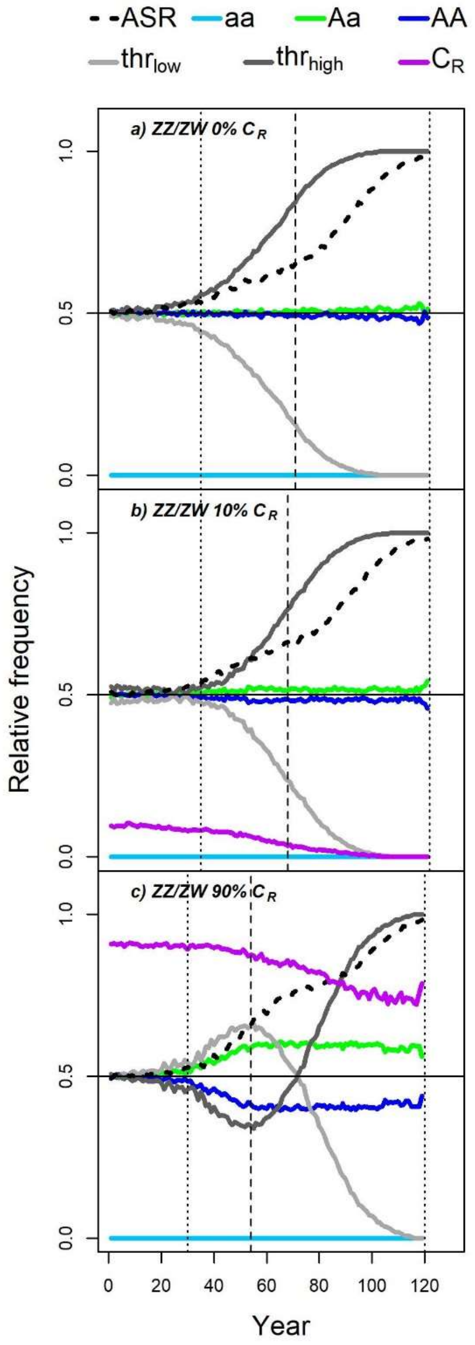
