## Supplementary material for "Evolutionary and demographic consequences of temperature-induced masculinization under climate warming: the effects of mate choice": SI3

#################
# Supporting Information 3 - Model code

# Evolutionary and demographic consequences of temperature-induced masculinization under climate warming: the effects of mate choice.
# Nemeshazi E1,2, Kover Sz3, Bokony V1
# 1 Lendulet Evolutionary Ecology Research Group, Plant Protection Institute, Centre for Agricultural Research, Herman Otto ut 15, 1022 Budapest, Hungary.
# 2 Konrad Lorenz Institute of Ethology Department of Interdisciplinary Life Sciences University of Veterinary Medicine, Savoyenstr. 1a, A-1160 Vienna, Austria
# 3 Conservation Genetics Research Group, Department of Ecology, University of Veterinary Medicine Budapest, Istvan utca 2, 1078 Budapest, Hungary


#################
#  Settings  #
#################

  ## Parameters referring to population size and sex chromosome frequencies in the initial population. During the burn-in period the concerned values will change according to several settings that refer to sexual development, selection and life history.
    # Aa0: relative frequency of genetic females among offspring and juveniles in the initial population. Default: 0.5.
    # NF0: number of adult females and adult males in the initial population. Default: 100.

  ## Parameters referring to sexual development. Default settings refer to a ZW/ZZ system
    # sig_a: level of genetic male signal produced by a single "a" allele on the sex determining gene. Default: 1.
    # sig_A: level of genetic male signal produced by a single "A" allele on the sex determining gene. Default: 1.5.
    # n_thr0: the number of different threshold alleles present in early population balance. If set_thr=TRUE, value of this paramater is ignored. Default: 2.
    # thr0: has two functions. 1) It defines the starting genetic system: if thr0*2 >= sig_a+sig_A, than we start with a ZW/ZZ system. Else, we startwith XX/XY. 
      # 2) If set_thr=FALSE, thr0 is the mean value of the normal distribution of which thr0 alleles will be chosen. Default: 1.375.
    # thr_prob0: if set_thr_prob=TRUE, thr_prob0 defines the relative frequency of each thr allele in the initial population. Relative frequency vaules must be given in the same order as allele values. Default: c(0.5, 0.5).
    # def_thr0: if set_thr=TRUE, def_thr0 defines the values of thr alleles. Default: c(1.31, 1.375).
    # set_thr: if set_thr is TRUE, parameter 'def_thr0' defines the values of thr0 alleles. Else, thr0 alleles are chosen randomly. Default: TRUE.
    # set_thr_prob: this boolean defines if relative frequencies of thr alleles are set manually in the initial population (i.e. as defined by thr_prob0). If set_thr_prob is FALSE, than relative frequencies will be assigned randomly to the thr alleles (this works exclusively with two thr alleles!).  Default: TRUE.
    # sig_env0: mean level of environmental male signal in early population balance. Default: 0.
    # b_sig: extent of annual increase in environmental male signal, referring to climate warming. Default: 0.003.
    # sd_sig: standard deviation of annual increase in environmental male signal, referring to annual stochasticity in average environmental temperature around the breeding season. Default: 0.01.
    # sd_indivsig: standard deviation of environmental male signal produced by an individual, referring to individual differnces in incubation temperature during the sensitive period of sexual development. Default: 0.05.
    # aa_no_masc: if this boolean is set to TRUE, individuals of genotype 'aa' will always become females, regardless of environmental temperature. Default: FALSE.
    
  ## Parameters determining breeding success of masculinized individuals.    
    # Relative fertility of masculinized individuals compared to AA males, regarding genotype. Default: 1 each.
      # m_fert_aa
      # m_fert_Aa
    # libido: maximum number of clutches a male can fertilize each breeding season. Default: 3.
    # Female preference in mate choice:
      # def_pref0: vector containing the preference alleles present in the initial population (values between 0 and 1 should be chosen). Individual female preference is calculated as A) average value of the two alleles carried by the female, or B) value of the dominant allele. Meaning of individual preference: probability that the female would choose the normal male from a pool of one normal male and one sex-reversed male. Default values: c(0.1, 0.9)
      # pref_prob0: relative frequency of each pref0 allele in the initial population (ranging from 0 to 1). Relative frequency vaules must be given in the same order as allele values. Default: c(0.5, 0.5).
      # dom_pref: if dom_pref= TRUE, type of inheritance of female preference is dominant/recessive. Else, it is intermediate. Default: FALSE.
      # dominant: value of the dominant allele (used if dom_pref=TRUE). Default: 0.1
      # calc_success: if this boolean is set to TRUE, proportion of female offspring will be calculated for females with respect for their relative preference for sex-reversed or normal males. NOTE: this function works only if def_pref0 contains exactly two values! Default: FALSE.
	  # calc_linkage: if this boolean is set to TRUE,  linkage disequilibrium will be calculated between the allele encoding relative preference for sex-reversed males and 1) the threshold allele of lower value, 2) chromosome A. Furthermore, proportion of this preference allele will be calculated among alleles inherited from 1) fathers and 2) mothers. NOTE: this function works only if exactly two preference alleles and two threshold alleles are set in the initial population! Default: FALSE.
    
  ## Parameters referring to selection regarding the environmental carrying capacity and annual survival.
    # N_max: maximum number of individuals which can undergo full metamorphosis. Refers to a density-dependent selection in the breeding lake. Default: 2000. Setting to a large number will considerably increase the running time!
    # Individual survival rate between leaving the natal lake and the end of the first winter is determined by genotype. Default: 0.3 for each genotype, but can be set to any number between 0 and 1.
      # phi_AA
      # phi_Aa
      # phi_aa
    # phi_juv: annual survival rate in juvenile stage. Independent from geno- or phenotype. Default: 0.4.
    # Annual survival rate of mature individuals depends on phenotypic sex.
      # phi_m: annual survival of adult males. Default: 0.5.
      # phi_f: annual survival of adult females. Default: 0.5.
      
  ## Parameters referring to life history.  
    # f_fert: mean number of offspring annually produced by a female which can leave the natal lake after metamorphism in a population of low density. Default: 200.
    # mat_m: age of sexual maturity in males (measured in years). If mat_m is set to 1, the selection specified in phi_juv will not affect males. Default: 2.
    # mat_f: age of sexual maturity in females (measured in years). If mat_f is set to 1, the selection specified in phi_juv will not affect females. Default: 2.
    # lifespan: maximum age that an individual can reach (measured in years). Default: 12.
    
  ## Parameters referring to further running detailes.
    # n_runs: number of independent runs of the same settings. Default: 1.
    # t_Max: maximum number of years to simulate after the burn-in period. Running will stop earlier if the population goes extinct. Default: 400.
    # t_burn_in: number of years without climate change. The burn-in period allows the population to reach a state of balance. Necessary time depends on settings of the initial population, selection and life-history. Default: 50.
    # show_burn_in: this boolean determines if plots of the burn-in period are to be shown from the first run. Useful to specify the optimal value of t_burn_in. Default: TRUE.

    
#############
#  Output  #
#############

  ## Output matrices
    # Returned data contain results from the burn-in period as well!
    # Within each matrix, a columns refer to runs and a rows refer to years.
    # Considered numbers and proportions are counted in the hypothetical autumn (after reproduction and before winter mortality).

    # Number of individuals, regarding age group and overall:
      # N_Adult: number of adults
      # N_Juvenile: number of juveniles
      # N_Offspring: number of offspring leaving the natal lake
      # Pop_Size: total population size

    # Portion of phenotypic males regarding age group and overall:
      # ASR: proportion of phenotypic males among adults
      # Male_Offspring: proportion of phenotypic males among offspring
      # Male_Juvenile: proportion of phenotypic males among juveniles
      # Male_All: overall proportion of phenotypic males across age groups

    # Proportion of AA and Aa genotypes regarding age group and overall, and chromosome A overall:
      # AA_Offspring: proportion of AA individuals among offspring
      # Aa_Offspring: proportion of Aa individuals among offspring    
      # AA_Juvenile: proportion of AA individuals among juveniles
      # Aa_Juvenile: proportion of Aa individuals among juveniles
      # AA_Adult_Male: proportion of AA individuals among adults males
      # Aa_Adult_Male: proportion of Aa individuals among adults males
      # aa_Adult_Male: proportion of aa individuals among adults males
      # AA_Adult_Female: proportion of AA individuals among adults females
      # Aa_Adult_Female: proportion of Aa individuals among adults females
      # aa_Adult_Female: proportion of aa individuals among adults females
      # AA_All: overall proportion of AA individuals across age groups      
      # Aa_All: overall proportion of Aa individuals across age groups
      # A_All: proportion of chromosome A across age groups

    # Proportion of sex-reversed individuals in certain groups
      # SR_Male: proportion of sex-reversed (masculinized) individuals among adult phenotypic males
      # SR_Female: proportion of sex-reversed (feminized) individuals among adult phenotypic females
      # SR_Father: proportion of sex-reversed (masculinized) individuals among fathers
      # Masc_rate: relative frequency of phenotypic males among individuals with female genotype(s) of the initial sex determination system (i.e. masculinization rate). When initial sex determination system is ZW/ZZ, then female genotypes are Aa and aa; but when we start with XX/XY, only aa is condisdered to be a female genotype.
    
    # Allele values and descriptive statistics of threshold alleles in certain groups
      # thr0: threshold allele values set in the initial population
      # Prop_thr0_Female: relative frequency of each thr0 allele among adult females
      # Prop_thr0_Male: relative frequency of each thr0 allele among adult males

    # Allele values and descriptive statistics of female choice (i.e. preference) alleles in certain groups
      # pref0: values of female choice alleles set in the initial population
      # Prop_pref_Fem: relative frequency of each female choice allele among adult females
      # Prop_pref_Male: relative frequency of each female choice allele among adult males (males only carry but express them)
      # Prop_daughter_CR: proportion of phenotypic females among the offspring of females with relative preference for sex-reversed males. These data will be saved only if calc_success is set to TRUE!
      # Prop_daughter_CN:  proportion of phenotypic females among the offspring of females with relative preference for normal males. These data will be saved only if calc_success is set to TRUE!
	  # LD_CR_lowthr: linkage disequilibrium between the allele encoding relative preference for sex-reversed males and the threshold allele of lower value. These data will be saved only if calc_linkage is set to TRUE!
	  # LD_CR_A: linkage disequilibrium between the allele encoding relative preference for sex-reversed males and chromosome A. These data will be saved only if calc_linkage is set to TRUE!
	  # Prop_father_CR: proportion of the allele encoding relative preference for sex-reversed males among preference alleles inherited from fathers. These data will be saved only if calc_linkage is set to TRUE!
	  # Prop_mother_CR: proportion of the allele encoding relative preference for sex-reversed males among preference alleles inherited from mothers. These data will be saved only if calc_linkage is set to TRUE!
	  
    
  ## Output vectors
    # Returned data refer to years calculated from the beginning of the "global warming period" (burn-in period not counted)
    # Within vectors, elements refer to runs.
    
    # t_ext_pop: the year of 'population extinction' (i.e. one of the sexes completely disappeared from the population -> reproduction is not possible). The burn-in period is excluded from this calculation!
    # t_ext_AA: the year when AA genotype first vanished from the population (it might have reappeared later). The burn-in period is excluded from this calculation!
    # t_ext_Aa: the year when Aa genotype first vanished from the population (it might have reappeared later). The burn-in period is excluded from this calculation!
    # t_ext_A: the year when chromosome A vanished from the population. The burn-in period is excluded from this calculation!
    
    
    

Sexsel_Sexrev_Aa = function(Aa0=0.5, NF0=100, sig_a=1, sig_A=1.5,
  n_thr0=2, thr0=1.375, thr_prob0=c(0.5, 0.5), def_thr0=c(1.31, 1.375), set_thr=TRUE, set_thr_prob=TRUE,
  sig_env0=0, b_sig=0.003, sd_sig=0.01, sd_indivsig=0.05, aa_no_masc=FALSE,
  def_pref0=c(0.1, 0.9), pref_prob0=c(0.5, 0.5), dom_pref=FALSE, dominant=0.1, calc_success=FALSE, calc_linkage=FALSE,
  N_max=2000, phi_AA=0.3, phi_Aa=0.3, phi_aa=0.3, phi_juv=0.4, phi_m=0.5, phi_f=0.5,
  f_fert=200, m_fert_aa=1, m_fert_Aa=1, libido=3, mat_m=2, mat_f=2, lifespan=12,
  n_runs=1, t_Max=400, t_burn_in=50, show_burn_in=TRUE) {
  
  tmax = t_Max + t_burn_in  # tmax is the maximum number of years simulated in a run, including burn-in
  
  if(set_thr==TRUE) {
    n_thr0 = length(def_thr0) # if thr0 alleles are set manually, n_thr0 is calculated from the number of alleles set
  }
  
  #############################################################################
  # Save memory for matrices and vectors containing information from each run #
  #############################################################################
  
  if (calc_success==TRUE & length(def_pref0)!=2) { # if calc_success is set to TRUE, the total number of preference alleles must be 2!
      print("Error: Number of preference alleles should be 2 or calc_success should be set to FALSE!")
      stop()
  }
  
  if (calc_linkage==TRUE & (length(def_pref0)!=2 | n_thr0!=2)) { # if calc_linkage is set to TRUE, the total number of preference alleles must be 2!
    print("Error: Number of both preference alleles and threshold alleles should be 2 or calc_linkage should be set to FALSE!")
    stop()
  }

  if (set_thr_prob==TRUE & length(thr_prob0)!=n_thr0) {
      print("Error: Length of vectors thr_prob0 and def_thr0 should be the same or set_thr_prob should be set to FALSE!")
      stop() 
  }
  
  if (length(pref_prob0)!=length(def_pref0)) {
      print("Error: Length of vectors pref_prob0 and def_pref0 should be the same!")
      stop() 
  }
  
  if (set_thr_prob==FALSE & n_thr0!=2) {  # The model currently can generate probabilities (between 0.25 and 0.75) only for 2 alleles!
      print("Error: n_thr0 (i.e. number of threshold alleles to generate) should be 2 or set_thr_prob should be set to TRUE and probabilities for n thr alleles should be set manually!")
      stop()
  }
  
  Male_matrix = matrix(NA, nrow=tmax+1, ncol=n_runs)  # proportion of phenotypic males among adults
  SR_in_Male_matrix = matrix(NA, nrow=tmax+1, ncol=n_runs)  # proportion of sex-reversed individuals among phenotypic adult males
  SR_in_Female_matrix = matrix(NA, nrow=tmax+1, ncol=n_runs)  # proportion of sex-reversed individuals among phenotypic adult females
  SR_in_Father_matrix = matrix(NA, nrow=tmax+1, ncol=n_runs)  # proportion of sex-reversed individuals among fathers
  Male_Offspring_matrix = matrix(NA, nrow=tmax+1, ncol=n_runs)  # proportion of males among offspring
  Male_Juvenile_matrix = matrix(NA, nrow=tmax+1, ncol=n_runs)  # proportion of males among juveniles
  Male_All_matrix = matrix(NA, nrow=tmax+1, ncol=n_runs)  # proportion of males across age groups
  Masc_rate_matrix = matrix(NA, nrow=tmax+1, ncol=n_runs)  # proportion of males among adults with female genotypes
  
  AA_Adult_matrix = matrix(NA, nrow=tmax+1, ncol=n_runs)    # relative frequency of AA genotype among adults
  Aa_Adult_matrix = matrix(NA, nrow=tmax+1, ncol=n_runs)    # relative frequency of Aa genotype among adults
  AA_Offspring_matrix = matrix(NA, nrow=tmax+1, ncol=n_runs)    # relative frequency of AA genotype among offspring
  Aa_Offspring_matrix = matrix(NA, nrow=tmax+1, ncol=n_runs)    # relative frequency of Aa genotype among offspring
  AA_Juvenile_matrix = matrix(NA, nrow=tmax+1, ncol=n_runs)    # relative frequency of AA genotype among juveniles
  Aa_Juvenile_matrix = matrix(NA, nrow=tmax+1, ncol=n_runs)      # relative frequency of Aa genotype among juveniles
  Aa_All_matrix = matrix(NA, nrow=tmax+1, ncol=n_runs)  # relative frequency of Aa genotype across age groups
  AA_All_matrix = matrix(NA, nrow=tmax+1, ncol=n_runs)  # relative frequency of AA genotype across age groups
  A_All_matrix = matrix(NA, nrow=tmax+1, ncol=n_runs)  # relative frequency of chromosome A across age groups
  
  AA_Adult_Male_matrix = matrix(NA, nrow=tmax+1, ncol=n_runs)  # proportion of of AA individuals among adult males
  Aa_Adult_Male_matrix = matrix(NA, nrow=tmax+1, ncol=n_runs)  # proportion of of Aa individuals among adult males
  aa_Adult_Male_matrix = matrix(NA, nrow=tmax+1, ncol=n_runs)  # proportion of of aa individuals among adult males

  AA_Adult_Female_matrix = matrix(NA, nrow=tmax+1, ncol=n_runs)  # proportion of of AA individuals among adult females
  Aa_Adult_Female_matrix = matrix(NA, nrow=tmax+1, ncol=n_runs)  # proportion of of Aa individuals among adult females
  aa_Adult_Female_matrix = matrix(NA, nrow=tmax+1, ncol=n_runs)  # proportion of of aa individuals among adult females
    

  Pop_Size_matrix = matrix(NA, nrow=tmax+1, ncol=n_runs)  # total population size including all age groups
  N_Juvenile_matrix = matrix(NA, nrow=tmax+1, ncol=n_runs)  # number of juveniles
  N_Offspring_matrix = matrix(NA, nrow=tmax+1, ncol=n_runs)  # number of offspring
  N_Adult_matrix = matrix(NA, nrow=tmax+1, ncol=n_runs)  # number of adults
    
  t_ext_pop_vector = numeric(n_runs)  # year of population extinction
  t_ext_A_vector = numeric(n_runs)  # year of extinction of A chromosome  
  t_ext_AA_vector = numeric(n_runs)  # year when AA genotype first vanished from the population
  t_ext_Aa_vector = numeric(n_runs)  # year when Aa genotype first vanished from the population

  thr0_matrix = matrix(NA, nrow=n_thr0, ncol=n_runs)  # list of thr alleles set before burn-in
  
  pref0_matrix = matrix(NA, nrow=length(def_pref0), ncol=n_runs)  # list of preference alleles set before burn-in
  Prop_pref_Female_matrix = matrix(NA, nrow=tmax+1, ncol=n_runs*length(def_pref0)) # relative frequency of each preference allele among adult females in each run
  Prop_pref_Male_matrix = matrix(NA, nrow=tmax+1, ncol=n_runs*length(def_pref0)) # relative frequency  of each preference allele among adult males in each run  
  
  Prop_thr0_Female_matrix = matrix(NA, nrow=tmax+1, ncol=n_runs*n_thr0)  # relative frequency of each thr allele among adult females, by-column following their by-row order in thr0_matrix
  Prop_thr0_Male_matrix = matrix(NA, nrow=tmax+1, ncol=n_runs*n_thr0)  # relative frequency of each thr allele among adult males, by-column following their by-row order in thr0_matrix

  Prop_daughter_CR_matrix = matrix(NA, nrow=tmax+1, ncol=n_runs)  # proportion of phenotypic females among the offspring of females with relative preference for sex-reversed males (i.e. mascuinized)
  Prop_daughter_CN_matrix = matrix(NA, nrow=tmax+1, ncol=n_runs)  # proportion of phenotypic females among the offspring of females with relative preference for normal males
  
  LD_CR_lowthr_matrix = matrix(NA, nrow=tmax+1, ncol=n_runs)  # linkage disequilibrium between the allele encoding relative preference for sex-reversed males and the threshold allele of lower value
  LD_CR_A_matrix = matrix(NA, nrow=tmax+1, ncol=n_runs)  # linkage disequilibrium between the allele encoding relative preference for sex-reversed males and chromosome A
  Prop_father_CR_matrix = matrix(NA, nrow=tmax+1, ncol=n_runs)  # proportion of the allele encoding relative preference for sex-reversed males among preference alleles inherited from fathers
  Prop_mother_CR_matrix = matrix(NA, nrow=tmax+1, ncol=n_runs)  # proportion of the allele encoding relative preference for sex-reversed males among preference alleles inherited from mothers
  
  # Function for sampling two elements from a vector of any length (if length=1, then both sampled elements will be the same)
  sample.ok = function(t=c(0,0), s=2, p=c(1,1)) {
    if (length(t)==1) return(rep(t,s))
    else return (sample(x=t, size=s, prob=p, replace=T))
  }
  
  for(k in 1:n_runs) {
  
    print(paste("n. of runs=",k))
    t_ext_AA_vector[k] = NA    # As long as the genotype is present in the population, this value remains NA. When the genotype first vanishes, the year will be recorded (excluding burn-in).
    t_ext_Aa_vector[k] = NA    
    t_ext_A_vector[k] = NA  # As long as the genotype chromosome A is present in the population, this value remains NA. When it becomes extinct the year will be recorded (excluding burn-in).

    #############################################################
    # Save memory for lists and vectors overwritten in each run #
    #############################################################

    # Number of individuals overall and regarding age groups
    N_Off = numeric(tmax+1)  # number of offspring that survived early density-dependent selection each year (i.e. in frogs: leaving the natal lake; counted before genotype-based selection)    
    N_Juven = numeric(tmax+1)  # number of juveniles (age between 1 year old and maturation age) in each year
    N_Indiv_All = numeric(tmax+1)  # full population size after reproduction in each year
    # Number of adults can be calculated as N_Female+N_Male
    
    # Number of individuals regarding phenotypic sex and age group
    N_Off_Male = numeric(tmax+1)     # number of phenotypic male offspring that survived early density-dependent selection, counted before genotype-based selection each year
    N_Off_Female = numeric(tmax+1)         
    N_Juv_Male = numeric(tmax+1)  # number of phenotypic male juveniles in each year
    N_Juv_Female = numeric(tmax+1)
    N_Male = numeric(tmax+1)    # number of adult phenotypic males in each year; N_Male = N_Male_aa + N_Male_Aa + N_Male_AA
    N_Female = numeric(tmax+1)    
    N_Male_All = numeric(tmax+1)  # number of males across age groups after reproduction in each year
    N_Female_All = numeric(tmax+1)

    # Number of individuals regarding sex, calculated before reproduction
    N_Male_JuvAd = numeric(tmax+1)  # total number of juveniles and adults with male phenotype in each year
    N_Female_JuvAd = numeric(tmax+1)      
    
    # Number of individuals regarding genotype and age group
    N_Offspring_aa = numeric(tmax+1) # number of aa offspring that survived early density-dependent selection, counted before genotype-based selection in each year
    N_Offspring_Aa = numeric(tmax+1) 
    N_Offspring_AA = numeric(tmax+1)
    N_Juv_aa = numeric(tmax+1)  # number of aa juveniles in each year
    N_Juv_Aa = numeric(tmax+1)  
    N_Juv_AA = numeric(tmax+1)
    N_Male_aa = numeric(tmax+1) # number of adult phenotypic males with genotype aa in each year
    N_Male_Aa = numeric(tmax+1)
    N_Male_AA = numeric(tmax+1)    
    N_Female_aa = numeric(tmax+1) # number of adult phenotypic females with genotype aa in each year
    N_Female_Aa = numeric(tmax+1)
    N_Female_AA = numeric(tmax+1)    

    # Frequency of each genotype among fathers in each year
    N_Father_aa = numeric(tmax+1) # number of aa males which became a father in each year
    N_Father_Aa = numeric(tmax+1)
    N_Father_AA = numeric(tmax+1)    
    
    # Vector containing threshold allele values
    thr0_vector = numeric(n_thr0)  # thr allele values present in the initial population
    N_thr_alleles_Offspring = numeric(n_thr0) # number of thr alleles among Offspring in each year
    N_thr_alleles_All = numeric(n_thr0) # total number of thr alleles in the population in each year
    
    # Statistics of the threshold alleles
    N_thr_alleles_Female = numeric(n_thr0)  # number of different thr alleles among adult females
    Min_thr_alleles_Female = numeric(n_thr0)  # minimum value of thr alleles among adult females
    Max_thr_alleles_Female = numeric(n_thr0)  # maximum value of thr alleles among adult females
    # Median_thr_alleles_Female = numeric(n_thr0)  # Median of thr alleles among adult females  
    N_thr_alleles_Male = numeric(n_thr0)  # number of different thr alleles among adult females
    Min_thr_alleles_Male = numeric(n_thr0)  # minimum value of thr alleles among adult females
    Max_thr_alleles_Male = numeric(n_thr0)  # maximum value of thr alleles among adult females
    # Median_thr_alleles_Male = numeric(n_thr0)  # Median of thr alleles among adult females        
    N_thr_alleles_Offspring = numeric(n_thr0)  # number of thr alleles present among Offspring
    N_thr_alleles_All = numeric(n_thr0)  # total number of thr alleles present in the population
    N_thr_alleles_Adult = numeric(n_thr0)  # total number of thr alleles present in adults
    
    # Vectors containging both copies of thr alleles carried by each individual in certain groups
    each_thr_offspring_long = numeric(N_max*2)
    each_thr_juvenile_long = numeric((N_max*max(mat_m, mat_f))*2)
    each_thr_female_long = numeric((lifespan-mat_f)*2)  # adult females
    each_thr_male_AA_long = numeric(2*N_max*(lifespan-mat_m))  # adult AA males
    each_thr_male_Aa_long = numeric(2*N_max*(lifespan-mat_m))
    each_thr_male_aa_long = numeric(2*N_max*(lifespan-mat_m))  

    # Vectors containging both copies of preference alleles carried by each individual in certain groups    
    each_pref_female_long = numeric((lifespan-mat_f)*2)  # adult females
    each_pref_male_aa_long = numeric((lifespan-mat_m)*2)  # adult aa males
    each_pref_male_Aa_long = numeric((lifespan-mat_m)*2)
    each_pref_male_AA_long = numeric((lifespan-mat_m)*2)
  
    prop_daughter_CR = numeric(tmax+1) # proportion of females among offspring of females with relative preference for sex-reversed males
    prop_daughter_CN = numeric(tmax+1) # proportion of females among offspring of females with relative preference for normal males
	
    ld_CR_lowthr = numeric(tmax+1)  # linkage disequilibrium between the allele encoding relative preference for sex-reversed males and the threshold allele of lower value
    ld_CR_A = numeric(tmax+1)  # linkage disequilibrium between the allele encoding relative preference for sex-reversed males and chromosome A
    prop_father_CR = numeric(tmax+1)  # proportion of the allele encoding relative preference for sex-reversed males among preference alleles inherited from fathers
    prop_mother_CR = numeric(tmax+1)  # proportion of the allele encoding relative preference for sex-reversed males among preference alleles inherited from mothers
      
    # Matrices containing relative frequency of each threshlod or preference allele among adult females and adult males, respectively. Each column refers to an allele.  
    pref_prop_fem_matrix = matrix(0, nrow=tmax+1, ncol=length(def_pref0)) # relative frequency of each preference allele among adult females in the current run
    pref_prop_male_matrix = matrix(0, nrow=tmax+1, ncol=length(def_pref0))    
    thr_prop_fem_matrix = matrix(0, nrow=tmax+1, ncol=n_thr0)  # relative frequency of each thr allele among adult females in the current run
    thr_prop_male_matrix = matrix(0, nrow=tmax+1, ncol=n_thr0)
        
    # Relative frequency of each genotype regarding age group
    aa_Offspring = numeric(tmax+1) # relative frequency of genotype aa among offspring that survived early density-dependent selection (similar to that in zygote stage)
    Aa_Offspring = numeric(tmax+1)
    AA_Offspring = numeric(tmax+1)    
    aa_Adult = numeric(tmax+1) # relative frequency of genotype aa among the adults in each year
    Aa_Adult = numeric(tmax+1)  
    AA_Adult = numeric(tmax+1)
    Aa_All = numeric(tmax+1) # overall proportion of genotype Aa in the population
    AA_All = numeric(tmax+1)
      
    # Relative frequency of each phenotypic sex in certain age groups      
    Male = numeric(tmax+1)   # proportion of all phenotypic males within the adults in each year
    Male_Offspring = numeric(tmax+1)  # proportion of Offspring with male phenotype
    Female_Offspring = numeric(tmax+1)
    
    # Proportion of sex-reversed individuals among certain groups
    SR_in_Male = numeric(tmax+1)      # proportion of masculinized individuals within all phenotypic males in each year
    SR_in_Female = numeric(tmax+1)      # proportion of feminized individuals within all phenotypic females in each year    
    SR_in_Father = numeric(tmax+1)      # proportion of masculinized individuals across all fathering events in each year
    Aa_in_Father = numeric(tmax+1)      # proportion of Aa masculinized individuals across all fathering events in each year
    aa_in_Father = numeric(tmax+1)      # proportion of aa masculinized individuals across all fathering events in each year

    # Relative frequency of each chromosome among adults regarding phenotypic sex, and in the population overall
    Prop_A_in_males = numeric(tmax+1)  # relative frequency of chromosome 'A' within phenotypic males in each year
    Prop_a_in_males = numeric(tmax+1)  # relative frequency of chromosome 'a' within phenotypic males in each year
    Prop_A_in_females = numeric(tmax+1)  # relative frequency of chromosome 'A' within phenotypic females in each year
    Prop_a_in_females = numeric(tmax+1)  # relative frequency of chromosome 'a' within phenotypic females in each year
    A_All = numeric(tmax+1) # overall relative frequency of chromosome 'A' in the population    
  
    # List of individuals in each age group 
    Offspring = vector("list",N_max)  # List of individuals that survived the early density-dependent selection (overwritten each year)
    Juvenile = vector("list",N_max*max(mat_m, mat_f)) # List of juvenile individuals (overwritten each year)
    Male_aa = vector("list",N_max*(lifespan-mat_m))  # List of aa adults with male phenotype (overwritten each year)
    Male_Aa = vector("list",N_max*(lifespan-mat_m))
    Male_AA = vector("list",N_max*(lifespan-mat_m))
    Female = vector("list",(lifespan-mat_f))  # List of adults with female phenotype: aa and Aa overall (overwritten each year)
    
    
    #################################
    # Create an initial population  #
    #################################
  
    if(set_thr==TRUE) {
      thr0_vector = def_thr0
    } else {
      thr0_vector = sort(c(rnorm(n_thr0, mean=thr0, sd=(sig_A-sig_a-4*(sd_indivsig+sd_sig))/4)))   # chose n_thr0 allele values, sort them in increasing order
    }
    
    if(set_thr_prob==TRUE) {
    
      thr_prob_vector = thr_prob0   # relative frequency of each thr allele in the initial population is set manually
      
    } else if (n_thr0==2) {  # set probability values for 2 thr alleles (forced to fall between 0.25 and 0.75)

      thr_prob1 = rnorm(1, mean=0.5, sd=0.15)    
      thr_prob1[thr_prob1<0.25] = 0.25
      thr_prob1[thr_prob1>0.75] = 0.75    
      thr_prob_vector = c(thr_prob1, 1-thr_prob1)
      
    }


    N = min(N_max,NF0*f_fert)  # N Offspring and Juvenile will be generated in the intial popoulation (N_max: environmental carrying capacity, 
                  # NF0: number of females (and males) in the initial population, f_fert: mean number of offspring annually produced by a female which could survive until the first winter if density is low.
                  # Number of juveniles in the initial population is arbitrarily set equal to the number of offspring.
        
    for(i in 1:(N_max*(lifespan-mat_m))) {    # save memory for parameters in each adult Male list
      Male_AA[[i]] = list(Chr=c("A","A"), Sex="m", Age=mat_m, thr=c(sample.ok(t=thr0_vector, p=thr_prob_vector)), pref=c(sample.ok(t=def_pref0, p=pref_prob0)))
      Male_Aa[[i]] = list(Chr=c("A","a"), Sex="m", Age=mat_m, thr=c(sample.ok(t=thr0_vector, p=thr_prob_vector)), pref=c(sample.ok(t=def_pref0, p=pref_prob0)))
      Male_aa[[i]] = list(Chr=c("a","a"), Sex="m", Age=mat_m, thr=c(sample.ok(t=thr0_vector, p=thr_prob_vector)), pref=c(sample.ok(t=def_pref0, p=pref_prob0)))
    }

    if(thr0*2 >= sig_a+sig_A) {  # if we start with a ZW/ZZ-type genetic sex determination system
      for(i in 1:(N_max*(lifespan-mat_f))) {   # save memory for parameters in the adult Female list
        Female[[i]] = list(Chr=c("A","a"), Sex="f", Age=mat_f, thr=c(sample.ok(t=thr0_vector, p=thr_prob_vector)), pref=c(sample.ok(t=def_pref0, p=pref_prob0)))
      }
      for(i in 1:(N_max*max(mat_m, mat_f))) { # save memory for parameters in the Juvenile list
        Juvenile[[i]] = list(Chr=c("A","A"), Sex="m", Age=1, thr=c(sample.ok(t=thr0_vector, p=thr_prob_vector)), pref=c(sample.ok(t=def_pref0, p=pref_prob0)))
      }
      for (i in 1:N_max) { # save memory for parameters in the Offspring list. Default: AA male.
        Offspring[[i]] = list(Chr=c("A","A"), Sex="m", Age=0, thr=c(sample.ok(t=thr0_vector, p=thr_prob_vector)), pref=c(sample.ok(t=def_pref0, p=pref_prob0)))
      }      
    }
    
    else {  # we start with an XX/XY-type genetic sex determination system
      for(i in 1:(N_max*(lifespan-mat_f))) { # save memory for parameters in the adult Female list
        Female[[i]] = list(Chr=c("a","a"), Sex="f", Age=mat_f, thr=c(sample.ok(t=thr0_vector, p=thr_prob_vector)), pref=c(sample.ok(t=def_pref0, p=pref_prob0)))
      }  
      for(i in 1:(N_max*max(mat_m, mat_f))) { # save memory for parameters in the Juvenile list
        Juvenile[[i]] = list(Chr=c("a","a"), Sex="f", Age=1, thr=c(sample.ok(t=thr0_vector, p=thr_prob_vector)), pref=c(sample.ok(t=def_pref0, p=pref_prob0)))
      }  
      for (i in 1:N_max) { # save memory for parameters in the Offspring list. Default: aa female.
        Offspring[[i]] = list(Chr=c("a","a"), Sex="f", Age=0, thr=c(sample.ok(t=thr0_vector, p=thr_prob_vector)), pref=c(sample.ok(t=def_pref0, p=pref_prob0)))
      }        
    }

    ###
    # Create N Offspring in the initial population 
    for (i in 1:N){ 
      if (runif(1)<Aa0){   # a proportion of offspring will be Aa
          if(thr0*2 >= sig_a+sig_A) {  # if we start with ZW/ZZ system
          Offspring[[i]]$Chr[sample(2,1)] = "a"
          if ( sig_a+sig_A+sig_env0 <= Offspring[[i]]$thr[1]+Offspring[[i]]$thr[2] ) {  # if total individual male signal does not exceed the individual threshold: change default sex to female
            Offspring[[i]]$Sex = "f"
          }
        } else {  # if we start with XX/XY system
          Offspring[[i]]$Chr[sample(2,1)] = "A"
          if ( sig_a+sig_A+sig_env0 > Offspring[[i]]$thr[1]+Offspring[[i]]$thr[2] ) {  # if total individual male signal does exceed the individual threshold: change default sex to male
            Offspring[[i]]$Sex = "m"
          }
        }

      } # if ends: some offspring are males, others are females
    } # for ends: initial Offspring list is done
          
  
    ###
    # Create N Juveniles in the initial population
    for(i in 1:N) {
      if(runif(1)<Aa0) { # a proportion of juveniles will be Aa

        if(thr0*2 >= sig_a+sig_A) {  # if we start with ZW/ZZ system
          Juvenile[[i]]$Chr[sample(2,1)] = "a"
          if ( sig_a+sig_A+sig_env0 <= Juvenile[[i]]$thr[1]+Juvenile[[i]]$thr[2] ) {  # if total individual male signal does not exceed the individual threshold: change default sex to 
            Juvenile[[i]]$Sex = "f"        
          }
        } else {  # if we start with XX/XY system
          Juvenile[[i]]$Chr[sample(2,1)] = "A"
          if ( sig_a+sig_A+sig_env0 > Juvenile[[i]]$thr[1]+Juvenile[[i]]$thr[2] ) {  # if total individual male signal does exceed the individual threshold: change default sex to male
            Juvenile[[i]]$Sex = "m"        
          }        
        }
      }
      
      # Age of each individual will be chosen randomly from an exponential distribution, with decimals being truncated. (min: 1 and max: age of maturation)
      if(Juvenile[[i]]$Sex=="f") {
        Juvenile[[i]]$Age = 1 + min(mat_f-1, trunc(rexp(1,1)))
      } else {
        Juvenile[[i]]$Age = 1 + min(mat_m-1, trunc(rexp(1,1)))
      }
    }  # for ends: initial Juvenile list is done
    
    ###
    # Create NF0 females (Aa) and the same number of AA males. Sex-reversal does not occur in the initial population!

    if(thr0*2 >= sig_a+sig_A) {  # if we start with ZW/ZZ system
      for(i in 1:NF0) {
        Male_AA[[i]]$Age = mat_m + min(lifespan-mat_m, trunc(rexp(1,1))) # Age of each individual will be chosen randomly from an exponential distribution, with decimals being truncated.(min: age of maturation, max: lifespan)
        Female[[i]]$Age = mat_f + min(lifespan-mat_f, trunc(rexp(1,1)))
      }
    } else { # else: we start with XX/XY system
      for(i in 1:NF0) {
        Male_Aa[[i]]$Age = mat_m + min(lifespan-mat_m, trunc(rexp(1,1)))
        Female[[i]]$Age = mat_f + min(lifespan-mat_f, trunc(rexp(1,1)))
      }
    } # initial lists of adult males and females are done.

    
    # Initial Population is done
    ############################
    

    t = 0  # t is for counting years in each run
    extinct = F # booleans to follow extinction of the population, genotypes and chromosome A
    extinct_AA = F
    extinct_Aa = F
    extinct_A = F

    
    #############################################
    # Simulated population changes across years #
    #############################################
    
    # We will follow changes year by year, as long as the population persists.

    while(t<=tmax & !extinct) {
      t = t+1  

      m_sig_env = ifelse(t<=t_burn_in, rnorm(1, mean=sig_env0, sd=sd_sig), rnorm(1, mean=(t-t_burn_in)*b_sig + sig_env0, sd=sd_sig)) # mean value of environmental male signal (increases with time)
        
      #########
      ### Survival of ADULTS from PREVIOUS YEAR
        # Selection based on phenotype!
        
      # ADULT MALES

      if(thr0*2>=sig_a+sig_A) {  # if we start with ZW/ZZ system
      
        N_Male_AA_Prev= ifelse(t==1, NF0, N_Male_AA[t-1])  # Temporary counter for AA males from previuos year. In the first year it is based on the initial population.
        N_Male_Aa_Prev = ifelse(t==1, 0, N_Male_Aa[t-1])  # Aa males from previuos year
        N_Male_aa_Prev = ifelse(t==1, 0, N_Male_aa[t-1])  # aa males from previuos year

      } else { # we start with XX/XY system
        N_Male_AA_Prev= ifelse(t==1, 0, N_Male_AA[t-1])  # Temporary counter for AA males from previuos year. In the first year it is based on the initial population.
        N_Male_Aa_Prev = ifelse(t==1, NF0, N_Male_Aa[t-1])  # Aa males from previuos year
        N_Male_aa_Prev = ifelse(t==1, 0, N_Male_aa[t-1])  # aa males from previuos year
      }

      # Adult AA males
      if (N_Male_AA_Prev>0) { # If there were adult AA males in the previous year, check each of them.
        for(i in 1:N_Male_AA_Prev) {
          rand = runif(1)
          if(rand<phi_m & Male_AA[[i]]$Age < lifespan) {  # if it survived the winter
            N_Male_AA[t] = N_Male_AA[t]+1  # count it as adult AA male in the CURRENT YEAR
            Male_AA[[N_Male_AA[t]]] = Male_AA[[i]]  # put it at the end of the current list of adult AA males
            Male_AA[[N_Male_AA[t]]]$Age = Male_AA[[N_Male_AA[t]]]$Age+1  # gets one year older

            each_pref_male_AA_long[(2*N_Male_AA[t])-1] = Male_AA[[N_Male_AA[t]]]$pref[1]  # put pref allele values of this individual in the pref allele vector of adult AA males
            each_pref_male_AA_long[2*N_Male_AA[t]] = Male_AA[[N_Male_AA[t]]]$pref[2]              
            each_thr_male_AA_long[(2*N_Male_AA[t])-1] = Male_AA[[N_Male_AA[t]]]$thr[1]  # put thr allele values  of this individual in the thr allele vector of adult AA males
            each_thr_male_AA_long[2*N_Male_AA[t]] = Male_AA[[N_Male_AA[t]]]$thr[2]                
          }
        } # for ends: CURRENT adult AA male list from surviving adults is done
      } # if ends: mature AA male survival done
      
      # Adult Aa males
      if (N_Male_Aa_Prev>0) {  # If there were any adult Aa males in the previous year, check each of them.
        for(i in 1:N_Male_Aa_Prev) {
          rand = runif(1)
          if(rand<phi_m & Male_Aa[[i]]$Age < lifespan) {  # if it survived the winter
            N_Male_Aa[t] = N_Male_Aa[t]+1  # count it as adult Aa male in the CURRENT YEAR
            Male_Aa[[N_Male_Aa[t]]] = Male_Aa[[i]]  # put it at the end of the current list of adult Aa males
            Male_Aa[[N_Male_Aa[t]]]$Age = Male_Aa[[N_Male_Aa[t]]]$Age+1  # gets one year older

            each_pref_male_Aa_long[(2*N_Male_Aa[t])-1] = Male_Aa[[N_Male_Aa[t]]]$pref[1]  # put pref allele values of this individual in the pref allele vector of adult Aa males
            each_pref_male_Aa_long[2*N_Male_Aa[t]] = Male_Aa[[N_Male_Aa[t]]]$pref[2]
            each_thr_male_Aa_long[(2*N_Male_Aa[t])-1] = Male_Aa[[N_Male_Aa[t]]]$thr[1]  # put thr allele values  of this individual in the thr allele vector of adult Aa males
            each_thr_male_Aa_long[2*N_Male_Aa[t]] = Male_Aa[[N_Male_Aa[t]]]$thr[2]
          }
        } # for ends: CURRENT adult Aa male list from surviving adults is done
      } # if ends: mature Aa male survival done
      
      # Adult aa males
      if (N_Male_aa_Prev>0) {  # If there were any adult aa males in the previous year, check each of them.
        for(i in 1:N_Male_aa_Prev) {
          rand = runif(1)
          if (rand<phi_m & Male_aa[[i]]$Age < lifespan) {  # if it survived the winter
            N_Male_aa[t] = N_Male_aa[t]+1 # count it as adult aa male in the CURRENT YEAR
            Male_aa[[N_Male_aa[t]]] = Male_aa[[i]] # put it at the end of the current list of adult aa males
            Male_aa[[N_Male_aa[t]]]$Age = Male_aa[[N_Male_aa[t]]]$Age+1 # gets one year older

            each_pref_male_aa_long[(2*N_Male_aa[t])-1] = Male_aa[[N_Male_aa[t]]]$pref[1]  # put pref allele values of this individual in the pref allele vector of adult aa males
            each_pref_male_aa_long[2*N_Male_aa[t]] = Male_aa[[N_Male_aa[t]]]$pref[2]
            each_thr_male_aa_long[(2*N_Male_aa[t])-1] = Male_aa[[N_Male_aa[t]]]$thr[1]  # put thr allele values  of this individual in the thr allele vector of adult aa males
            each_thr_male_aa_long[2*N_Male_aa[t]] = Male_aa[[N_Male_aa[t]]]$thr[2]              
          }
        } # for ends: CURRENT adult aa male list from surviving adults is done
      } # if ends: mature aa male survival done

      # FEMALES
      # Aa and aa are listed together, but are counted together and separately as well.
      N_Female_Prev = ifelse(t==1, NF0, N_Female[t-1])  # Temporary counter for females from previuos year. In the first year it is based on the initial population.
    
      if (N_Female_Prev>0) { # if there were any mature females in the previous year, check each of them.
        for(i in 1:N_Female_Prev) {
          if (runif(1)<phi_f & Female[[i]]$Age < lifespan) {  # if it survived the winter
            N_Female[t] = N_Female[t]+1  # count it as adult female in the CURRENT YEAR
            Female[[N_Female[t]]] = Female[[i]] # put it at the end of the current list of adult females
            Female [[N_Female[t]]]$Age = Female [[N_Female[t]]]$Age+1 # gets one year older

            each_pref_female_long[(2*N_Female[t])-1] = Female[[N_Female[t]]]$pref[1]  # put pref allele values of this individual in the pref allele vector of adult females
            each_pref_female_long[2*N_Female[t]] = Female[[N_Female[t]]]$pref[2]              
            each_thr_female_long[(2*N_Female[t])-1] = Female[[N_Female[t]]]$thr[1]  # put thr allele values  of this individual in the thr allele vector of adult females
            each_thr_female_long[2*N_Female[t]] = Female[[N_Female[t]]]$thr[2]
              
            if (Female[[i]]$Chr[1]!=Female[[i]]$Chr[2]) {  # if it is Aa female
              N_Female_Aa[t] = N_Female_Aa[t]+1  # count it as adult Aa female in the CURRENT YEAR        
            } else if(Female[[i]]$Chr[1]=="A" & Female[[i]]$Chr[2]=="A") {  # AA female
              N_Female_AA[t] = N_Female_AA[t]+1
            } else {  # aa female
              N_Female_aa[t] = N_Female_aa[t]+1
            }
          } # if ends: all surviving females has been counted and aged
        } # for ends: CURRENT adult female list from surviving adults is done
      } # if ends: mature female survival done

      
      #########
      ### Survival and maturation of JUVENILES from PREVIOUS YEAR
        # Selection is independent of geno- or phenotype!
      
      N_Juven_Prev = ifelse(t==1, N, N_Juven[t-1])  # Temporary counter for juveniles from previuos year. In the first year it is based on the initial population.
      if (N_Juven_Prev>0) { # if there are any juveniles to deal with
        for(i in 1:N_Juven_Prev) {
          rand=runif(1)
          
          if (rand<phi_juv) {  # if it survived the winter
          
            Juvenile[[i]]$Age =Juvenile[[i]]$Age+1  # gets one year older
                
            if (Juvenile[[i]]$Sex=="m") {  # MALE 
            
              if (Juvenile[[i]]$Age<=mat_m) {  # IMMATURE MALE
                  N_Juven[t] = N_Juven[t]+1  # count it as a juvenile in the CURRENT YEAR
                  N_Juv_Male[t] = N_Juv_Male[t]+1  # count it as a juvenile male
                  
                  Juvenile[[N_Juven[t]]] = Juvenile[[i]]  # put it to the end of the actual Juvenile list
                  
                  if(Juvenile[[i]]$Chr[1]=="A" & Juvenile[[i]]$Chr[2]=="A") {
                    N_Juv_AA[t] = N_Juv_AA[t]+1  # count it as AA juvenile in the CURRENT YEAR
                  } else if(Juvenile[[i]]$Chr[1]=="a" & Juvenile[[i]]$Chr[2]=="a") {
                    N_Juv_aa[t] = N_Juv_aa[t]+1    # count it as aa juvenile in the CURRENT YEAR
                  } else if(Juvenile[[i]]$Chr[1]!=Juvenile[[i]]$Chr[2]) {
                    N_Juv_Aa[t] = N_Juv_Aa[t]+1  # count it as Aa juvenile in the CURRENT YEAR
                  }

                  each_thr_juvenile_long[(N_Juven[t]*2)-1] = Juvenile[[N_Juven[t]]]$thr[1]  # put thr allele values of this individual in the thr allele vector of juveniles
                  each_thr_juvenile_long[N_Juven[t]*2] = Juvenile[[N_Juven[t]]]$thr[2]
          
              } else {  # MALE MATURING by the current breeding period

                  if (Juvenile[[i]]$Chr[1]=="A" & Juvenile[[i]]$Chr[2]=="A") {  # AA male
          
                    N_Male_AA[t] = N_Male_AA[t]+1    # count it as adult AA male in the CURRENT YEAR
                    Male_AA[[N_Male_AA[t]]] = Juvenile[[i]]    # put it at the end of the current list of adult AA males
                    each_thr_male_AA_long[(2*N_Male_AA[t])-1] = Male_AA[[N_Male_AA[t]]]$thr[1]  # put pref allele values of this individual in the pref allele vector of adult AA males
                    each_thr_male_AA_long[2*N_Male_AA[t]] = Male_AA[[N_Male_AA[t]]]$thr[2]  
                    each_pref_male_AA_long[(2*N_Male_AA[t])-1] = Male_AA[[N_Male_AA[t]]]$pref[1]  # put thr allele values of this individual in the thr allele vector of adult AA males
                    each_pref_male_AA_long[2*N_Male_AA[t]] = Male_AA[[N_Male_AA[t]]]$pref[2]
          
                  } else if (Juvenile[[i]]$Chr[1]!=Juvenile[[i]]$Chr[2]) {  # Aa male
          
                    N_Male_Aa[t] = N_Male_Aa[t]+1    # count it as adult Aa male in the CURRENT YEAR
                    Male_Aa[[N_Male_Aa[t]]] = Juvenile[[i]]    # put it at the end of the current list of adult Aa males
                    each_thr_male_Aa_long[(2*N_Male_Aa[t])-1] = Male_Aa[[N_Male_Aa[t]]]$thr[1]  # put pref allele values of this individual in the pref allele vector of adult Aa males
                    each_thr_male_Aa_long[2*N_Male_Aa[t]] = Male_Aa[[N_Male_Aa[t]]]$thr[2]
                    each_pref_male_Aa_long[(2*N_Male_Aa[t])-1] = Male_Aa[[N_Male_Aa[t]]]$pref[1]  # put thr allele values of this individual in the thr allele vector of adult Aa males
                    each_pref_male_Aa_long[2*N_Male_Aa[t]] = Male_Aa[[N_Male_Aa[t]]]$pref[2]
          
                  } else if (Juvenile[[i]]$Chr[1]=="a" & Juvenile[[i]]$Chr[2]=="a") {  # aa male
          
                    N_Male_aa[t] = N_Male_aa[t]+1    # count it as adult aa male in the CURRENT YEAR
                    Male_aa[[N_Male_aa[t]]] = Juvenile[[i]]  # put it at the end of the current list of adult aa males
                    each_thr_male_aa_long[(2*N_Male_aa[t])-1] = Male_aa[[N_Male_aa[t]]]$thr[1]  # put pref allele values of this individual in the pref allele vector of adult aa males
                    each_thr_male_aa_long[2*N_Male_aa[t]] = Male_aa[[N_Male_aa[t]]]$thr[2]
                    each_pref_male_aa_long[(2*N_Male_aa[t])-1] = Male_aa[[N_Male_aa[t]]]$pref[1]  # put thr allele values of this individual in the thr allele vector of adult aa males
                    each_pref_male_aa_long[2*N_Male_aa[t]] = Male_aa[[N_Male_aa[t]]]$pref[2]
                  }
                } # if ends: the male survived the last winter
                
            } else {  # FEMALE   
            
              if (Juvenile[[i]]$Age<=mat_f) {  # IMMATURE FEMALE
        
                  N_Juven[t] = N_Juven[t]+1  # count it as a juvenile
                  N_Juv_Female[t] = N_Juv_Female[t]+1  # count it as a juvenile female in the CURRENT YEAR
                      
                  Juvenile[[N_Juven[t]]] = Juvenile[[i]]  # put it to the end of the actual Juvenile list
                      
                  if (Juvenile[[i]]$Chr[1]=="a" & Juvenile[[i]]$Chr[2]=="a") {  # aa juvenile in the CURRENT YEAR
                    N_Juv_aa[t] = N_Juv_aa[t]+1
                  } else if (Juvenile[[i]]$Chr[1]!=Juvenile[[i]]$Chr[2]) {  # Aa juvenile in the CURRENT YEAR
                    N_Juv_Aa[t] = N_Juv_Aa[t]+1
                  } else {  # AA juvenile in the CURRENT YEAR
                    N_Juv_AA[t] = N_Juv_AA[t]+1
                  }

                  each_thr_juvenile_long[(N_Juven[t]*2)-1] = Juvenile[[N_Juven[t]]]$thr[1]  # put thr allele values of this individual in the thr allele vector of juveniles
                  each_thr_juvenile_long[N_Juven[t]*2] = Juvenile[[N_Juven[t]]]$thr[2]
          
              } else {  # FEMALE MATURING by the current breeding period
        
                  N_Female[t] = N_Female[t]+1  # count it as an adult female in the CURRENT YEAR
                  Female[[N_Female[t]]] = Juvenile[[i]]  # put it at the end of the current list of adult females
                  
                  if (Juvenile[[i]]$Chr[1]!=Juvenile[[i]]$Chr[2]) { # Count it regarding its genotype
                    N_Female_Aa[t] = N_Female_Aa[t]+1  # adult Aa female
                  } else if (Juvenile[[i]]$Chr[1]=="A" & Juvenile[[i]]$Chr[2]=="A") { # adult AA female
                    N_Female_AA[t] = N_Female_AA[t]+1
                  } else {  # adult aa female
                    N_Female_aa[t] = N_Female_aa[t]+1
                  }
          
                  each_pref_female_long[(2*N_Female[t])-1] = Female[[N_Female[t]]]$pref[1]  # put pref allele values of this individual in the pref allele vector of adult females
                  each_pref_female_long[2*N_Female[t]] = Female[[N_Female[t]]]$pref[2]    
                  each_thr_female_long[(2*N_Female[t])-1] = Female[[N_Female[t]]]$thr[1]  # put thr allele values of this individual in the thr allele vector of adult females
                  each_thr_female_long[2*N_Female[t]] = Female[[N_Female[t]]]$thr[2]    
          
              } # else ends: maturing females are done
              
            } # if ends: juvenile females from the previous year are done 
      
          }  # if ends: juveniles who survived the winter are done    
        } # for ends: Juveniles from previous years matured, survived as juveniles or died
      } # if ends: survival and maturation of juveniles from previous year is done
          
      #########
      ### Survival and aging of OFFSPRING from PREVIOUS YEAR
        # Selection based on the genotypes!
            
      if(N>0) {  # if reproduction occurred last year
        for(i in 1:N) {  # N: the number of offspring that survived the early density-dependent selection
          rand = runif(1)

          if (Offspring[[i]]$Chr[1]!= Offspring[[i]]$Chr[2] & rand<phi_Aa) { # Aa offpsring from last year and survived the winter

            Offspring[[i]]$Age = 1  # becomes one year old

            if (Offspring[[i]]$Sex=="f" & Offspring[[i]]$Age>=mat_f) {  # if it becomes a mature female
      
              N_Female[t] = N_Female[t]+1 # count it as an adult female
              N_Female_Aa[t] = N_Female_Aa[t]+1 # count it as an adult Aa female
              Female[[N_Female[t]]]=Offspring[[i]]  # put it at the end of the current list of adult females
              each_pref_female_long[(2*N_Female[t])-1] = Female[[N_Female[t]]]$pref[1]  # put pref allele values of this individual in the pref allele vector of adult females
              each_pref_female_long[2*N_Female[t]] = Female[[N_Female[t]]]$pref[2]  
              each_thr_female_long[(2*N_Female[t])-1] = Female[[N_Female[t]]]$thr[1]  # put thr allele values of this individual in the thr allele vector of adult females
              each_thr_female_long[2*N_Female[t]] = Female[[N_Female[t]]]$thr[2]
        
            } else if (Offspring[[i]]$Sex=="m" & Offspring[[i]]$Age>=mat_m) {  # if it becomes a mature male
      
              N_Male_Aa[t]=N_Male_Aa[t]+1    # count it as an adult Aa male
              Male_Aa[[N_Male_Aa[t]]] = Offspring[[i]]    # put it at the end of the current list of adult Aa Males
              each_pref_male_Aa_long[(2*N_Male_Aa[t])-1] = Male_Aa[[N_Male_Aa[t]]]$pref[1]  # put pref allele values of this individual in the pref allele vector of adult Aa males
              each_pref_male_Aa_long[2*N_Male_Aa[t]] = Male_Aa[[N_Male_Aa[t]]]$pref[2]                  
              each_thr_male_Aa_long[(2*N_Male_Aa[t])-1] = Male_Aa[[N_Male_Aa[t]]]$thr[1]  # put thr allele values of this individual in the thr allele vector of adult Aa males
              each_thr_male_Aa_long[2*N_Male_Aa[t]] = Male_Aa[[N_Male_Aa[t]]]$thr[2]
        
            }  else { # it is not mature yet
      
              N_Juven[t] = N_Juven[t]+1  # count it as a Juvenile
              N_Juv_Aa[t] = N_Juv_Aa[t]+1 # count it as a Aa juvenile
              Juvenile[[N_Juven[t]]] = Offspring[[i]]  # put it at the end of the current list of Juveniles
              each_thr_juvenile_long[(N_Juven[t]*2)-1] = Juvenile[[N_Juven[t]]]$thr[1]  # put thr allele values of this individual in the thr allele vector of juveniles
              each_thr_juvenile_long[N_Juven[t]*2] = Juvenile[[N_Juven[t]]]$thr[2]
        
              if(Juvenile[[N_Juven[t]]]$Sex=="m") {  # male
        N_Juv_Male[t] = N_Juv_Male[t]+1  # count it as a juvenile male
        } else {  # female
        N_Juv_Female[t] = N_Juv_Female[t]+1  # count it as a juvenile female
        }
        
            }  # aging is done
          }  # if ends: we dealt with the Aa offspring
          
          if (Offspring[[i]]$Chr[1]=="A" & Offspring[[i]]$Chr[2]=="A" & rand<phi_AA) { # AA offspring from last year and survived the winter

            Offspring[[i]]$Age = 1  # becomes one year old
          
            if (Offspring[[i]]$Sex=="m" & Offspring[[i]]$Age>=mat_m) {  # if it becomes a mature AA male
      
              N_Male_AA[t] = N_Male_AA[t]+1    # count it as an adult AA male
              Male_AA[[N_Male_AA[t]]] = Offspring[[i]]  # put it at the end of the current list of adult AA Males 
              each_pref_male_AA_long[(2*N_Male_AA[t])-1] = Male_AA[[N_Male_AA[t]]]$pref[1]  # put pref allele values of this individual in the pref allele vector of adult AA males
              each_pref_male_AA_long[2*N_Male_AA[t]] = Male_AA[[N_Male_AA[t]]]$pref[2]  
              each_thr_male_AA_long[(2*N_Male_AA[t])-1] = Male_AA[[N_Male_AA[t]]]$thr[1]  # put thr allele values of this individual in the thr allele vector of adult AA males
              each_thr_male_AA_long[2*N_Male_AA[t]] = Male_AA[[N_Male_AA[t]]]$thr[2]
        
            } else if (Offspring[[i]]$Sex=="f" & Offspring[[i]]$Age>=mat_f) {  # if it becomes a mature AA female
      
              N_Female[t] = N_Female[t]+1 # count it as an adult female
              N_Female_AA[t] = N_Female_AA[t]+1 # count it as an adult Aa female
              Female[[N_Female[t]]] = Offspring[[i]]  # put it at the end of the current list of adult females
              each_pref_female_long[(2*N_Female[t])-1] = Female[[N_Female[t]]]$pref[1]  # put pref allele values of this individual in the pref allele vector of adult females
              each_pref_female_long[2*N_Female[t]] = Female[[N_Female[t]]]$pref[2]                                  
              each_thr_female_long[(2*N_Female[t])-1] = Female[[N_Female[t]]]$thr[1]  # put thr allele values of this individual in the thr allele vector of adult females
              each_thr_female_long[2*N_Female[t]] = Female[[N_Female[t]]]$thr[2]
        
            } else { # it is not mature yet
      
              N_Juven[t] = N_Juven[t]+1  # count it as a Juvenile
              N_Juv_AA[t] = N_Juv_AA[t]+1  # count it as AA juvenile
              Juvenile[[N_Juven[t]]] = Offspring[[i]]  # put it at the end of the current list of Juveniles
              each_thr_juvenile_long[(N_Juven[t]*2)-1] = Juvenile[[N_Juven[t]]]$thr[1]  # put thr allele values of this individual in the thr allele vector of juveniles
              each_thr_juvenile_long[N_Juven[t]*2] = Juvenile[[N_Juven[t]]]$thr[2]
              
              if (Offspring[[i]]$Sex=="m") {  # male
          N_Juv_Male[t] = N_Juv_Male[t]+1 # count it as a juvenile male
        } else {  # female
          N_Juv_Female[t] = N_Juv_Female[t]+1  # count it as a juvenile female
        }
        
            }  # aging is done
          }  # if ends: we dealt with the AA Offspring
          
          if (Offspring[[i]]$Chr[1]=="a" & Offspring[[i]]$Chr[2]=="a" & rand<phi_aa) {  # aa offpsring from last year and survived the winter
            
            Offspring[[i]]$Age = 1  # becomes one year old
            
            if (Offspring[[i]]$Sex=="f" & Offspring[[i]]$Age>=mat_f) {  # if it becomes a mature female
      
              N_Female[t] = N_Female[t]+1 # count it as an adult female
              N_Female_aa[t] = N_Female_aa[t]+1 # count it as an adult aa female
              Female[[N_Female[t]]] = Offspring[[i]]  # put it at the end of the current list of adult females
              each_pref_female_long[(2*N_Female[t])-1] = Female[[N_Female[t]]]$pref[1]  # put pref allele values of this individual in the pref allele vector of adult females
              each_pref_female_long[2*N_Female[t]] = Female[[N_Female[t]]]$pref[2]          
              each_thr_female_long[(2*N_Female[t])-1] = Female[[N_Female[t]]]$thr[1]  # put thr allele values of this individual in the thr allele vector of adult females
              each_thr_female_long[2*N_Female[t]] = Female[[N_Female[t]]]$thr[2]
        
            } else if (Offspring[[i]]$Sex=="m" & Offspring[[i]]$Age>=mat_m) {  # becomes mature male
              N_Male_aa[t] = N_Male_aa[t]+1    # count it as an adult aa male
              Male_aa[[N_Male_aa[t]]] = Offspring[[i]]  # put it at the end of th current list of adult aa Males
              each_pref_male_aa_long[(2*N_Male_aa[t])-1] = Male_aa[[N_Male_aa[t]]]$pref[1]  # put pref allele values of this individual in the pref allele vector of adult aa males
              each_pref_male_aa_long[2*N_Male_aa[t]] = Male_aa[[N_Male_aa[t]]]$pref[2]  
              each_thr_male_aa_long[(2*N_Male_aa[t])-1] = Male_aa[[N_Male_aa[t]]]$thr[1]  # put thr allele values of this individual in the thr allele vector of adult aa males
              each_thr_male_aa_long[2*N_Male_aa[t]] = Male_aa[[N_Male_aa[t]]]$thr[2]  

            }  else {  # it is not mature yet
              N_Juven[t] = N_Juven[t]+1  # count it as a Juvenile
              N_Juv_aa[t] = N_Juv_aa[t]+1 # count it as a aa juvenile
              Juvenile[[N_Juven[t]]] = Offspring[[i]]  # put it at the end of the current list of Juveniles
              each_thr_juvenile_long[(N_Juven[t]*2)-1] = Juvenile[[N_Juven[t]]]$thr[1]  # put thr allele values of this individual in the thr allele vector of juveniles
              each_thr_juvenile_long[N_Juven[t]*2] = Juvenile[[N_Juven[t]]]$thr[2]              
              
              if (Offspring[[i]]$Sex=="m") {  # male
          N_Juv_Male[t] = N_Juv_Male[t]+1  # count it as a juvenile male
        } else {  # female
          N_Juv_Female[t] = N_Juv_Female[t]+1  # count it as a juvenile female
        }
        
            }  # aging is done
          }  # if ends: we dealt with the aa Offspring
      
        }   # for ends: a proportion of previous year offspring survived the winter and now they are either juveniles or adults
      }  # if ends: we dealt with Offspring from previous year
      
      # Counting all existing males and females in the population before reproduction (no Offspring exists at the moment)  
      N_Male[t] = N_Male_aa[t] + N_Male_Aa[t] + N_Male_AA[t]  # Total number of adult males (survived the winter as adults or became mature this year).
      N_Male_JuvAd[t] = N_Juv_Male[t] + N_Male[t]
      N_Female_JuvAd[t] = N_Juv_Female[t] + N_Female[t]

      ############
      ### Reproduction
        # Production of new Offspring and assignment of their phenotypic sex according to their individual male signal level (influenced by genotype and incubation temperature).
        # Counting Offspring genotypes and phenotypes (sex).
        # Density-dependent selection may occur, if the current population is abundant.
    
      if(calc_success==TRUE) {  # save vectors for counting offpring of mothers with alleles preference for sex-reversed (CR) and normal (CN) males
	  
	    breeding_success_CR = numeric(length=2)  # number of male and female offspring of females with relative preference for sex-reversed males each year
        breeding_success_CN = numeric(length=2)   # number of male and female offspring of females with relative preference for normal males each year
        is_mot_CR = FALSE # boolean for checking if the mother carries CR
      }	  
	  
      if(calc_linkage==TRUE) {  # save vectors for counting gametes that meet certain requirements related to linkage disequilibrium
        n_CR_lowthr_together = 0  # number of co-occurrences of CR allele and the lower thr allele among gametes of successfully breeding adults
        n_CR_A_together = 0  # number of co-occurrences of allele CR and chromosome A among gametes of successfully breeding adults
        n_lowthr = 0 # total number of copies of the thr allele of lower value among gametes of successfully breeding adults
        n_father_CR = 0  # total number of copies of allele CR inherited from fathers among offspring
        n_mother_CR = 0  # total number of copies of allele CR inherited from mothers among offspring
      }
        
      N_parents=0  # the number of existing mother-father dyads 
                
      if (N_Male_JuvAd[t]>0 & N_Female_JuvAd[t]>0) { # if both phenotypic sexes occur in the population

        if (N_Female[t]>0 & N_Male[t]>0) { # if there are mature males and females currently in the population
    
          mother_father_list = vector("list", N_Female[t])  # This list will contain the individual IDs of members of each mother-father dyad mating this year.
                                                            # Each female will be represented maximum once and each male maxmum 'libido' times.
  
          libido_AA_male = rep(libido,N_Male_AA[t])  # a vector containing the possible number of copulations left for each AA male
          libido_Aa_male = rep(libido,N_Male_Aa[t])
          libido_aa_male = rep(libido,N_Male_aa[t])
          
          if(length(libido_AA_male)==0) libido_AA_male = 0  # we avoid empty vectors
          if(length(libido_Aa_male)==0) libido_Aa_male = 0
          if(length(libido_aa_male)==0) libido_aa_male = 0
                                      
          permuted_female_IDs = sample(N_Female[t], N_Female[t])  # this vector will contain all IDs from the Female list permuted (the existing adult females)
          father_fert_vector = numeric(N_Female[t])  #  this vector will contain the fertility of the father for each mother-father dyad (in the same order as they occur in the mother_father_list)

          for (i in 1:N_Female[t]) { # choose maximum N_female[t] mother-father dyads  (maximum one mate for each female)
          
            total_male_cop = sum(libido_aa_male) + sum(libido_Aa_male) + sum(libido_AA_male)  # total number of copulations available for this female (i.e. sum libido of males)
          
            male_prob_aa = ifelse(total_male_cop>0, sum(libido_aa_male) / total_male_cop, 0)  # probability that an aa male could become her mating partner (hereafter 'father') based on the proportion and total libido of aa males
            male_prob_Aa = ifelse(total_male_cop>0, sum(libido_Aa_male) / total_male_cop, 0)
            male_prob_AA = ifelse(total_male_cop>0, sum(libido_AA_male) / total_male_cop, 0)  
          
            if (dom_pref==TRUE & dominant%in%(Female[[permuted_female_IDs[i]]]$pref)==TRUE) {  # if the female carries a dominant preference allele, choose according to its value
                      
              if(thr0*2 >= sig_a+sig_A) {  # if we started with ZW/ZZ system: 0 means that the female would exclusively choose an aa or Aa male, 1 means that it would exclusively choose an AA male.
                father_prob_vector = c((1-dominant)*male_prob_aa, (1-dominant)*male_prob_Aa, dominant*male_prob_AA)  # probability of the father being aa, Aa or AA male, respectively            
              } else {  # if we started with XX/XY system: 0 means that the female would exclusively choose an aa male, 1 means that it would exclusively choose an Aa or AA male.
                father_prob_vector = c((1-dominant)*male_prob_aa, dominant*male_prob_Aa, dominant*male_prob_AA)  # probability of the father being aa, Aa or AA male, respectively            
              }
              
            } else {  # there is no dominancy or the female does not carry the dominant allele: she will choose according to the mean value of her preference alleles
              
              if (thr0*2 >= sig_a+sig_A) {  # if we started with ZW/ZZ system: 0 means that the female would exclusively choose an aa or Aa male, 1 means that it would exclusively choose an AA male.
                father_prob_vector = c((1-mean(Female[[permuted_female_IDs[i]]]$pref))*male_prob_aa, (1-mean(Female[[permuted_female_IDs[i]]]$pref))*male_prob_Aa, mean(Female[[permuted_female_IDs[i]]]$pref)*male_prob_AA)  # probability of the father being aa, Aa or AA male, respectively            
              } else {  # if we started with XX/XY system: 0 means that the female would exclusively choose an aa male, 1 means that it would exclusively choose an Aa or AA male.
                father_prob_vector = c((1-mean(Female[[permuted_female_IDs[i]]]$pref))*male_prob_aa, mean(Female[[permuted_female_IDs[i]]]$pref)*male_prob_Aa, mean(Female[[permuted_female_IDs[i]]]$pref)*male_prob_AA)  # probability of the father being aa, Aa or AA male, respectively
              }
              
            } # mate genotype probability vector for female choice is done
            
            
            if (sum(father_prob_vector)>0) {  # if the chosen mother can find a suitable male
            
              N_parents = N_parents+1  # we have a new mother-father dyad

              mother_father_list[[N_parents]] = list(mot_id=0, fat_gen="NA", fat_id=0)  # create a new mother-father dyad           
              
              mother_father_list[[N_parents]]$mot_id = permuted_female_IDs[i]  # First column: the ID of the mother from the Female list                                    
              mother_father_list[[N_parents]]$fat_gen = sample(x=c("aa","Aa","AA"), size=1, prob=father_prob_vector)  # Second column: pick the father's genotype based on the probabilities present in father_prob_vector  
                          
              # The father could be an adult aa, Aa or AA male, with a probability defined by the m_ferts (relative fertility), abundance, and female preference:
              if(mother_father_list[[N_parents]]$fat_gen=="aa") {  # aa father
        
                  father_fert_vector[N_parents] = m_fert_aa  # fertility of the father relative to AA
                  mother_father_list[[N_parents]]$fat_id = sample.ok(t=c(1:N_Male_aa[t]),s=1,p=libido_aa_male) # Third column: choose one father, based on its number in the current list of adult aa males 
                  libido_aa_male[mother_father_list[[N_parents]]$fat_id] = libido_aa_male[mother_father_list[[N_parents]]$fat_id]-1  # decrease the number of possible copulations left for this male

              } else if(mother_father_list[[N_parents]]$fat_gen=="Aa") {  # Aa father
        
                  father_fert_vector[N_parents] = m_fert_Aa  # fertility of the father relative to AA
                  mother_father_list[[N_parents]]$fat_id = sample.ok(t=c(1:N_Male_Aa[t]),s=1,p=libido_Aa_male) # Third column: choose one father, based on its number in the current list of adult Aa males 
                  libido_Aa_male[mother_father_list[[N_parents]]$fat_id] = libido_Aa_male[mother_father_list[[N_parents]]$fat_id]-1  # decrease the number of possible copulations left for this male
          
              } else if(mother_father_list[[N_parents]]$fat_gen=="AA") { # AA father
        
                  father_fert_vector[N_parents] = 1  # fertility of AA males is always 1
                  mother_father_list[[N_parents]]$fat_id = sample.ok(t=c(1:N_Male_AA[t]),s=1,p=libido_AA_male) # Third column: choose one father, based on its number in the current list of adult AA males
                  libido_AA_male[mother_father_list[[N_parents]]$fat_id] = libido_AA_male[mother_father_list[[N_parents]]$fat_id]-1  # decrease the number of possible copulations left for this male
          
              }  # mate choice is done
              
              
            }  # if ends: a mother-father dyad has been created with the female if possible
        
          }  # for ends: N_parents mother-father dyads have been created
          
          if (N_parents>0) {  # if at least one adult female found at least one suitable mate among the adult males
            
            N = min(N_max,N_parents*f_fert*mean(father_fert_vector[1:N_parents]))  # the number of offspring to be produced this year
            
            for (i in 1:N) { # produce N offspring
                                      
              parents_ID = sample.ok(t=c(1:N_parents),s=1, p=father_fert_vector[1:N_parents])  # choose a parent pair for the offspring, with relative probabilities being based on the fathers' fertility

              sig_env = rnorm(1, mean = m_sig_env, sd = sd_indivsig)  # individual level of environmental male signal
                  
              # Offspring inherits sex chromosome, threshold and preference genotypes
              if(mother_father_list[[parents_ID]]$fat_gen=="aa") {  # the father is aa male
        
                N_Father_aa[t] = N_Father_aa[t]+1
                Offspring[[i]]$Chr[1] = Male_aa[[mother_father_list[[parents_ID]]$fat_id]]$Chr[sample(2,1)]
                Offspring[[i]]$Chr[2] = Female[[mother_father_list[[parents_ID]]$mot_id]]$Chr[sample(2,1)]
                Offspring[[i]]$thr[1] = Male_aa[[mother_father_list[[parents_ID]]$fat_id]]$thr[sample(2,1)]
                Offspring[[i]]$thr[2] = Female[[mother_father_list[[parents_ID]]$mot_id]]$thr[sample(2,1)]
                Offspring[[i]]$pref[1] = Male_aa[[mother_father_list[[parents_ID]]$fat_id]]$pref[sample(2,1)]
                Offspring[[i]]$pref[2] = Female[[mother_father_list[[parents_ID]]$mot_id]]$pref[sample(2,1)]
        
              } else if(mother_father_list[[parents_ID]]$fat_gen=="Aa") {  # the father is Aa male
        
                N_Father_Aa[t] = N_Father_Aa[t]+1
                Offspring[[i]]$Chr[1] = Male_Aa[[mother_father_list[[parents_ID]]$fat_id]]$Chr[sample(2,1)]
                Offspring[[i]]$Chr[2] = Female[[mother_father_list[[parents_ID]]$mot_id]]$Chr[sample(2,1)]
                Offspring[[i]]$thr[1] = Male_Aa[[mother_father_list[[parents_ID]]$fat_id]]$thr[sample(2,1)]
                Offspring[[i]]$thr[2] = Female[[mother_father_list[[parents_ID]]$mot_id]]$thr[sample(2,1)]
                Offspring[[i]]$pref[1] = Male_Aa[[mother_father_list[[parents_ID]]$fat_id]]$pref[sample(2,1)]
                Offspring[[i]]$pref[2] = Female[[mother_father_list[[parents_ID]]$mot_id]]$pref[sample(2,1)]         
        
              } else if (mother_father_list[[parents_ID]]$fat_gen=="AA") {  # the father is AA male
        
                N_Father_AA[t] = N_Father_AA[t]+1
                Offspring[[i]]$Chr[1] = Male_AA[[mother_father_list[[parents_ID]]$fat_id]]$Chr[sample(2,1)]
                Offspring[[i]]$Chr[2] = Female[[mother_father_list[[parents_ID]]$mot_id]]$Chr[sample(2,1)]
                Offspring[[i]]$thr[1] = Male_AA[[mother_father_list[[parents_ID]]$fat_id]]$thr[sample(2,1)]
                Offspring[[i]]$thr[2] = Female[[mother_father_list[[parents_ID]]$mot_id]]$thr[sample(2,1)]
                Offspring[[i]]$pref[1] = Male_AA[[mother_father_list[[parents_ID]]$fat_id]]$pref[sample(2,1)]
                Offspring[[i]]$pref[2] = Female[[mother_father_list[[parents_ID]]$mot_id]]$pref[sample(2,1)]
        
              }  # Offspring genotype has been set
             
       # Determine the phenotypic sex and count geno- and phenotypes:
              if (Offspring[[i]]$Chr[1]=="A" & Offspring[[i]]$Chr[2]=="A") {  # AA offspring
                N_Offspring_AA[t] = N_Offspring_AA[t]+1
        
                if (sig_A*2 + sig_env > Offspring[[i]]$thr[1] + Offspring[[i]]$thr[2]) {    
                  Offspring[[i]]$Sex = "m"      
                } else {
                  Offspring[[i]]$Sex = "f"
                }
        
              } else if (Offspring[[i]]$Chr[1]!= Offspring[[i]]$Chr[2]) {  # Aa offspring
                N_Offspring_Aa[t]=N_Offspring_Aa[t]+1
        
                if (sig_A + sig_a + sig_env > Offspring[[i]]$thr[1] + Offspring[[i]]$thr[2]) {    
                  Offspring[[i]]$Sex = "m"      
                } else {
                  Offspring[[i]]$Sex = "f"
                }
        
              } else if (Offspring[[i]]$Chr[1]=="a" & Offspring[[i]]$Chr[2]=="a") {  # aa offspring
                N_Offspring_aa[t] = N_Offspring_aa[t]+1
                
                if(aa_no_masc==TRUE) {  # if aa cannot masculinize
        
                  Offspring[[i]]$Sex = "f"
          
                } else if(sig_a*2 + sig_env > Offspring[[i]]$thr[1] + Offspring[[i]]$thr[2]) {  # aa can masculinize and the individual becomes a male
                  Offspring[[i]]$Sex = "m"      
                } else {  # aa can masculinize but this individual is a female
                  Offspring[[i]]$Sex = "f"
                }
        
              } # Sex of the offspring has been determined

              each_thr_offspring_long[(i*2)-1] = Offspring[[i]]$thr[1]  # put thr allele values of this individual in the thr allele vector of offspring
              each_thr_offspring_long[i*2] = Offspring[[i]]$thr[2]                    

              if (Offspring[[i]]$Sex=="m") {  # male
                N_Off_Male[t] = N_Off_Male[t]+1  #count it as a male offspring
              } else {  # female
                N_Off_Female[t] = N_Off_Female[t]+1  # count it as a female offspring
              }
			  
              if (calc_success==TRUE) {

  	            # The mother is shows relative preference for sex-reversed males if it carries the preference allele of lower value, and that allele is either dominant or the alleles show intermediate inheritance. 
  	            if (dom_pref==TRUE & dominant==min(def_pref0) & dominant%in%Female[[mother_father_list[[parents_ID]]$mot_id]]$pref==TRUE) {	# CR is dominant:CR females are CR CN or CR CR
                  is_mot_CR = TRUE
  	            } else if (dom_pref==TRUE & dominant==max(def_pref0) & dominant%in%Female[[mother_father_list[[parents_ID]]$mot_id]]$pref==FALSE) {  # CN is dominant: CR females are CR CR
                  is_mot_CR = TRUE				
  	            } else if (dom_pref==FALSE & min(def_pref0)%in%Female[[mother_father_list[[parents_ID]]$mot_id]]$pref==TRUE) {  # intermediate inheritance: CR females are CR CN or CR CR
                  is_mot_CR = TRUE
  	            } else {
                  is_mot_CR = FALSE
  	            }

                if (is_mot_CR==TRUE) {  # the mother shows a relative preference for sex-reversed males
                  if (Offspring[[i]]$Sex=="m") {
                    breeding_success_CR[1] = breeding_success_CR[1] + 1  # increase the number of male offspring by one
                  } else {
                    breeding_success_CR[2] = breeding_success_CR[2] + 1  # increase the number of female offspring by one                  
                  }
  	            } else {  # the mother shows a relative preference for normal males
                  if (Offspring[[i]]$Sex=="m") {
                    breeding_success_CN[1] = breeding_success_CN[1] + 1  # increase the number of male offspring by one
                  } else {
                    breeding_success_CN[2] = breeding_success_CN[2] + 1  # increase the number of female offspring by one                  
                  }
                }

              }  # if ends: male and female offspring has been counted regarding the mother's mating preference, if required
			  
              if(calc_linkage==TRUE) {
			  
                if(Offspring[[i]]$pref[1]==min(def_pref0) & Offspring[[i]]$thr[1]==min(thr0_vector)) { # inherited both CR and the lower thr allele from the father
                  n_CR_lowthr_together = n_CR_lowthr_together + 1  # count gametes that carried both allele CR and the lower thr allele
                }
				
                if(Offspring[[i]]$pref[2]==min(def_pref0) & Offspring[[i]]$thr[2]==min(thr0_vector)) { # inherited both CR and the lower thr allele from the mother
                  n_CR_lowthr_together = n_CR_lowthr_together + 1  # count gametes that carried both allele CR and the lower thr allele
                }

                if(Offspring[[i]]$pref[1]==min(def_pref0) & Offspring[[i]]$Chr[1]=="A") {  # inherited both CR and A from the father
                  n_CR_A_together = n_CR_A_together + 1  # count gametes that carried both allele CR and chromosome A
                }
				
                if(Offspring[[i]]$pref[2]==min(def_pref0) & Offspring[[i]]$Chr[2]=="A") {  # inherited both CR and A from the mother
                  n_CR_A_together = n_CR_A_together + 1  # count gametes that carried both allele CR and chromosome A
                }
				
                n_lowthr = n_lowthr + sum(c(Offspring[[i]]$thr)==min(thr0_vector)) # total number of copies of the thr allele of lower value
                n_father_CR = n_father_CR + sum(c(Offspring[[i]]$pref[1])==min(def_pref0))  # total number of copies of allele CR inherited from fathers
                n_mother_CR = n_mother_CR + sum(c(Offspring[[i]]$pref[2])==min(def_pref0))  # total number of copies of allele CR inherited from mothers
				
              }  # if ends: co-occurrences of certain alleles have been counted
        
            } # for ends: offspring production is done
              

              N_Off[t] = N  # Number of Offspring in year t. N has been recalculated at the beginning of the Reproduction!
      
              # Proportion of each geno- and phenotype among the Offspring
              AA_Offspring[t] = N_Offspring_AA[t] / N
              Aa_Offspring[t] = N_Offspring_Aa[t] / N
              aa_Offspring[t] = N_Offspring_aa[t] / N
              Male_Offspring[t] = N_Off_Male[t] / N
              Female_Offspring[t] = N_Off_Female[t] / N
			  
          } else {  # there were no matings, although both adult males and females occurred
            N=0    # if no reproduction occurred, number of Offspring=0 in the year
            AA_Offspring[t] = 0
            Aa_Offspring[t] = 0
            aa_Offspring[t] = 0
            Male_Offspring[t] = 0
            Female_Offspring[t] = 0    
            N_Off[t] = 0
          } # A total of N_parents created N offspring that survived density-dependent selection (N may be 0).

        } else {  # there were no matings, because only one sex was represented among adults (although both sexes occur on population level)
    
          N=0    # if no reproduction occurred, number of Offspring=0 in the year
          AA_Offspring[t] = 0
          Aa_Offspring[t] = 0
          aa_Offspring[t] = 0
          Male_Offspring[t] = 0
          Female_Offspring[t] = 0    
          N_Off[t] = 0
      
        } # Mating (if occurred) is done.
            
      } else { # if either one of the phenotypic sexes disappeared from the population, it goes extinct
    
        extinct=T
    
      } # Reproduction is done or if there are no individuals left, the populations is asssumed to be extinct!
    

      if(t/50==round(t/50)) {
	    print(paste("t=", t))  # print t in every 50 year, to show where we are in the simulation process
	  }

      ########
      # Count the overall relative frequency of Aa and AA genotypes and chromosome A in the population after reproduction
    
      N_Aa_all = N_Offspring_Aa[t] + N_Juv_Aa[t] + N_Female_Aa[t] + N_Male_Aa[t]  # temporary calculator: number of Aa individuals in the population after reproduction
      N_AA_all = N_Offspring_AA[t] + N_Juv_AA[t] + N_Male_AA[t] + N_Female_AA[t]  # temporary calculator: number of AA individuals in the population after reproduction

    
      N_Indiv_All[t] = N_Off[t] + N_Juven[t] + N_Male[t] + N_Female[t] # full population size after reproduction
      Aa_All[t] = N_Aa_all / N_Indiv_All[t]  # relative frequency of genotype Aa across all age groups
      AA_All[t] = N_AA_all / N_Indiv_All[t]
      A_All[t] = ((2*N_AA_all) + N_Aa_all) / (2*N_Indiv_All[t])  # relative frequency of chromosome A across all age groups

      N_Male_All[t] = N_Male_JuvAd[t] + N_Off_Male[t]  # number of males across all age groups after reproduction
      N_Female_All[t] = N_Female_JuvAd[t] + N_Off_Female[t]  # number of females across age groups after reproduction
      
      ########
      # By now, individuals in each age group have been counted. We check here if genotypes AA and Aa disappeared from the population:
    
      if (!extinct_AA & AA_All[t]==0) {  # if AA individual(s) were continuously present in the population until last year, but now there are none
        t_ext_AA_vector[k] = t - t_burn_in  # we record that genotype AA now vanished from the population for the first time
        extinct_AA = T  # extinction time is calculated from the beginning of climate warming period (number of years since the end of the burn-in period)
      }
      
      if (!extinct_Aa & Aa_All[t]==0) {  # if Aa individual(s) were continuously present in the population until last year, but now there are none
        t_ext_Aa_vector[k] = t - t_burn_in
        extinct_Aa = T
      }
      
      if (!extinct_A & A_All[t]==0) {  # if chromosome A just disappeared from the population
        t_ext_A_vector[k] = t - t_burn_in
        extinct_A = T
      }
      
      #######
    # Calculate proportion of sex-reversed individuals among phenotypic males and females, respectively.
    
      if (N_Male[t]>0) {  # there were adult males
        if (thr0*2 >= sig_a+sig_A) {  # if we started with ZW/ZZ system
          SR_in_Male[t] = (N_Male_Aa[t]+N_Male_aa[t])/N_Male[t]  # proportion of masculinized genetic females within all phenotypic males
        } else {  # if we started with XX/XY system
          SR_in_Male[t] = N_Male_aa[t]/N_Male[t] # proportion of masculinized genetic females within all phenotypic males
        }
      } else {
        SR_in_Male[t] = NA
      }
      
      if (N_Female[t]>0) {  # there were adult females
        
        if (thr0*2 >= sig_a+sig_A) {  # if we started with ZW/ZZ system
          SR_in_Female[t] = N_Female_AA[t]/N_Female[t]  # proportion of feminized genetic males within all phenotypic females
        } else {  # if we started with XX/XY system
         SR_in_Female[t] = (N_Female_Aa[t] + N_Female_AA[t])/N_Female[t] # proportion of feminized genetic males within all phenotypic females
        }
      } else {
        SR_in_Female[t]=NA
      }
      
      if (N_Off[t]>0) {  # if offspring was produced
      
        if (thr0*2 >= sig_a+sig_A) {  # if we started with ZW/ZZ system
          SR_in_Father[t] = (N_Father_Aa[t] + N_Father_aa[t])/N_Off[t]  # proportion of masculinized genetic females within all fathering events
        } else {  # if we started with XX/XY system
          SR_in_Father[t] = N_Father_aa[t]/N_Off[t]  # proportion of masculinized genetic females within all fathering events
        }

        Aa_in_Father[t] = N_Father_Aa[t]/N_Off[t]      # proportion of Aa masculinized genetic females within all fathering events
        aa_in_Father[t] = N_Father_aa[t]/N_Off[t]      # proportion of aa masculinized genetic females within all fathering events
		
        if(calc_success==TRUE) {
          prop_daughter_CR[t] = breeding_success_CR[2] / (breeding_success_CR[2]  + breeding_success_CR[1]) # propoprtion of females among offpsring of mothers with relative preference for sex-reversed males
          prop_daughter_CN[t] = breeding_success_CN[2] / (breeding_success_CN[2]  + breeding_success_CN[1]) # propoprtion of females among offpsring of mothers with relative preference for normal males
        }
		
		if(calc_linkage==TRUE) {  # linkage disequilibrium between preference allele CR and the lower thr allele, and CR and chromosome A should be calculated!
			
          n_A = 2*N_Offspring_AA[t] + N_Offspring_Aa[t]  # total number of A copies among gametes that produced offspring
          n_CR = n_father_CR + n_mother_CR  # total number of CR copies among gametes that produced offspring
		  
          ld_CR_lowthr[t] = n_CR_lowthr_together/(2*N_Off[t]) - (n_CR/(2*N_Off[t]))*(n_lowthr/(2*N_Off[t]))  # linkage disequilibrium calculated as: P(CR,lowthr) - P(CR)*P(lowthr)
          ld_CR_A[t] = n_CR_A_together/(2*N_Off[t]) - (n_CR/(2*N_Off[t]))*(n_A/(2*N_Off[t]))
			  
          prop_father_CR[t] = n_father_CR / N_Off[t]  # proportion of allele CR among preference alleles inherited from fathers
          prop_mother_CR[t] = n_mother_CR / N_Off[t]
			  
        }
		
      } else { # to avoid error when N_Off = 0
        SR_in_Father[t] = NA
        Aa_in_Father[t] = NA
        aa_in_Father[t] = NA
        prop_daughter_CR[t] = NA
        prop_daughter_CN[t] = NA
        ld_CR_lowthr[t] = NA
        ld_CR_A[t] = NA
        prop_father_CR[t] = NA
        prop_mother_CR[t] = NA
      }

    #######
      # List pref and thr alleles present present in certain lists of individuals
      each_pref_female = numeric(2*N_Female[t]) # save memory for temporary vectors with length differing between years each year: these vectors are re-written every year!  
      each_pref_male_aa = numeric(2*N_Male_aa[t])
      each_pref_male_Aa = numeric(2*N_Male_Aa[t])
      each_pref_male_AA = numeric(2*N_Male_AA[t])      
      each_thr_female = numeric(2*N_Female[t])
      each_thr_male_AA = numeric(2*N_Male_AA[t])
      each_thr_male_Aa = numeric(2*N_Male_Aa[t])
      each_thr_male_aa = numeric(2*N_Male_aa[t])
      each_thr_juvenile = numeric(2*N_Juven[t])
      each_thr_offspring = numeric(2*N_Off[t])
      

      
      if(N_Female[t]>0) {  # if there are adult females in the population, save statistics for thr alleles in females

        each_pref_female = head(each_pref_female_long,n=2*N_Female[t])  #  all copies of preference alleles actually occurring among the adult females in this year
        pref_female = unique(each_pref_female) # list each allele present in adult females, with duplicate elements removed
      
        each_thr_female = head(each_thr_female_long,n=2*N_Female[t])  #  all copies of threshold alleles actually occurring among the adult females in this year
        thr_female = unique(each_thr_female) # list each allele present in adult females, with duplicate elements removed

        N_thr_alleles_Female[t] = length(thr_female)  # number of different thr alleles among adult females
        Min_thr_alleles_Female[t] = min(thr_female)  # minimum value of thr alleles among adult females
        Max_thr_alleles_Female[t] = max(thr_female)  # maximum value of thr alleles among adult females

      } else {
        each_pref_female = NULL
        each_thr_female = NULL
        thr_female = NULL
        N_thr_alleles_Female[t] = 0
        Min_thr_alleles_Female[t] = NA
        Max_thr_alleles_Female[t] = NA
      }
      
      if (N_Male_AA[t]>0) {
        each_thr_male_AA = head(each_thr_male_AA_long, n=2*N_Male_AA[t])  # all copies of threshold alleles actually occurring among the adult AA males in this year
        each_pref_male_AA = head(each_pref_male_AA_long,n=2*N_Male_AA[t])  # all copies of preference alleles actually occurring among the adult AA males in this year
      }
      
      if (N_Male_Aa[t]>0) {
        each_thr_male_Aa = head(each_thr_male_Aa_long, n=2*N_Male_Aa[t])  # all copies of threshold alleles actually occurring among the adult Aa males in this year
        each_pref_male_Aa = head(each_pref_male_Aa_long,n=2*N_Male_Aa[t])  # all copies of preference alleles actually occurring among the adult Aa males in this year  
      }
      
      if (N_Male_aa[t]>0) {
        each_thr_male_aa = head(each_thr_male_aa_long, n=2*N_Male_aa[t])  # all copies of threshold alleles actually occurring among the adult aa males in this year
        each_pref_male_aa = head(each_pref_male_aa_long,n=2*N_Male_aa[t])  # all copies of preference alleles actually occurring among the adult aa males in this year  
      }
      
      if (N_Male_AA[t]+N_Male_Aa[t]+N_Male_aa[t]>0) {  # if there are adult males in the population, save statistics for thr alleles in males
      
        thr_male = unique(c(each_thr_male_AA, each_thr_male_Aa, each_thr_male_aa))  # list each allele present in adult males, with duplicate elements removed

        N_thr_alleles_Male[t] = length(thr_male)  # number of different thr alleles among adult males
        Min_thr_alleles_Male[t] = min(thr_male)  # minimum value of thr alleles among adult males
        Max_thr_alleles_Male[t] = max(thr_male)  # maximum value of thr alleles among adult males

      } else {
        thr_male = NULL
        N_thr_alleles_Male[t] = 0
        Min_thr_alleles_Male[t] = NA
        Max_thr_alleles_Male[t] = NA
      }
      
      if(N_Juven[t]>0) {
        each_thr_juvenile = head(each_thr_juvenile_long, n=2*N_Juven[t])  # all copies of threshold alleles actually occurring among the adult juveniles in this year   
        thr_juvenile = unique(each_thr_juvenile)
      } else {
        thr_juvenile=NULL
      }
      
      if(N_Off[t]>0) {
        each_thr_offspring = head(each_thr_offspring_long, n=2*N_Off[t])  # all copies of threshold alleles actually occurring among the adult offspring in this year   
        thr_offspring = unique(each_thr_offspring)
      } else {
        thr_offspring = NULL
      }
          
      
      # Save the total number of alleles present in the population:
      thr_all = unique(c(thr_female, thr_male, thr_juvenile, thr_offspring))
      thr_adult = unique(c(thr_female, thr_male))
      N_thr_alleles_All[t] = length(thr_all)  # number of thr alleles present in the total population
      N_thr_alleles_Adult[t] = length(thr_adult)  # number of thr alleles present among adults
      
      for (j in 1:length(def_pref0)) {  # save proportion of each preference allele among adult females
          
        if(N_Female[t]>0) {  # if there are adult females in the population
          pref_prop_fem = sum(each_pref_female==def_pref0[j]) / (2*(N_Female_Aa[t] + N_Female_AA[t] + N_Female_aa[t]))
          if(pref_prop_fem>0) {
            pref_prop_fem_matrix[t,j] = pref_prop_fem  # if the allele of concern occurs in adult females, update the corresponding element of the matrix (otherwise its value remains 0)
          }      
        }

        if(N_Male[t]>0) {  # if there are adult males in the population
          pref_prop_male = sum(c(each_pref_male_aa, each_pref_male_Aa, each_pref_male_AA)==def_pref0[j]) / (2*(N_Male_Aa[t] + N_Male_AA[t] + N_Male_aa[t]))
          if(pref_prop_male>0) {
            pref_prop_male_matrix[t,j] = pref_prop_male  # if the allele of concern occurs in adult males, update the corresponding element of the matrix (otherwise its value remains 0)
          }
        }
      }
      
      for (j in 1:n_thr0) {
          
        if(N_Female[t]>0) {  # if there are adult females in the population
          allele_prop_fem = sum(each_thr_female==thr0_vector[j]) / (2*(N_Female_Aa[t] + N_Female_AA[t] + N_Female_aa[t]))
          if(allele_prop_fem>0) {
            thr_prop_fem_matrix[t,j] = allele_prop_fem  # if the allele of concern occurs in adult females, update the corresponding element of the matrix (otherwise its value remains 0)
          }
        }

        if(N_Male[t]>0) {  # if there are adult males in the population          
          allele_prop_male = sum(c(each_thr_male_AA, each_thr_male_Aa, each_thr_male_aa)==thr0_vector[j])/(2*(N_Male_Aa[t]+N_Male_AA[t]+N_Male_aa[t]))      
          if(allele_prop_male>0) {
            thr_prop_male_matrix[t,j] = allele_prop_male  # if the allele of concern occurs in adult males, update the corresponding element of the matrix (otherwise its value remains 0)
          }
        }
      }
      
      
    } # while ends: the simulation ran for t_burn_in + t_Max years, or the population went extinct

    
    #####################################################
    # Create summary vectors and some informative plots #
    #####################################################
    
    t_end=t-1  # plots will be drawn up to the year before the last one (because if the population went extinct, reproduction and SR_in_Father cannot be calculated in the last year)
    print(paste("t_end=", t_end))
      
    # Count relative frequency of each chromosome among ADULT males and females, respectively
    Prop_A_in_males = (N_Male_Aa + 2*N_Male_AA)  / (N_Male_Aa + N_Female_Aa + 2*(N_Male_AA + N_Female_AA))
    Prop_A_in_females = N_Female_Aa+N_Female_AA  / (N_Male_Aa + N_Female_Aa + 2*(N_Male_AA + N_Female_AA))
    Prop_a_in_males = (N_Male_Aa + 2*N_Male_aa)  / (N_Male_Aa + N_Female_Aa + 2*(N_Male_aa + N_Female_aa))
    Prop_a_in_females = (N_Female_Aa + 2*N_Female_aa)  / (N_Male_Aa + N_Female_Aa + 2*(N_Male_aa + N_Female_aa))
    
    # Relative frequency of geno- and phenotypes among juveniles
    AA_Juv = N_Juv_AA / N_Juven
    Aa_Juv = N_Juv_Aa / N_Juven
    aa_Juv = N_Juv_aa / N_Juven
    Male_Juv = N_Juv_Male / N_Juven  # proportion of all phenotypic males among juveniles

    # Relative frequency of geno- and phenotypes among adults    
    aa_Adult = (N_Male_aa+N_Female_aa) / (N_Male+N_Female)
    Aa_Adult = (N_Male_Aa+N_Female_Aa) / (N_Male+N_Female)
    AA_Adult = (N_Male_AA+N_Female_AA) / (N_Male+N_Female)
    Male = N_Male/(N_Male+N_Female)  # proportion of all phenotypic males among adults
    
    # Relative frequency of genotypes in the overall population (after reproduction!)
    Male_All = N_Male_All / (N_Off + N_Juven + N_Male + N_Female)

    if (thr0*2 >= sig_a+sig_A) {  # if we started with ZW/ZZ system
      Masc_rate = (N_Male_Aa + N_Male_aa) / (N_Male_Aa+N_Female_Aa + N_Male_aa+N_Female_aa)  # Masculinized: aa and Aa males
    } else {  # if we started with XX/XY system
      Masc_rate = N_Male_aa / (N_Male_aa + N_Female_aa)  # Masculinized: aa males
    }    
    
    
    if (k==1) {  # plots will be drawn from the first run only!

      color_vector = 1:n_thr0
    
    
      #########
      ### Plots from the Burn-in period
    
      if (show_burn_in==TRUE) {
      
        par(mfrow=c(2,2))
        par(oma=c(0,0,2,0))
        
        B1= matrix(c(aa_Adult[1:t_burn_in], Aa_Adult[1:t_burn_in], AA_Adult[1:t_burn_in],
          Prop_a_in_males[1:t_burn_in], Prop_A_in_males[1:t_burn_in]), ncol=5)
        B2=matrix(c(Male[1:t_burn_in], SR_in_Male[1:t_burn_in], SR_in_Father[1:t_burn_in], SR_in_Female[1:t_burn_in]), ncol=4)

        matplot(0:(t_burn_in-1), B1, xlab="burn-in year", pch=c(16,16,16,17,17), ylab="Proportion in adults", 
          col=c("red", "green", "blue", "pink", "yellow"), ylim=c(0,1))
        legend("topleft", ncol=2, legend=c("aa","Aa","AA","a in males","A in males"), pch=c(16,16,16,17,17), col=c("red", "green", "blue", "pink", "yellow"))
        matplot(0:(t_burn_in-1), B2, xlab="burn-in year", ylab="Proportion in adults", ylim=c(0,1),
          col=c(1, "purple1", "orange","green"), pch=16:19)
        legend("topleft", legend=c("males in adults","sex-reversed in males","sex-reversed in fathers", "sex-reversed in females"), 
      col=c(1, "purple1", "orange", "green"), pch=16:19)    

        pch_vector=  c(numeric(length=n_thr0)+1) # vector of symbols: circles depict thr alleles  
        

        Prop_pref_burnin = pref_prop_fem_matrix[1:t_burn_in, 1:length(def_pref0)]
        Prop_fem_burnin = thr_prop_fem_matrix[1:t_burn_in, 1:n_thr0]
        Prop_male_burnin = thr_prop_male_matrix[1:t_burn_in, 1:n_thr0]
        
        matplot(0:(t_burn_in-1), Prop_pref_burnin, xlab="year", ylab="Prop preference in adult female",
          col=1:(length(def_pref0)), pch=c(numeric(length=length(def_pref0)+15)), ylim=c(0,1))
        legend("topleft", ncol=3, legend=round(def_pref0,digits=4), col=1:(length(def_pref0)), pch=c(numeric(length=length(def_pref0)+15)), cex = 0.75)

          
        matplot(0:(t_burn_in-1), Prop_fem_burnin, xlab="year", ylab="Prop thr in adult female",
          col=color_vector, pch=pch_vector, ylim=c(0,1))
        legend("topleft", ncol=3, legend=round(thr0_vector[1:n_thr0],digits=4), col=color_vector, pch=pch_vector, cex = 0.75)

        matplot(0:(t_burn_in-1), Prop_male_burnin, xlab="year", ylab="Prop thr in adult male",
          col=color_vector, pch=pch_vector, ylim=c(0,1))
        legend("topleft", ncol=3, legend=round(thr0_vector[1:n_thr0], digits=4), col=color_vector, pch=pch_vector, cex = 0.75)

        
      } # if ends: plots for the burn-in period have been created
      
      par(mfrow=c(2,2),mar=c(4,4,1,1))
      par(oma=c(0,0,0,0))
    
      ########
      # Plots after the burn-in period 
      
      ## Number of adults regarding phenotypic sex and number of sex-reversed individuals 
      # in the beginning, in the middle and at the end of the run:
    
      t1 = t_burn_in
      t2 = trunc(t_burn_in+(t_end-t_burn_in)/2)
      t3 = t_end
    
      Num_Sexes_Adult=matrix(c( N_Female_AA[t1], N_Female_Aa[t1], N_Female_aa[t1], N_Male_AA[t1], N_Male_Aa[t1], N_Male_aa[t1],
        N_Female_AA[t2], N_Female_Aa[t2], N_Female_aa[t2], N_Male_AA[t2], N_Male_Aa[t2], N_Male_aa[t2],
        N_Female_AA[t3], N_Female_Aa[t3], N_Female_aa[t3], N_Male_AA[t3], N_Male_Aa[t3], N_Male_aa[t3]), ncol=3)
      Num_Ages=matrix(c(N_Off[t1], N_Juven[t1], (N_Female[t1]+N_Male[t1]),
        N_Off[t2], N_Juven[t2], (N_Female[t1]+N_Male[t2]),
        N_Off[t3], N_Juven[t3], (N_Female[t1]+N_Male[t3])), ncol=3)
    
      barplot (Num_Ages, col=c("yellow", "green", "pink"),names.arg=c(paste("t=",as.character(0)),
          paste("t=",as.character(trunc((t_end-t_burn_in)/2))), paste("t=",as.character(t_end-t_burn_in))), beside=T, ylab="No. individuals")
      legend("topleft", legend=c("Offspring", "Juvenile", "Adult"), fill=c("yellow", "green", "pink"), bty="n")

      barplot (Num_Sexes_Adult, col=c("orchid","coral", "rosybrown1", "royalblue1", "skyblue", "orchid4"),names.arg=c(paste("t=",as.character(0)),
        paste("t=",as.character(trunc((t_end-t_burn_in)/2))), paste("t=",as.character(t_end-t_burn_in))), beside=T, ylab="No. individuals")
      legend("topleft", legend=c("AA Female","Aa Female", "aa Female", "AA Male", "Aa Male", "aa Male"),
      fill=c("orchid","coral", "rosybrown1", "royalblue1", "skyblue", "orchid4"), bty="n")

      ## Continuous changes in the relative frequency of each genotype and chromosome among adults,
        # proportion of males among adults, sex-reversed among males and sex-reversed among fathers:
    
      M1= matrix(c(aa_Adult[t_burn_in:(t_end-1)], Aa_Adult[t_burn_in:(t_end-1)], AA_Adult[t_burn_in:(t_end-1)],
        Prop_a_in_males[t_burn_in:(t_end-1)], Prop_A_in_males[t_burn_in:(t_end-1)]), ncol=5)
      
      M2=matrix(c(Male[t_burn_in:(t_end-1)], SR_in_Male[t_burn_in:(t_end-1)], SR_in_Father[t_burn_in:(t_end-1)], SR_in_Female[t_burn_in:(t_end-1)], 
        Masc_rate[t_burn_in:(t_end-1)]), ncol=5)
        
      matplot(0:(t_end-t_burn_in-1), M1, xlab="Year", pch=c(16,16,16,17,17), ylab="Proportion in adults",
        col=c("red", "green", "blue", "pink", "yellow"), ylim=c(0,1))
      legend("topleft", ncol=2, legend=c("aa","Aa","AA","a_m","A_m"), pch=c(16,16,16,17,17), col=c("red", "green", "blue", "pink", "yellow"))
    
      matplot(0:(t_end-t_burn_in-1), M2, xlab="Year", ylab="Proportion in adults", ylim=c(0,1),
        col=c(1, "purple1", "orange", "green","red"), pch=c(16,17,19,17,1))
      abline(h=0.5)    
      legend("topleft", legend=c("males in adults", "sex-reversed in males","sex-reversed in fathers",
          "sex-reversed in females","males in genetic females"), col=c(1, "purple1", "orange", "green","red"), pch=c(16,17,19,17,1))

      par(mfrow=c(2,1),mar=c(4,4,1,1))
      
      ## Continuous changes in the relative frequencies of preference alleles among adult females
      Prop_pref=pref_prop_fem_matrix[t_burn_in:(t_end-1), 1:length(def_pref0)]

      matplot(0:(t_end-t_burn_in-1), Prop_pref, xlab="Year", ylab="Prop preference in adult female",
        col=1:(length(def_pref0)) , pch=c(numeric(length=length(def_pref0)+15)), ylim=c(0,1))
      legend("topleft", ncol=3, legend=round(def_pref0,digits=4), col=1:(length(def_pref0)), pch=c(numeric(length=length(def_pref0)+15)), cex = 0.75)
      

      ## Continuous changes in proportions of thr alleles among adult females and males, respectively      
      Prop_fem_masc = thr_prop_fem_matrix[t_burn_in:(t_end-1), 1:n_thr0]
      Prop_male_masc = thr_prop_male_matrix[t_burn_in:(t_end-1), 1:n_thr0]

      pch_vector = c(numeric(length=n_thr0)+1) 

      matplot(0:(t_end-t_burn_in-1), Prop_fem_masc, xlab="Year", ylab="Prop thr in adult female",
        col=color_vector, pch=pch_vector, ylim=c(0,1))
      legend("topleft", ncol=3, legend=round(thr0_vector[1:n_thr0], digits=4), col=color_vector, pch=pch_vector, cex = 0.75)
      
      matplot(0:(t_end-t_burn_in-1), Prop_male_masc, xlab="Year", ylab="Prop thr in adult male",
        col=color_vector, pch=pch_vector, ylim=c(0,1))
      legend("topleft", ncol=3, legend=round(thr0_vector[1:n_thr0],digits=4), col=color_vector, pch=pch_vector, cex = 0.75)
      

    } # if ends: plots from the first run are drawn

    
    ###############################################################
    # Create output matrices and vectors from results of each run #
    ###############################################################

    # Within each matrix, a column refers to the run and a row refers to the year!
      # Considered numbers and proportions are counted in the hypothetical autumn (after reproduction and before winter mortality).
    # Within vectors, each element refers to one run.

    ## Output matrices
      # Returned data will contain results from the burn-in period as well!    
    
    Male_Offspring_matrix[1:t_end,k] = Male_Offspring[1:t_end]  # proportion of phenotypic males among offspring (returned as Male_Offspring)
    Male_Juvenile_matrix[1:t_end,k] = Male_Juv[1:t_end]  # proportion of phenotypic males among juveniles (returned as Male_Juvenile)
    Male_matrix[1:t_end,k] = Male[1:t_end]  # proportion of phenotypic males among adults (returned as ASR)
    Male_All_matrix[1:t_end,k] = Male_All[1:t_end]  # overall proportion of phenotypic males across age groups (returned as Male_All)
    
    AA_Juvenile_matrix[1:t_end,k] = AA_Juv[1:t_end]  # proportion of AA individuals among juveniles (returned as AA_Juvenile)
    Aa_Juvenile_matrix[1:t_end,k] = Aa_Juv[1:t_end]  # proportion of Aa individuals among juveniles (returned as Aa_Juvenile)
    AA_Offspring_matrix[1:t_end,k] = AA_Offspring[1:t_end]  # proportion of AA individuals among offspring (returned as AA_Offspring)
    Aa_Offspring_matrix[1:t_end,k] = Aa_Offspring[1:t_end]  # proportion of Aa individuals among offspring (returned as Aa_Offspring)
    Aa_All_matrix[1:t_end,k] = Aa_All[1:t_end]  # overall proportion of Aa individuals across age groups (returned as Aa_All) 
    AA_All_matrix[1:t_end,k] = AA_All[1:t_end]  # overall proportion of AA individuals across age groups (returned as AA_All) 
    
    AA_Adult_Male_matrix[1:t_end,k] = N_Male_AA[1:t_end] / N_Male[1:t_end]  # proportion of of AA individuals among adult males (returned as AA_Adult_Male)
    Aa_Adult_Male_matrix[1:t_end,k] = N_Male_Aa[1:t_end] / N_Male[1:t_end]  # proportion of of Aa individuals among adult males (returned as Aa_Adult_Male)
    aa_Adult_Male_matrix[1:t_end,k] = N_Male_aa[1:t_end] / N_Male[1:t_end]  # proportion of of aa individuals among adult males (returned as aa_Adult_Male)

    AA_Adult_Female_matrix[1:t_end,k] = N_Female_AA[1:t_end] / N_Female[1:t_end]  # proportion of of AA individuals among adult females (returned as AA_Adult_Female)
    Aa_Adult_Female_matrix[1:t_end,k] = N_Female_Aa[1:t_end] / N_Female[1:t_end]  # proportion of of Aa individuals among adult females (returned as Aa_Adult_Female)  
    aa_Adult_Female_matrix[1:t_end,k] = N_Female_aa[1:t_end] / N_Female[1:t_end]  # proportion of of aa individuals among adult females (returned as aa_Adult_Female)
    
    SR_in_Male_matrix[1:t_end,k] = SR_in_Male[1:t_end]  # proportion of sex-reversed individuals among adult phenotypic males (returned as SR_Male)
    SR_in_Female_matrix[1:t_end,k] = SR_in_Female[1:t_end]  # proportion of sex-reversed individuals among adult phenotypic females (returned as SR_Female)
    SR_in_Father_matrix[1:t_end,k] = SR_in_Father[1:t_end]  # proportion of sex-reversed individuals among fathers (returned as SR_Father)
    Masc_rate_matrix[1:t_end,k] = Masc_rate[1:t_end]  # proportion of males among adults with female genotypes. When starting with ZW/ZZ: Aa and aa, 
                                                   # when starting with XX/XY: aa. (returned as Masc_rate)
    
        
    A_All_matrix[1:t_end,k] = A_All[1:t_end]  # relative frequency of chromosome A across age groups (returned as A_All)
    Pop_Size_matrix[1:t_end,k] = N_Indiv_All[1:t_end]  # total population size (returned as Pop_Size)
    N_Adult_matrix[1:t_end,k] = N_Male[1:t_end] + N_Female[1:t_end]  # number of adults (returned as N_Adult)
    N_Juvenile_matrix[1:t_end,k] = N_Juven[1:t_end]  # number of juveniles (returned as N_Juvenile)
    N_Offspring_matrix[1:t_end,k] = N_Off[1:t_end]  # number of offspring (returned as N_Offspring)

    thr0_matrix[,k] = thr0_vector  # list of thr alleles set in the initial population (returned as thr0)

    Prop_thr0_Female_matrix[1:t_end,(k*n_thr0-n_thr0+1):(k*n_thr0)] = thr_prop_fem_matrix[1:t_end,1:n_thr0]  # relative frequency of each thr allele among adult females, 
                                                                                                           # by-column following their by-row order in thr0_matrix (returned as Prop_thr0_Female)
    Prop_thr0_Male_matrix[1:t_end,(k*n_thr0-n_thr0+1):(k*n_thr0)] = thr_prop_male_matrix[1:t_end,1:n_thr0]  # relative frequency of each thr allele among adult males, 
                                                                                                        # by-column following their by-row order in thr0_matrix (returned as Prop_thr0_Male)

    pref0_matrix[,k] = def_pref0  # list of preference alleles set in the initial population (returned as pref0)
  
    Prop_pref_Female_matrix[1:t_end,(k*length(def_pref0)-length(def_pref0)+1):(k*length(def_pref0))] = pref_prop_fem_matrix[1:t_end,1:length(def_pref0)] # proportion of each preference allele
      # among adult females, by-column following their by-row order in pref_prp_fem_matrix (returned as Prop_pref_Fem)
    
    Prop_pref_Male_matrix[1:t_end,(k*length(def_pref0)-length(def_pref0)+1):(k*length(def_pref0))] = pref_prop_male_matrix[1:t_end,1:length(def_pref0)] # proportion of each preference allele
      # among adult males, by-column following their by-row order in pref_prp_male_matrix (returned as Prop_pref_Male)
  
    Prop_daughter_CR_matrix[1:t_end,k] = prop_daughter_CR[1:t_end]  # proportion of phenotypic females among the offspring of females with relative preference for sex-reversed males (returned as Prop_daughter_CR)
  
    Prop_daughter_CN_matrix[1:t_end,k] = prop_daughter_CN[1:t_end]  # proportion of phenotypic females among the offspring of females with relative preference for normal males (returned as Prop_daughter_CN)
	
    LD_CR_lowthr_matrix[1:t_end,k] = ld_CR_lowthr[1:t_end]  # linkage disequilibrium between the allele encoding relative preference for sex-reversed males and the threshold allele of lower value (returned as LD_CR_lowthr)
	
    LD_CR_A_matrix[1:t_end,k] = ld_CR_A[1:t_end]  # linkage disequilibrium between the allele encoding relative preference for sex-reversed males and chromosome A  (returned as LD_CR_A) 
	
    Prop_father_CR_matrix[1:t_end,k] = prop_father_CR[1:t_end]  # proportion of the allele encoding relative preference for sex-reversed males among preference alleles inherited from fathers (returned as Prop_father_CR)
	
    Prop_mother_CR_matrix[1:t_end,k] = prop_mother_CR[1:t_end]  # proportion of the allele encoding relative preference for sex-reversed males among preference alleles inherited from mothers (returned as Prop_mother_CR)


    ## Output vectors
      # Returned data refer to years calculated from the beginning of the "warming period" (that is, the burn-in period is excluded!)
    
    t_ext_pop_vector[k] = ifelse(extinct, t_end - t_burn_in + 1, NA )  # the year when the population became extinct (returned as t_ext_pop)
    # t_ext_AA_vector: saved earlier. Refers to the year when genotype AA first vanished from the population (returned as t_ext_AA)
    # t_ext_Aa_vector: saved earlier. Refers to the year when genotype Aa first vanished from the population (returned as t_ext_Aa)
    # t_ext_A_vector: saved earlier. Refers to the year when chromosome A vanished from the population (returned as t_ext_A)
    
  } # for (k in 1:n_runs) ends: all runs are done

  ##########
  # RETURN #
  ##########
  # Concatenate output matrices and vectors and return them in one list
  
  Output = list(N_Adult = N_Adult_matrix, N_Juvenile = N_Juvenile_matrix, N_Offspring = N_Offspring_matrix, Pop_Size = Pop_Size_matrix,
  ASR = Male_matrix, Male_Offspring = Male_Offspring_matrix, Male_Juvenile = Male_Juvenile_matrix, Male_All = Male_All_matrix,
  AA_Offspring = AA_Offspring_matrix, Aa_Offspring = Aa_Offspring_matrix, AA_Juvenile = AA_Juvenile_matrix, Aa_Juvenile = Aa_Juvenile_matrix,
  AA_Adult_Male = AA_Adult_Male_matrix, Aa_Adult_Male = Aa_Adult_Male_matrix, aa_Adult_Male = aa_Adult_Male_matrix,
  AA_Adult_Female = AA_Adult_Female_matrix, Aa_Adult_Female = Aa_Adult_Female_matrix, aa_Adult_Female = aa_Adult_Female_matrix,
  AA_All = AA_All_matrix, Aa_All = Aa_All_matrix, A_All = A_All_matrix,
  SR_Male = SR_in_Male_matrix,  SR_Female = SR_in_Female_matrix, SR_Father = SR_in_Father_matrix, Masc_rate = Masc_rate_matrix,
  thr0 = thr0_matrix, Prop_thr0_Female = Prop_thr0_Female_matrix, Prop_thr0_Male = Prop_thr0_Male_matrix,
  pref0 = pref0_matrix, Prop_pref_Fem = Prop_pref_Female_matrix, Prop_pref_Male = Prop_pref_Male_matrix, Prop_daughter_CR = Prop_daughter_CR_matrix, 
  Prop_daughter_CN = Prop_daughter_CN_matrix, LD_CR_lowthr = LD_CR_lowthr_matrix, LD_CR_A = LD_CR_A_matrix, Prop_father_CR = Prop_father_CR_matrix, 
  Prop_mother_CR = Prop_mother_CR_matrix, 
  t_ext_pop = t_ext_pop_vector, t_ext_AA = t_ext_AA_vector, t_ext_Aa = t_ext_Aa_vector,
  t_ext_A = t_ext_A_vector)
  
  return(Output)    
  
} # END of function Sexsel_Sexrev_Aa
