## Supplementary material for "Evolutionary and demographic consequences of temperature-induced masculinization under climate warming: the effects of mate choice": SI5

**Table of contents:**

**Figures showing specified results of 100 simulations of each scenario:**

Fig S10. Relative frequency of genotype *aa* among adults.

Fig S11. Relative frequency of genotype *Aa* among adults.

Fig S12. Adult sex ratio.

Fig S13. Masculinization rate.

Fig S14. Relative frequency of the $thr_{low}$ allele.

Fig S15. Relative frequency of preference allele $C_R$.

Fig S16. Effective population size.

**Figure showing median values across 100 simulations:**

Fig S17. Median effective population size during the last decades before extinction in originally XX/XY populations.

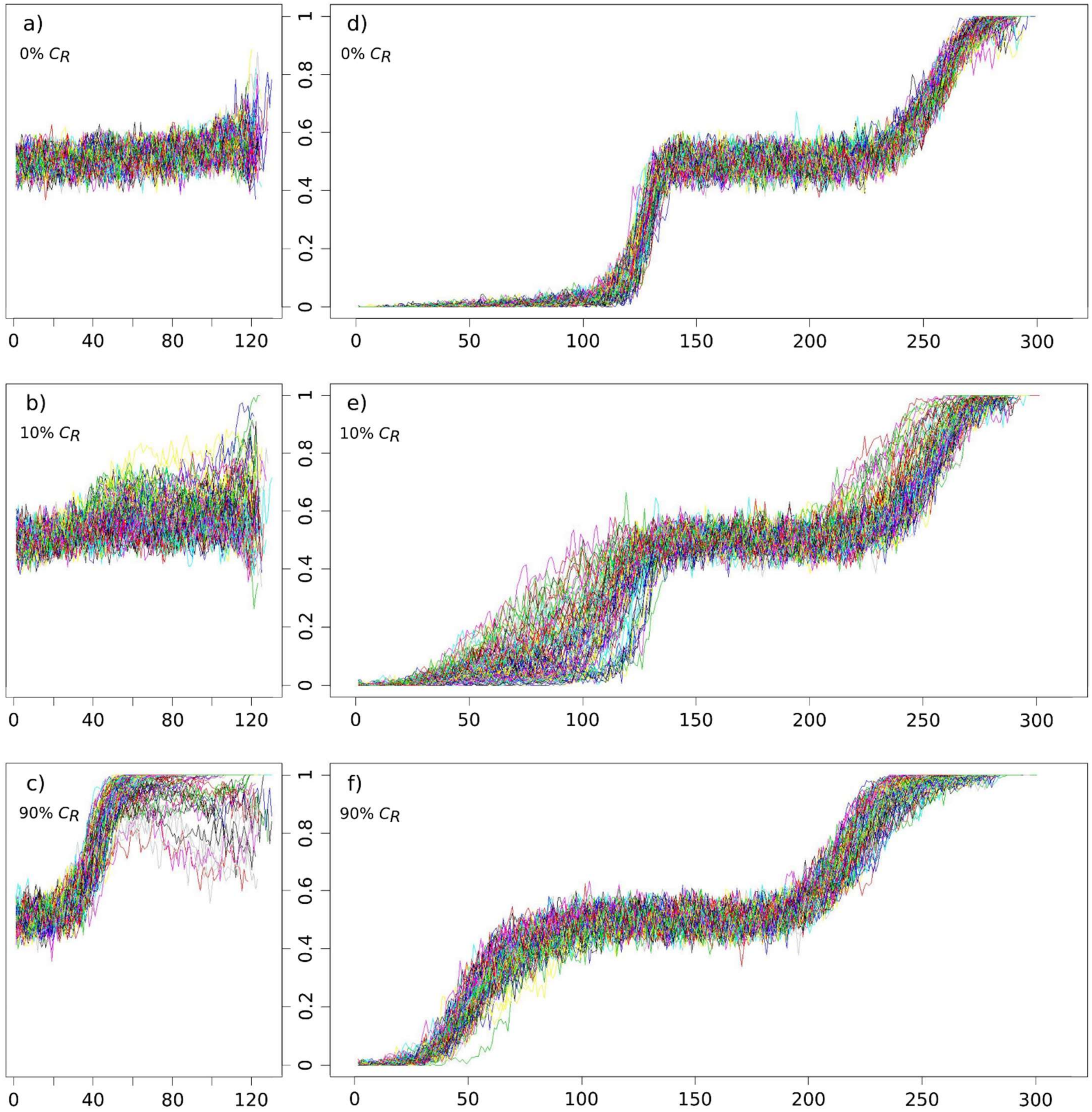


**Fig S10. Relative frequency of genotype *aa* among adults.** X axis: years of climate warming. Y axis: relative frequency. On the left: scenarios 0% $C_R$ (a), 10% $C_R$ (b) and 90% $C_R$ (c) starting with XX/XY system. On the right: scenarios 0% $C_R$ (d), 10% $C_R$ (e) and 90% $C_R$ (f) starting with ZW/ZZ system. Curves indicate separate simulations (100 repeated simulations per scenario).

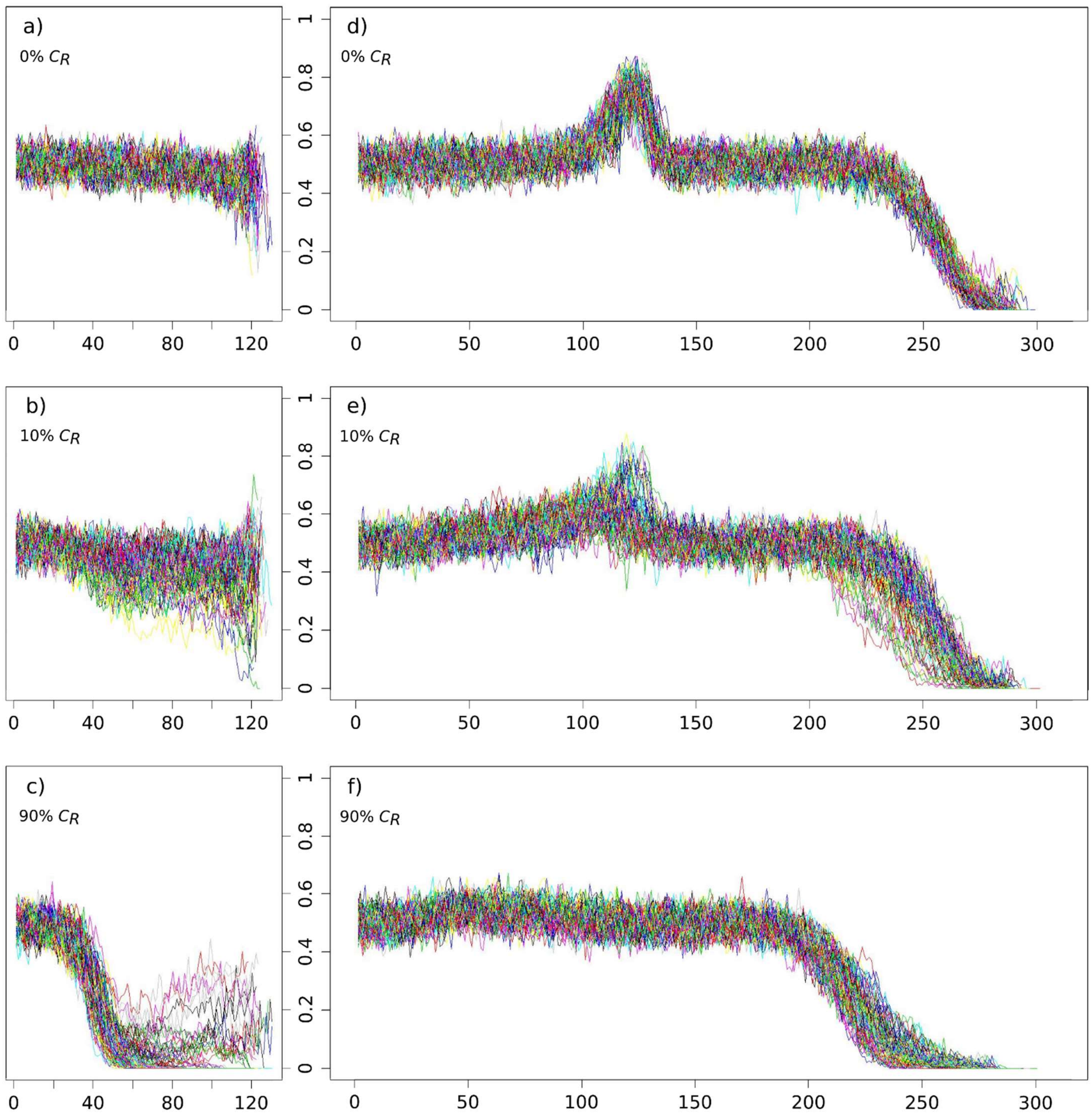


**Fig S11. Relative frequency of genotype *Aa* among adults.** X axis: years of climate warming. Y axis: relative frequency. On the left: scenarios 0% $C_R$ (a), 10% $C_R$ (b) and 90% $C_R$ (c) starting with XX/XY system. On the right: scenarios 0% $C_R$ (d), 10% $C_R$ (e) and 90% $C_R$ (f) starting with ZW/ZZ system. Curves indicate separate simulations (100 repeated simulations per scenario).

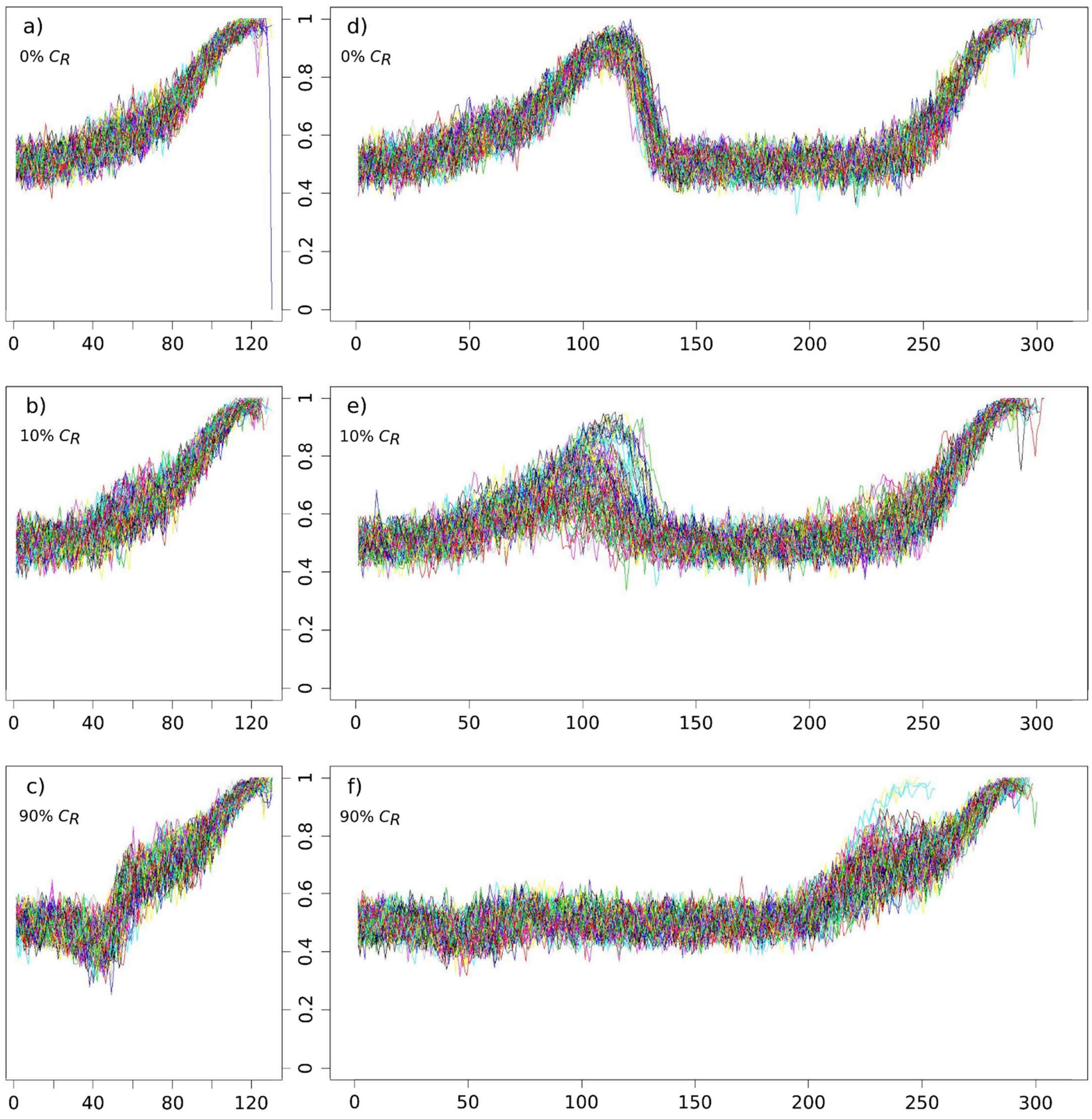


**Fig S12. Adult sex ratio.** X axis: years of climate warming. Y axis: proportion of males among adults. On the left: scenarios 0% $C_R$ (a), 10% $C_R$ (b) and 90% $C_R$ (c) starting with XX/XY system. On the right: scenarios 0% $C_R$ (d), 10% $C_R$ (e) and 90% $C_R$ (f) starting with ZW/ZZ system. Curves indicate separate simulations (100 repeated simulations per scenario).

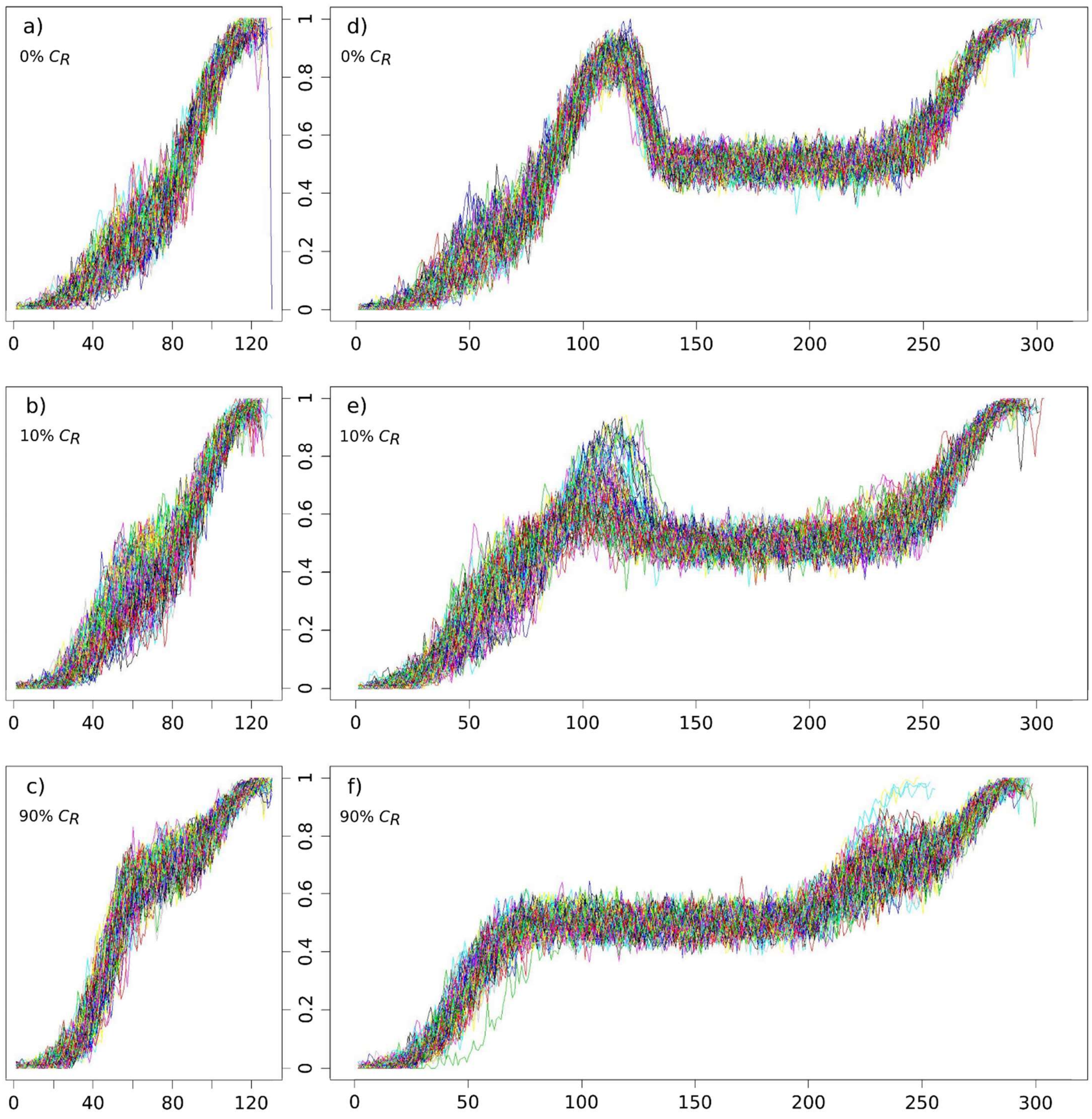


**Fig S13. Masculinization rate.** X axis: years of climate warming. Y axis: relative frequency of phenotypic males among genetically female individuals. On the left: scenarios 0% $C_R$ (a), 10% $C_R$ (b) and 90% $C_R$ (c) starting with XX/XY system (female genotypes: *Aa, aa*). On the right: scenarios 0% $C_R$ (d), 10% $C_R$ (e) and 90% $C_R$ (f) starting with ZW/ZZ system (female genotype: *aa*). Curves indicate separate simulations (100 repeated simulations per scenario).

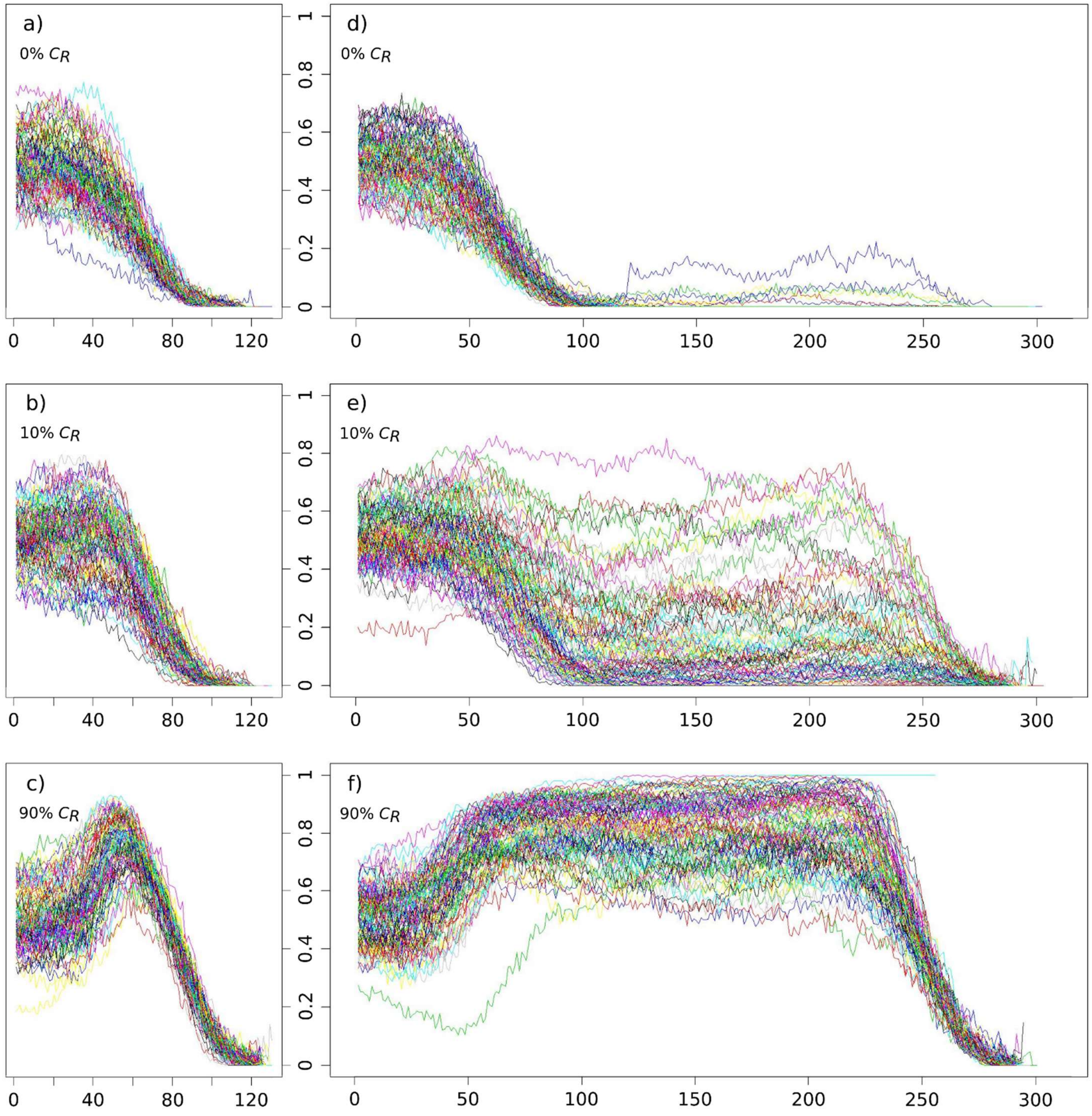


**Fig S14. Relative frequency of the *thr*$_{low}$ allele.** X axis: years of climate warming. Y axis: relative frequency. On the left: scenarios 0% $C_R$ (a), 10% $C_R$ (b) and 90% $C_R$ (c) starting with XX/XY system. On the right: scenarios 0% $C_R$ (d), 10% $C_R$ (e) and 90% $C_R$ (f) starting with ZW/ZZ system. Curves indicate separate simulations (100 repeated simulations per scenario).

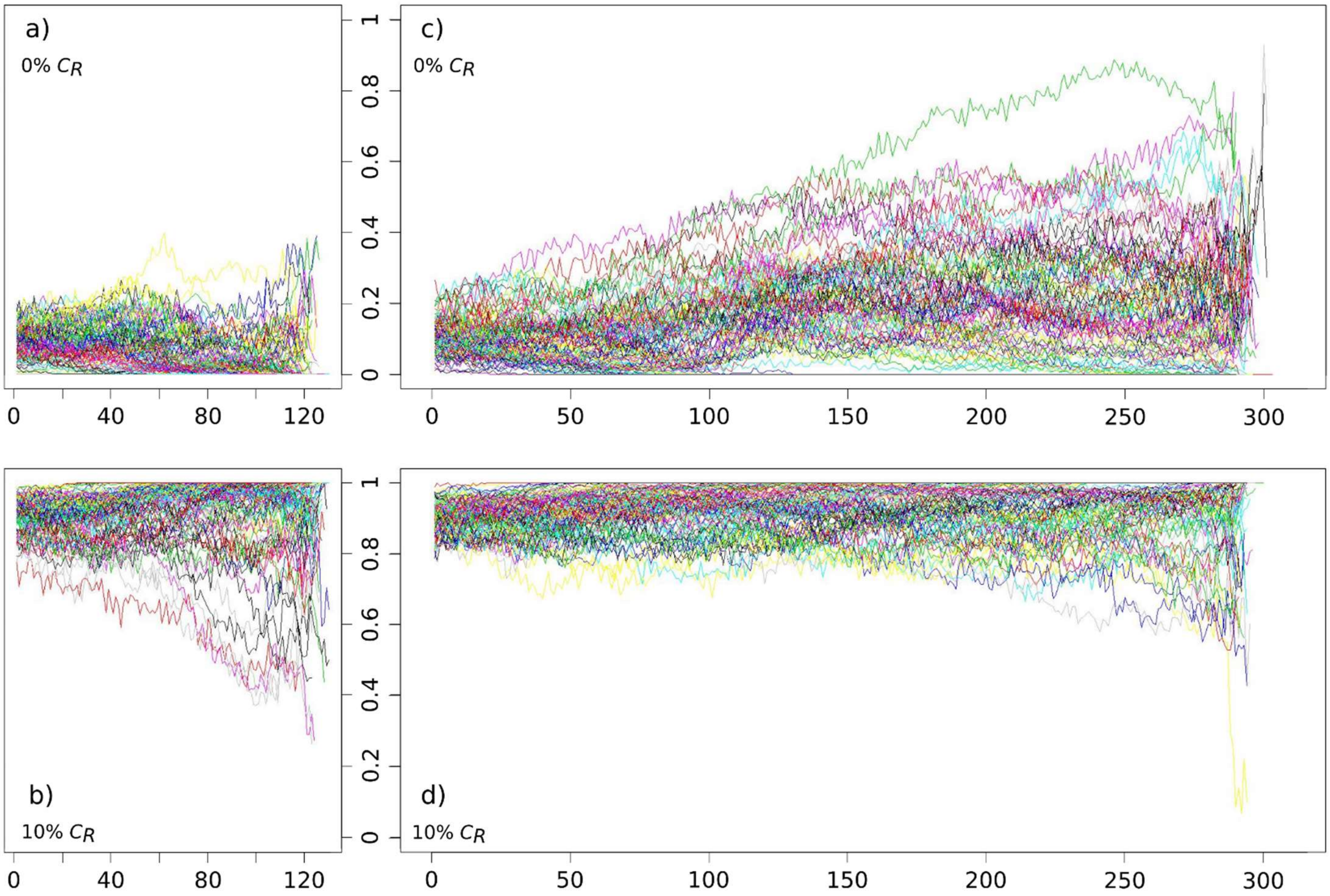


**Fig S15. Relative frequency of preference allele *$C_R$*.** X axis: years of climate warming. Y axis: relative frequency. On the left: scenarios 10% $C_R$ (a) and 90% $C_R$ (b) starting with XX/XY system. On the right: scenarios 10% $C_R$ (c) and 90% $C_R$ (d) starting with ZW/ZZ system. Curves indicate separate simulations (100 repeated simulations per scenario).

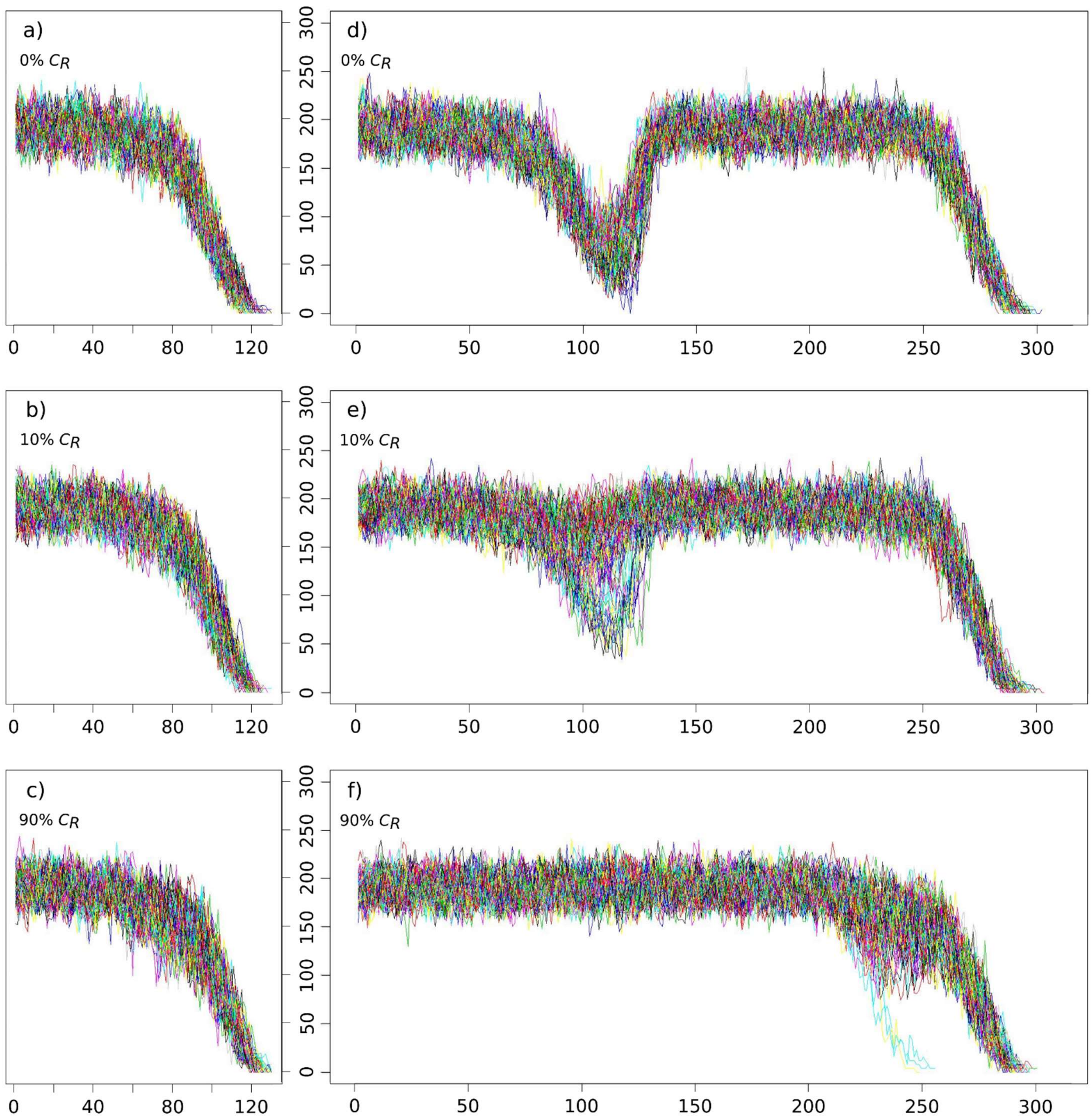


**Fig S16. Effective population size.** X axis: years of climate warming. Y axis: effective population size (calculated from the number of adult males and adult females present in the population). On the left: scenarios 0% $C_R$ (a), 10% $C_R$ (b) and 90% $C_R$ (c) starting with XX/XY system. On the right: scenarios 0% $C_R$ (d), 10% $C_R$ (e) and 90% $C_R$ (f) starting with ZZ/ZW system. Curves indicate separate simulations (100 repeated simulations per scenario).

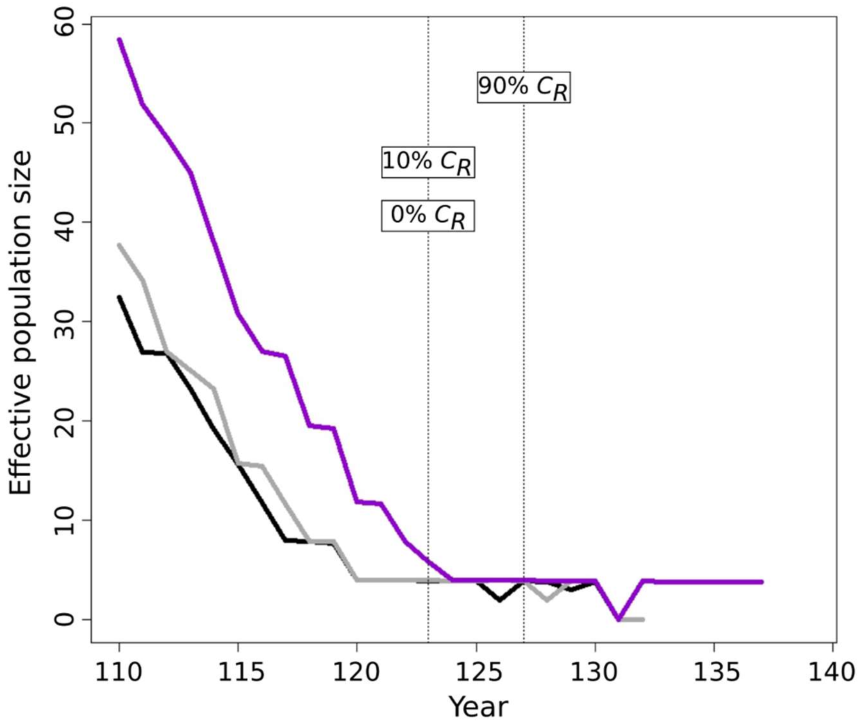


**Fig S17. Median effective population size during the last decades before extinction in originally XX/XY populations.** Years denote the number of years since the start of climate warming. Median effective population size calculated from the persisting populations out of a total of 100 in each year are shown for each scenario: 0% $C_R$ (black), 10% $C_R$ (grey) and 90% $C_R$ (prurple). Vertical dotted lines indicate the median extinction time for each scenario (note: this is the same in scenarios 0% $C_R$ and 10% $C_R$).
